# An activation-based high throughput screen identifies caspase-10 inhibitors

**DOI:** 10.1101/2024.12.15.625925

**Authors:** José O. Castellón, Constance Yuen, Brandon Han, Katrina H. Andrews, Samuel Ofori, Ashley R. Julio, Lisa M. Boatner, Maria F. Palafox, Nithesh Perumal, Robert Damoiseaux, Keriann M. Backus

**Affiliations:** Biological Chemistry Department, David Geffen School of Medicine, UCLA, Los Angeles, CA,90095, USA; Department of Chemistry and Biochemistry, UCLA, CA 90095, USA; California NanoSystems Institute (CNSI), UCLA, Los Angeles, CA, 90095, United States; Department of Molecular and Medical Pharmacology, UCLA, Los Angeles, CA, 90095, USA; Department of Human Genetics, David Geffen School of Medicine, UCLA, Los Angeles, CA,90095, USA; Department of Bioengineering, Samueli School of Engineering, UCLA, Los Angeles, CA, 90095, USA; Jonsson Comprehensive Cancer Center, UCLA, Los Angeles, CA 90095, USA; Eli and Edythe Broad Center of Regenerative Medicine and Stem Cell Research, UCLA, Los Angeles, CA 90095 USA; UCLA DOE Institute for Genomics and Proteomics, UCLA, Los Angeles, CA 90095, USA

## Abstract

Caspases are a family of highly homologous cysteine proteases that play critical roles in inflammation and apoptosis. Small molecule inhibitors are useful tools for studying caspase biology, complementary to genetic approaches. However, achieving inhibitor selectivity for individual members of this highly homologous enzyme family remains a major challenge in developing such tool compounds. Prior studies have revealed that one strategy to tackle this selectivity gap is to target the precursor or zymogen forms of individual caspases, which share reduced structural homology when compared to active proteases. To establish a screening assay that favors the discovery of zymogen-directed caspase-10 selective inhibitors, we engineered a low-background and high-activity *tobacco etch* virus (TEV)--activated caspase-10 protein. We then subjected this turn-on protease to a high-throughput screen of approximately 100,000 compounds, with an average Z’ value of 0.58 across all plates analyzed. Counter screening, including against TEV protease, delineated bona fide procaspase-10 inhibitors. Confirmatory studies identified a class of low-pH activatable caspase-10 inhibitors. In parallel, mode-of-action studies revealed that pifithrin-µ (**PFTµ**), a reported TP53 inhibitor, also functions as a promiscuous caspase inhibitor. Both inhibitor classes showed preferential zymogen inhibition. Given the generalized utility of activation assays, we expect our screening platform to have widespread applications in identifying state-specific protease inhibitors.

## Introduction

Human caspases are a family of 12 cysteine proteases, which are canonically associated with cell death^1–3^, but have also been tied to nearly all cellular processes, spanning activation^4, 5^, proliferation^6, 7^, differentiation^8, 9^, and cell migration^10^. Therefore, pharmacological manipulation of individual caspases represents an exciting opportunity to delineate caspase-specific functions and to intervene in human pathologies linked to dysregulated caspase activity, including metabolic^11^ and immune disorders^12–15^, cancers^16–20^, and neurodegenerative diseases^21^. Selective inhibitors are available for caspase-1^22^, caspase-2^23, 24^, caspase-6^25, 26^, and caspase-8^27, 28^. Despite these advances, obtaining highly selective inhibitors remains challenging, likely due in large part to the high sequence and structural homology of all caspases.

Caspase-10 is one such caspase that, while particularly intriguing from a target perspective due to its important functions in immune cell apoptosis, still lacks selective inhibitors. In fact, caspase-10 is one of the only caspases that is not labeled by many conventional peptide-based caspase inhibitors^29, 30^. Caspase-10 is absent in rodents^31^ and shares high sequence homology with caspase-8. Efforts to delineate caspase-10’s role in initiating apoptosis have been complicated by seemingly contradictory findings in immortalized cancer cells versus primary T cells^32, 33^. In cancer cells, caspase-8 rather than caspase-10 is the primary driver of extrinsic apoptosis^32–37^. Caspase-10 has even been implicated as a dominant negative regulator of programmed cell death in some cancer types^1, 38^. In contrast, in primary T cells, caspase-10 contributes to the initiation of apoptosis^13^; humans harboring inactivating mutations in caspase-10 experience autoimmunity and excessive T cell proliferation^39, 40^ because of decreased apoptosis. These human phenotypes indicate non-functionally redundant roles for caspase-8 and -10 and highlight the likely value of selective inhibitors targeting each caspase, both for further characterizing the unique and overlapping activities of each protease and, more broadly, towards the production of new chemical tool compounds to manipulate adaptive immune cell function.

Type II kinase inhibitors, which target the inactive form of enzymes, exemplify one strategy to improve inhibitor selectivity^41–44^. We took this approach in our prior work, which identified selective caspase-8 and caspase-2 inhibitors that function by targeting the zymogen, or uncleaved and inactive, precursor caspase proteoforms^24, 27, 45^. However, in our previous study, developing selective inhibitors for caspase-10 that did not cross-react with caspase-8 proved elusive despite our best efforts^27^. Consequently, there is an unmet need for new approaches to caspase-10 inhibitor discovery.

Here, we develop and apply a tobacco etch virus (TEV) activation-based screening platform to discover procaspase-10 inhibitors. To build this platform, we first generated an engineered caspase-10 protein (proCASP10TEV Linker) in which the caspase cleavage sites were replaced with TEV recognition sequences. This engineered protease showed low background, high stability, and robust TEV-dependent activity. After TEV activation, proCASP10TEV Linker protease showed comparable activity to recombinant active caspase-10. Enabled by this assay, we conducted a ∼100,000 compound screen, which had a hit rate of ∼0.22% (calculated as compounds with a z-score less than -3) and an average Z’-factor (a measure of how well positive and negative controls are separated)^46^ of 0.58 (**Table S1**). Our subsequent re-screening and counter-screening efforts validated hits and delineated TEV inhibitors from those targeting proCASP10. Resynthesis of a thiadiazine-containing hit compound (**SO265**) unexpectedly revealed that compound rearrangement was driving the observed caspase-10 inhibition. Additionally, mode-of-action studies revealed that the hit compound pifithrin-µ (**PFTµ**), a previously reported inhibitor of TP53^47^, is also a broadly reactive caspase inhibitor. Taken together, we expect that our hit compounds and innovative screening platform will advance efforts to discover potent and selective procaspase-10 inhibitors.

### Establishing a TEV-protease activatable caspase-10

Guided by previous reports of TEV- activatable caspases^24, 48, 49^, our first step was to engineer a TEV-activatable caspase-10 protein (**Figure 1A**). For our first attempt at engineering a TEV-cleavable construct (proCASP10TEV), we replaced the aspartate cleavage site, D415, with the TEV recognition site (**Figure S1**; D415ENLYFQG). While this protein did show substantial TEV-dependent increased activity, a high background in the absence of TEV protease was also observed (**Figure 1B**, **Figure S2**). We ascribed this background activity to the formation of activated protease in our purified protein sample (**Figure S3**). The presence of TEV-independent protease activity, and therefore, cleaved caspase, while somewhat varied between recombinant protein batches, was consistently observed in our purified protein after labeling with **Rho-DEVD-AOMK** (**Figure S3**). The background activity was also observed to increase markedly over time, even at nanomolar protein concentrations and with increasing concentration of the sodium citrate kosmotrope (**Figure S4**). Therefore, we deemed this construct incompatible with HTS, given the absolute requirement for a highly stable and consistently performing protein for large-scale screening applications.

**Figure 1.**
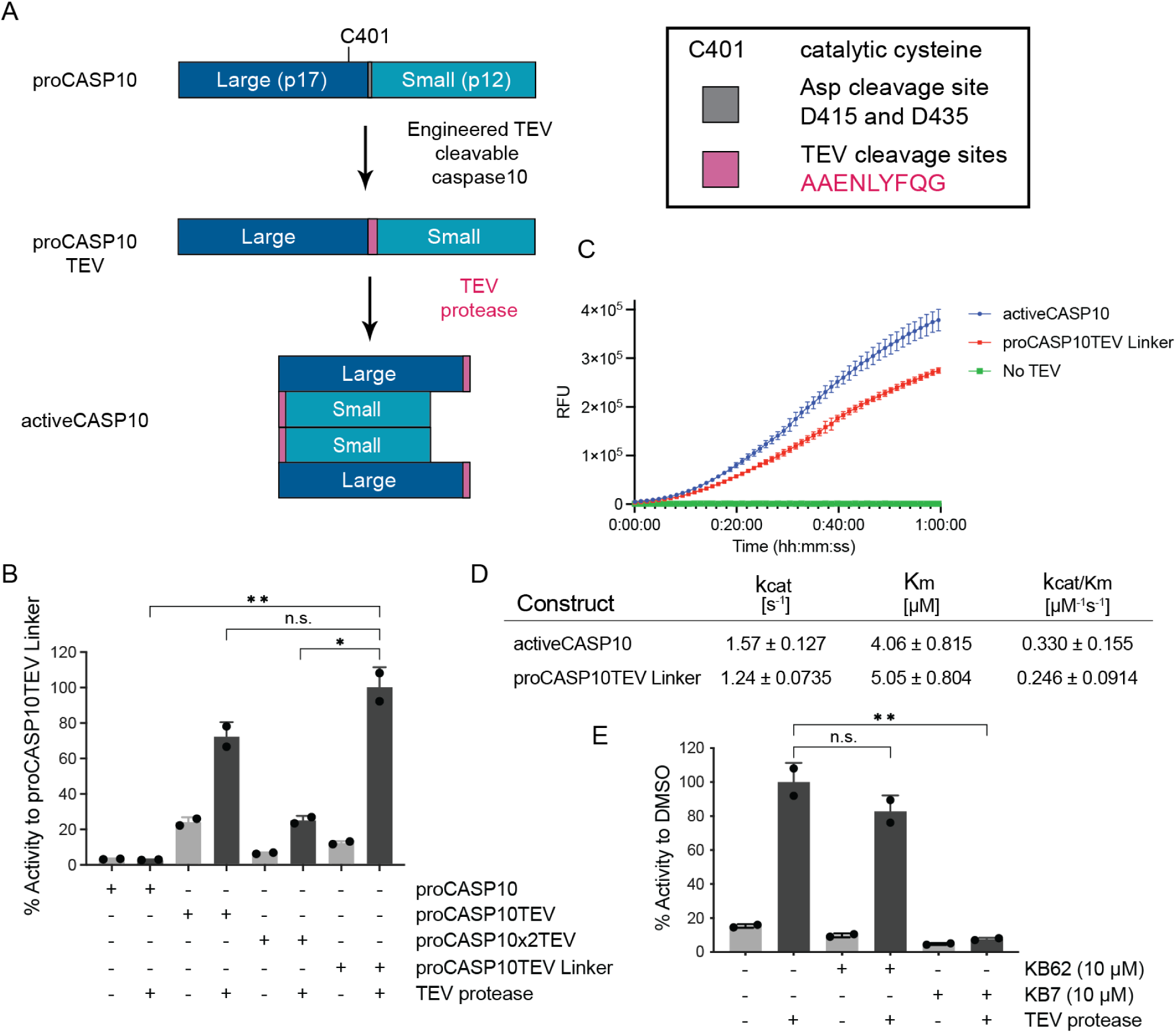
Robust TEV-cleavable functional caspase-10 is only cleaved in the presence of TEV protease and can be inhibited with a dual caspase-8/-10 inhibitor (KB7)^27^. (A) Design of engineered TEV cleavable caspase-10 containing TEV recognition site, ENLYFQG (pink). (B) Comparison of relative TEV-dependent and TEV-independent caspase activity for the indicated engineered proteins (333 nM) to cleave Ac-VDVAD-AFC fluorogenic substrate (10 µM) with/without TEV protease (667 nM). Sequences of engineered proteins can be found in Figure S1. (C) Time course comparing activeCASP10 (blue) versus proCASP10TEV Linker with (red) and without (green) TEV protease (667 nM) in the presence of Ac-VDVAD-AFC fluorogenic substrate (10 µM). (D) Michaelis-Menten kinetics comparing proCASP10TEV Linker (333 nM) activity to activeCASP10 (333 nM) activity as measured by cleavage of the fluorogenic substrate over time. (E) Assessing the relative inhibition of proCASP10 Linker protein by dual procaspase- 8/-10 inhibitor KB7 (10 µM) and negative control compound KB62 (10 µM)^27^. Assay conducted as in ‘B.’ For B and E, data represents mean value ± standard deviation for two biological replicates. For C and D, data represent mean values ± standard deviation for three biological replicates. Statistical significance was calculated with unpaired Student’s t-tests, * p<0.05, **p<0.001, ns, not significant p>0.05. Relative fluorescence units, RFU.

While caspase-10 is not known to harbor additional caspase cleavage sites, we postulated that D435 might also be recognized and cleaved, given the proximity to a likely caspase- recognition motif (PAED). Therefore, we inserted a second TEV motif to generate the proCASP2xTEV protein (**Figure S1**). Disappointingly, this enzyme showed low overall activity and negligible TEV-dependent activation, suggesting that sequence alterations at D435 are not well tolerated (**Figure 1B**).

While caspases show high selectivity for aspartyl residues, we postulated that the glutamate residue in the TEV recognition sequence could be recognized and cleaved at a low level, thereby rationalizing the observed high TEV-independent background for the proCASP10TEV protein. To test this hypothesis, we next generated a caspase-10 construct (proCASP10TEV Linker) in which a two-alanine spacer was included to reposition this glutamate further from the remainder of the caspase recognition motif. We additionally optimized our expression and purification protocol to reduce self-activation, both by decreasing the induction time and by not freezing the cell pellets prior to purification (see methods). Pleasingly, the proCASP10TEV Linker protein exhibited both higher overall activity and reduced background compared to our initial construct (**Figure 1B**). Further supporting the improved behavior of this protein, we detected no formation of cleaved caspase (∼20 kDa) by gel-based analysis (**Figure S5**).

### Assessing the suitability of proCASP10TEV Linker for high throughput screening (HTS)

To test whether the proCASP10TEV Linker protein would faithfully recapitulate proCASP10 activity and thus prove suitable for HTS, we further assessed the activity of this protein. Gratifyingly, upon the addition of TEV protease, we observed marked TEV-dependent increased catalytic activity (**Figure 1B, C**). TEV activation occurred rapidly and required only a modest (∼500 nM) concentration of TEV protease to achieve complete conversion of proCASP10TEV Linker (333 nM) to the cleaved proteoform (**Figure S6** and **Figure S7**). The proCASP10TEV Linker protein, once activated by TEV protease, showed near-comparable activity to activeCASP10 (**Figure 1C**), with only a modest decrease in kcat and Km (**Figure 1D**). We find that the protein labels robustly with our previously reported caspase-8/10 click probe **KB61** (**Figure S6**)^27^, which corroborates that the proCASP10TEV Linker protein behaves similarly to proCASP10. The TEV cleavable construct was also inhibited by the dual caspase-8 and caspase-10 inhibitor **KB7** (10 µM) and was not inhibited by the structurally matched inactive control, **KB62** (10 µM) (**Figure 1E**)^27^. These findings provided further evidence that our engineered protein behaves similarly to proCASP10 and, additionally, demonstrated that the protein is well suited to assess procaspase inhibition.

We next vetted our assay conditions for HTS compatibility, with the goal of ensuring both high activity and high stability over longer assay periods. Additionally, we found that the proCASP10TEV Linker protein showed increased activity in the presence of the kosmotrope sodium citrate at increasing concentrations (37 mM, 111 mM, and 333 mM) (**Figure S8**). This added activity was restricted to the TEV-cleaved/activated protein; little TEV-independent activity was observed after chaotrope addition in the absence of TEV protease. Providing evidence of high enzyme stability, a favorable property for screening applications, proCASP10TEV Linker showed no substantial change in enzyme activity after prolonged (18h) incubations times for both 4 °C and ambient temperature (**Figure S9**). Additionally, substrate turnover was observed to progress in a linear manner for the initial ∼2h period, with increasing activity observed up to 6h (**Figure S10**), which provides a wide time window for acquiring data. Thus, with high stability, low background, and high activity enzyme in hand, we turned to HTS implementation.

### A small-scale screen of two pharmacologically active library compounds confirms assay compatibility with HTS

Toward large-scale library screening, our next step was to validate our TEV-mediated caspase activation assay and establish a miniaturized (384-well plate) semi-automated workflow. Following the workflow shown in **Figure 2A**, we first screened the ‘Library of Pharmacologically Active Compounds’ (LOPAC@1280), which is a widely utilized library of bioactive compounds that contain known promiscuous protease inhibitors such as E-64, along with a second library consisting of structurally diverse FDA approved drugs covering a broad spectrum of therapeutic areas^50^. Key features of our screen include (1) the pre-incubation of compound library members with the proCASP10TEV Linker construct to favor the detection of procaspase inhibitors and (2) the automated dispensing of the premixed TEV protease and caspase substrate solution followed by automated plate reading. These two automation steps were designed to ensure the consistent timing of our assay to minimize both plate-to-plate and library-to-library variability. Confirming the robustness of our approach, this initial screen had a Z’-factor above 0.5 across 8 screened plates, with a total of 30 out of 2569 compounds showing less than a 50% decrease in caspase activity relative to the DMSO control (**Figure S11**). Illustrating the robust performance of our screen, the Z’-factor comparing our positive control KB7 inhibitor-treated wells to DMSO-treated wells^27^ was 0.90 (**Figure S12**)—values above 0.5 are considered an acceptable range for HTS Z’-factor^46^.

**Figure 2.**
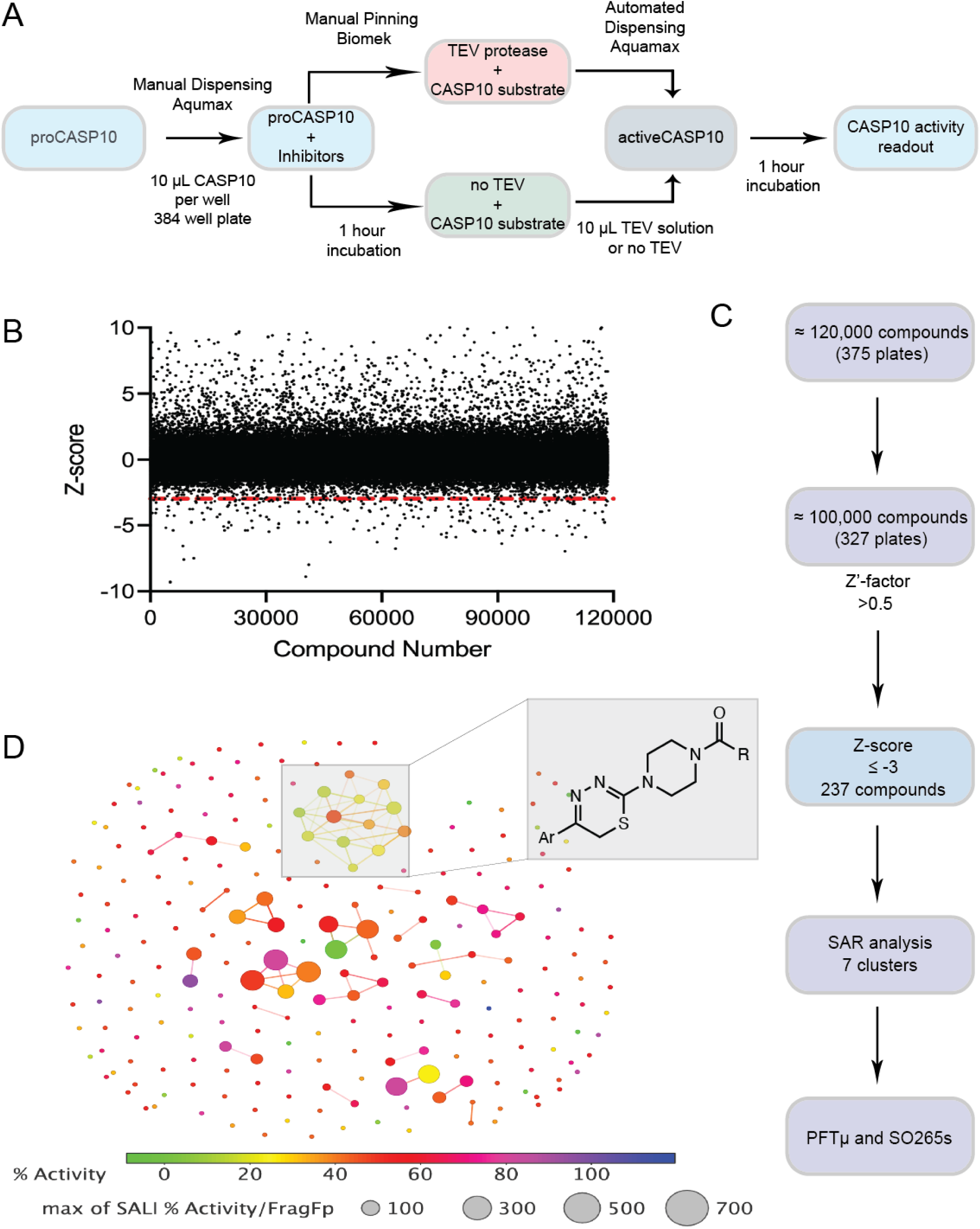
TEV cleavable proCASP10TEV Linker enables a >100,000 compound screen. (A) Scheme of HTS setup with proCASP10TEV Linker (333 nM concentration) incubated with screening compounds (10 µM) for 1h in a 384-well format followed by the addition of the fluorogenic substrate (Ac-VDAVAD-AFC, 10 µM). Endpoint reads were measured and collected after 1h incubation with the TEV substrate solution. (B) z-scores of >100,000 compounds (arbitrarily numbered) screened against proCASP10TEV Linker. The red line indicates hits below a z-score of -3. (C) Summary of filtering parameters for HTS screen. The first filtering step was to remove plates that were not within the desired range of Z’-values (0.5 - 1.0), followed by a second filtering step that identified a total of 237 compounds with a z-score of ≤ -3 (D) SAR analysis of 237 hit compounds identified from HTS using the structure-activity landscape index (SALI) analysis as described in the DataWarrior SAR analysis methods section. The structure of recurring hit chemotypes identified by SAR cluster analysis (DataWarrior, v06.01.04.)^51^ are shown for cluster 1. All screening data is in **Table S1.**

### Large-scale screen identifies hit caspase inhibitors

Guided by the successful implementation of our pilot screen, we next deployed our platform to screen 118,498 total compounds (all at 10 μM). In aggregate, this screen had Z’ values ranging from 0.4 - 1.0 and a hit rate of 0.81% for a total of 963 compounds, affording a 50% decrease in proCASP10TEV Linker activity (**Figure S13**), with 237 unique hits showing a ≤ -3 (**Figure 2B**). The screen filtering steps are summarized in **Figure 2C**. Re-screen validation of the 237 compounds results in 38 compounds with < 50% activity (**Figure S14A**). z-score analysis of the follow-up screen resulted in 97 (or approximately 41%) compounds below 3 standard deviations away from the mean of DMSO control samples (**Figure S14B**). Structural similarity clustering analysis using Datawarrior^51^ revealed that approximately 22% of the 237 hits (defined as z-score ≤ -3 or those values that are three standard deviations lower than the mean of the DMSO controls calculated per plate) clustered together, providing evidence of bona fide inhibitors. The 7 major hit clusters consisting of over 50 compounds are shown in **Figure 2D**. One prominent and intriguing chemotype was a class of thiadiazine-containing compounds identified in cluster 1 (**Figure 2D**). Upon closer inspection of the hits, several compounds with clear electrophilic groups stood out, including several organometallic species such as cisplatin (**Table S1**), several cyanoacryl sulfones, such as **HTS- 2** and the related **HTS-6,** which is a previously reported as non-specific κB kinase (IKK) inhibitor^52^, and pifithrin-µ (**PFTµ**), a previously reported TP53 inhibitor^47^ (**Scheme S1**). As caspase-10 is a cysteine protease, we anticipated that some of these electrophilic compounds might react with the catalytic nucleophile.

### Hit prioritization by re-screening and counter-screening against activeCASP10

To further interrogate our hits and confirm on-target activity against procaspase-10, our next step was determining if our hits were selective for proCASP10 over the active proteoform. Using our previously reported fluorogenic caspase assay^24^, we measured the relative inhibition of recombinant active CASP10 by each of our 237 prioritized hits. This screen had a Z’ value of 0.73, resulting in 9 of 237 compounds having ≤ 50% activeCASP10 activity (Figure **S15A** and **Table S1**), with 78 compounds being 3 standard deviations below the DMSO control (**Figure S15B**). Notably, several of our aforementioned hits, including **HTS-6**, **SO265**, and **PFTµ**, showed substantially reduced inhibition of activeCASP10 compared to the proCASP10TEV Linker (**Figure S14A and S15A**). SAR clustering analysis of the counter screen showed ∼20 different clustering types for activeCASP10, but overall, less reactivity towards active compared to the pro-form (**Figure S16**), consistent with more favorable inhibition of the proenzyme. Together, our prioritized compounds (**Scheme S1**) included 10 total inhibitors that showed preferential activity against proCASP10TEV when compared to activeCASP10.

### TEV assay identifies likely TEV inhibitors

As our proCASP10TEV Linker screen requires the addition of TEV protease for caspase activation, we anticipated that some hits might be bona fide TEV protease inhibitors rather than caspase inhibitors. To test this hypothesis and to filter out such compounds, we established a TEV protease activity assay using a customized fluorogenic substrate (see methods) inspired by prior TEV protease assays^53^. Following the workflow shown in **Figure 3A**, we first optimized both the substrate concentration and time points to ensure activity was within the enzyme’s linear range (**Figure S17**). We find that TEV protease was highly active towards our fluorogenic substrate, with complete consumption of substrate within five minutes (**Figure 3B**). Using automated dispensing to ensure the rapid and equal delivery of substrate across assay conditions, we observed comparable Michaelis-Menten kinetic parameters to those reported previously for TEV protease^54^ (**Figure 3C**).

**Figure 3.**
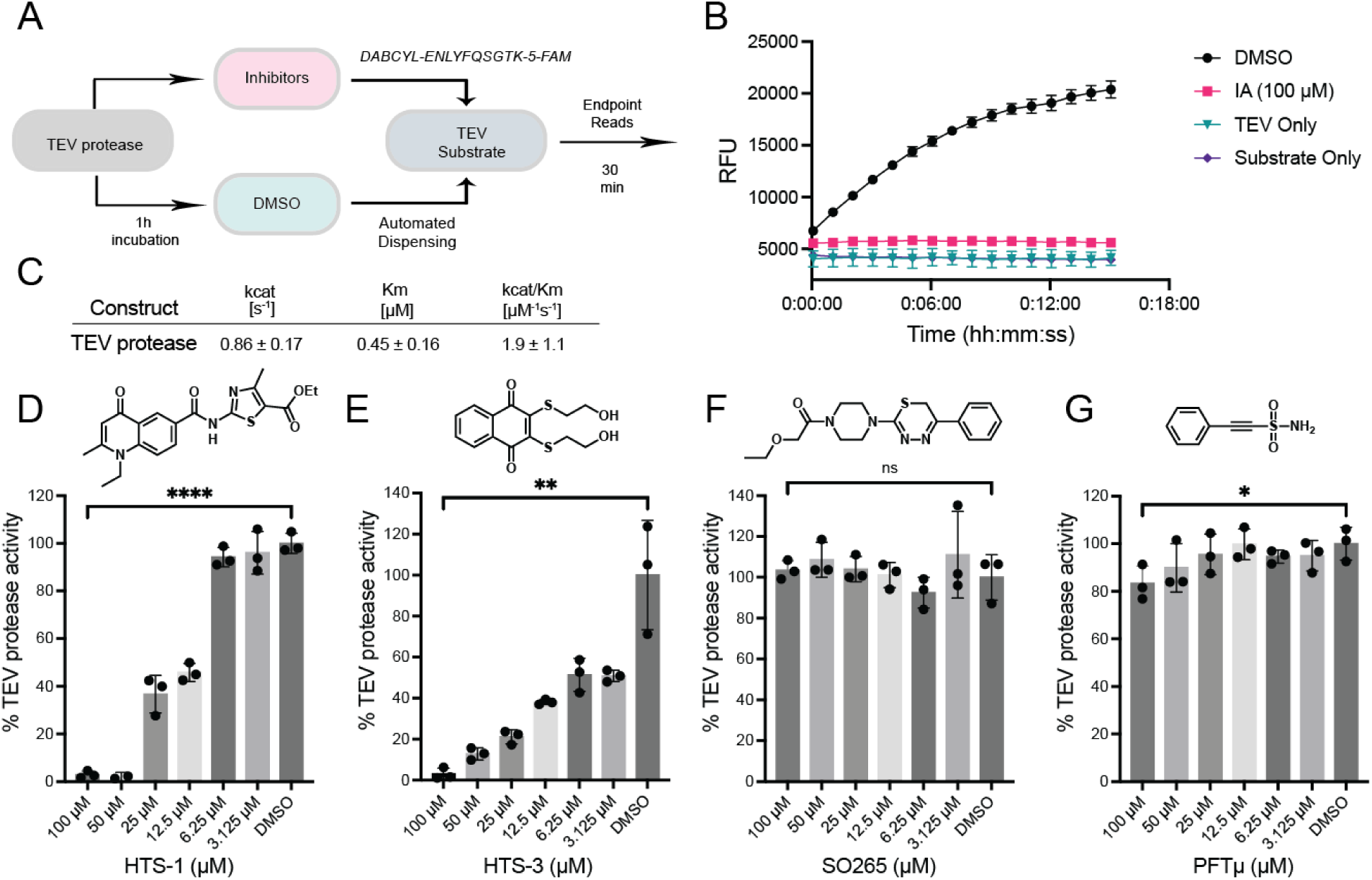
Procaspase-10 HTS identified TEV protease inhibitors. (A) General scheme of TEV protease activity assay with our in-house TEV cleavable fluorogenic substrate, **DABCYL-ENLYFQSGTK-5-FAM**. (B) Shows relative fluorescent units (RFUs) for time- dependent increased fluorescence (λex = 495 nm λem = 550 nm) for TEV protease (100 nM) exposed to either vehicle (DMSO) or iodoacetamide (IA). (C) Michaelis-Menten kinetics of TEV protease (667 nM) with TEV substrate. (D-G) TEV protease (50 nM) activity assay treated with the indicated screening compounds at the indicated concentrations for 1h in PBS buffer under ambient conditions. For D-G, data represent mean values ± standard deviation for three biological replicates. Statistical significance was calculated with unpaired Student’s t-tests, * p<0.05, **p<0.01, ****p<0.0001, ns, not significant p>0.05.

Using this optimized assay, we used freshly sourced compound stocks to evaluate the TEV protease inhibitory activity of ten prioritized hits (**Figure 3D-G**, **Figure S18,** and **Figure S19**). Notably, while most compounds were sourced commercially, compound **SO265** required in-house synthesis (**Scheme S2** for the synthetic route for **SO265**). We observed substantial dose-dependent inhibition of TEV protease for **HTS-1 (Figure 3D), HTS-2 (Figure S18A),** and **HTS-3** (**Figure 3E**). Some inhibition was also observed for **HTS-6** (**Figure S18B**), **HTS-4** (**Figure 18C**), and **HTS-5** (**Figure S18D**). Therefore, we excluded these likely TEV inhibitors from further analysis. Encouragingly, several HTS hits stood out as having no appreciable TEV protease inhibitory activity (**Figure 3F, Figure S19A,** and **Figure S19B)**. Notably, **PFTµ** afforded a slight, but not significant, decrease in TEV protease activity at the highest tested concentration (100 µM) (**Figure 3G**).

### Caspase-10 inhibition is confirmed by activity assays

We next turned to confirm that our screening hits engage both recombinant and endogenous proCASP10 protein. We first rescreened a subset of the prioritized hits in our proCASP10TEV Linker assay. While we observed that most compounds only showed modest caspase-10 inhibition (**Figure S20**), several compounds stood out, including **HTS-6** (**Figure S21**), the TEV protease inhibitor **HTS-2** (**Figure S22**), and **PFTµ** (**Figure S23**). We then turned to gel-based activity-based protein profiling (ABPP) analysis to further corroborate our inhibition data. Having previously established ABPP gel-based assays for procaspase-8 and -10 using the **KB61** click probe^27^ in HEK293T lysates, we first deployed this ABPP assay to assess compound engagement of recombinant procaspase-10 at the catalytic cysteine residue C401 (**Figure S24**), focusing on the compounds that contained obvious electrophilic moieties, namely **HTS-6** and **HTS-2** and **PFTµ**. We find that **PFTµ** shows similar potency when compared to established caspase-8/10 dual inhibitor **KB7**^27^. Both **HTS-6** and **HTS-2** exhibited high proteome-wide reactivity, indicating that their caspase engagement is likely driven by the high electrophilicity of the cyanoacryl sulfone moiety. Further illustrating the increased reactivity of these two compounds relative to **PFTµ**, we also observed increased competition of iodoacetamide-rhodamine (IA-Rho) (**Figure S25**), consistent with generalized cysteine reactivity rather than a caspase-10-specific effect. Due to their high reactivity, we excluded **HTS-6** and **HTS-2**, from further analyses.

### Compound rearrangement drives caspase inhibition for SO265, which shows preferential inhibition of the pro-form of caspase-10

As the **SO265** thiadiazine chemotype had shown pronounced procaspase-10 inhibition in our initial screen (approximately 80% proCASP10TEV Linker inhibition; **Figure 2D** and **Figure S14**), we were surprised by the lack of activity using our resynthesized compound (20% proCASP10TEV Linker inhibition) (**Figures S26**). Substituted thiadiazine rings can isomerize to thiol imidazole moieties, gaining aromaticity^55^. Therefore, we hypothesized that such a rearrangement of **SO265**, which could be favored by lower pH, could be driving its inhibitory activity (**Figure 4A**). To test this hypothesis, we subjected **SO265** to acidic conditions (10 mM HCl) prior to assessing caspase inhibitory activity. Consistent with our hypothesis, we observed increased inhibitory activity for stocks of **SO265** pretreated with acid (**Figure S26**). This activity was generally restricted to caspase-10, with a more modest increase in inhibition observed for TEV protease (**Figure S27**).

**Figure 4.**
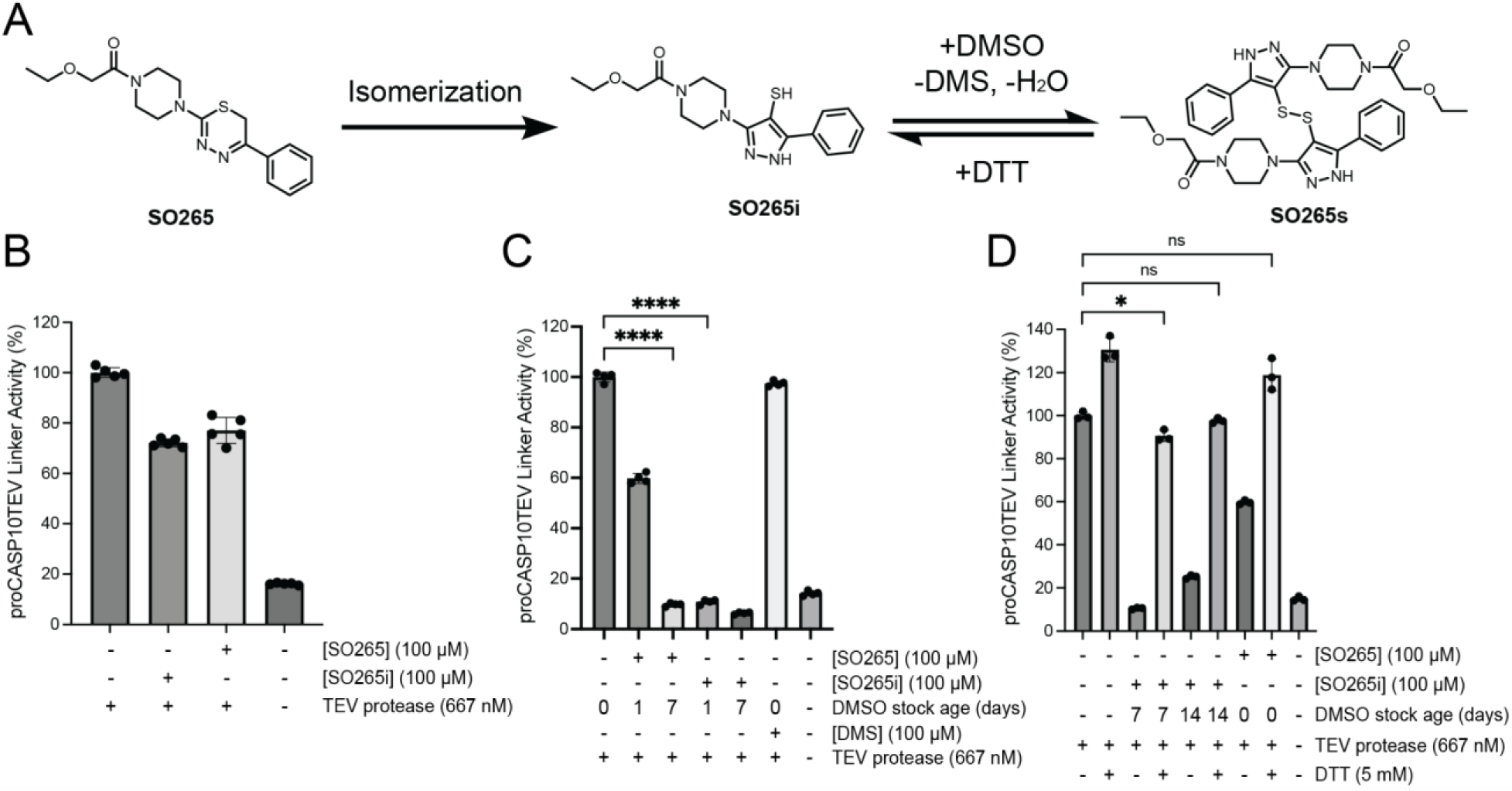
**The rearranged product of SO265 inhibits proCASP10TEV Linker activity**. (A) Proposed scheme of SO265 isomerization and formation of the disulfide product (B) Relative activity of proCASP10 Linker protein (333 nM) treated with the indicated compounds for 1h followed by addition of TEV protease (667 nM) and fluorogenic substrate (Ac-VDAVAD-AFC, 10 µM, and 333 mM sodium citrate) in PBS buffer. (C) The indicated compound stocks were subjected to either one- or seven-day incubation in DMSO at ambient conditions. Subsequently, compound-treated proCASP10 Linker protein was assayed as in ‘B’ with 1h pre-exposure to either dimethyl sulfide (DMS) or the indicated compounds. (D) Compounds incubated for the indicated days in DMSO were further subjected to DTT (5 mM DTT) followed by evaluation for caspase inhibition as described in ‘B.’ For B-D, data represent mean values ± standard deviation for four biological replicates. Statistical significance was calculated with unpaired Student’s t-tests, * p<0.05, ****p<0.0001, ns, not significant p>0.05.

Guided by this increased activity, we next tested the putative isomer, **SO265i** (**Figure 4A**), which we had isolated from the reaction mixture during the synthesis of **SO265.** While the masses of **SO265** and **SO265i** are identical, the two compounds can be distinguished via H-NMR analysis **(Figure S28A**) and LC-MS analysis (**Figure S28B-F**). Notably, the thiadiazine methylene signal was only present in **SO265**, with loss in the **SO265** spectra indicating that aromatization had indeed occurred. The **SO265i** spectra uniquely featured a signal for a thiol proton, consistent with the proposed isomerization. The presence of a thiol in **SO265i** was further corroborated by IR analysis (**Figure S29**). However, when tested in our activity assay, **SO265i** exhibited equivalent caspase-10 inhibition (30% inhibition) to **SO265**, indicating that the thiol imidazole was not the active species (**Figure 4B**).

Following a 2-day incubation at room temperature in DMSO, repeated NMR analysis revealed that the **SO265i** stock had converted into a second species. This species was distinguished by the loss of the thiol proton and an increase in a signal matching the expected shift of dimethylsulfide (DMS) (**Figure S28A**). We hypothesized that DMSO-mediated oxidation of the **SO265i** thiol to a disulfide (**SO265s**) was occurring; this is a reported redox reaction that produces DMS as a byproduct and is promoted by low thiol pKa values^56, 57^. Therefore, to further assess whether **SO265s** was the active species responsible for caspase-10 inhibition, we incubated 50 mM DMSO stocks of **SO265** and **SO265i** at room temperature for one, seven, and 14 days to promote the formation of **SO265s.** We then tested the activity of each of these compound stocks alongside DMS as a control (**Figure 4C**). Consistent with **SO265s** as the active species, we observed a time-dependent increase in inhibition of proCASP10TEV Linker for both **SO265** and **SO265i**; DMS had no effects on proCASP10TEV Linker activity. Providing further evidence of the likelihood that the rearranged disulfide structure is the active compound, the addition of 5 mM DTT to the compound stock completely abolished inhibitory activity (**Figure 4D**). **SO265s** also showed increased reactivity with glutathione when compared to **SO265** and **SO265i** (**Figure S30**). Taken together, these data provide evidence that **SO265s** is a cysteine-reactive compound that blocks proCASP10TEV Linker activity with preferential inhibition of the zymogen (**Figure S31).** More broadly, we find that **SO265s** show some caspase selectivity, also inhibiting active caspase-8 (**Figure S32**), whereas only a slight decrease in activity was observed for active caspase-3 (**Figure S33**) and caspase-9 (**Figure S34**).

### CETSA confirms caspase-10 labeling

As all our data, thus far, had been assessed for recombinant protein, we next opted to extend our analyses to assess the compound engagement of endogenous procaspase-8 and -10. For this, we turned to Cellular Thermal Shift Assay (CETSA)^58^ to measure binding-induced changes to protein thermal stability. To validate our assay, we subjected Jurkat lysates spiked with recombinant hexahistidine-tagged proCASP10 to CETSA analysis, comparing the thermal stability of the proCASP10 construct to that of the inactive catalytic cysteine mutant construct (proCASP10 C401A) with and without addition of **KB7**, as a positive control. We observe a marked decrease in thermal stability upon **KB7** treatment for both the endogenous (anti-caspase-10 signal) and recombinant (anti-His signal) caspase-10 proteins (**Figure 5A** and complete blots in **Figure S35**). The C401A mutant protein did not show a similar thermal shift (**Figure S36**), which is consistent with the covalent modification of the C401, which is the catalytic nucleophile, by **KB7**. Extension of this analysis revealed that, like **KB7**, **PFTµ** induced destabilization of the wildtype spiked proCASP10 and endogenous proCASP10 (**Figure 5B** and complete blots in **Figure S37**) but not the C401A mutant protein (**Figure 5C** and complete blots in **Figure S38**), which further confirms that **PFTµ** labels C401 Unexpectedly, the disulfide product (**SO265s)** caused some protein stabilization, suggesting an alternate mode of action compared to the two active site-directed inhibitors (**Figure 5D** and complete blots in **Figure S39**). Corroborating an alternate mode of engagement by **SO265s**, we observed no competition in gel-based ABPP analysis using the **KB61** click probe against procaspase-10 (**Figure S40A**), unlike **PFTµ** that labeled both procaspase-10 (**Figure S40A**) and procaspase-8 (**Figure S40B**). Disappointingly, we observed no similar compound-induced thermal stability shift for endogenous procaspase-8 upon **KB7** treatment (**Figure S35**) nor **PFTµ** treatment (**Figure S37**), indicating that the CETSA assay was not suitable for evaluation of caspase-8 target engagement. Therefore, we returned to enzyme activity assays for broader target analysis.

**Figure 5.**
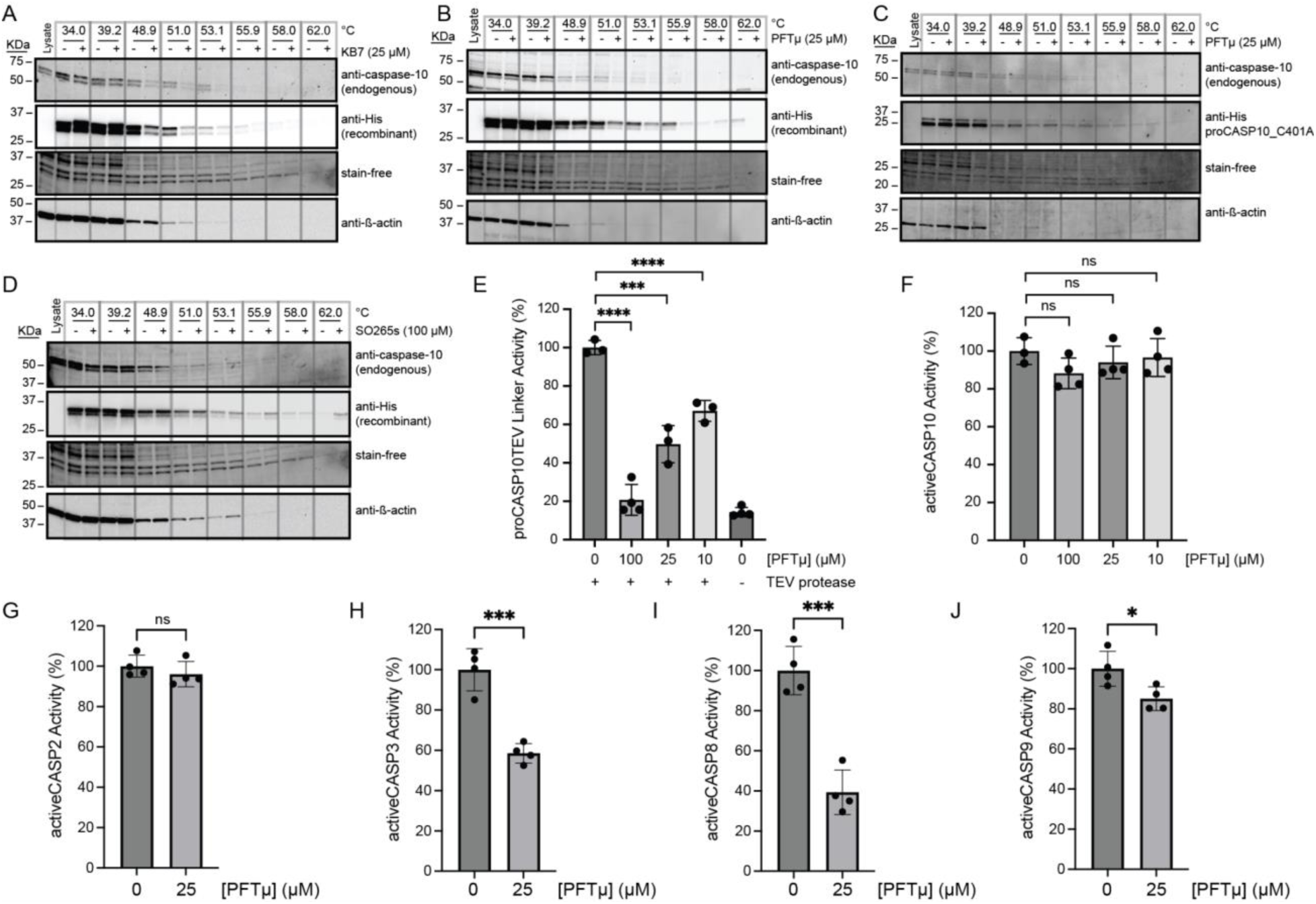
**PFTµ and SO265s impact the stability of both recombinant and endogenous procaspase-10**. (A-D) Cellular thermal shift assay (CETSA)^58^ analysis blot of Jurkat cell lysates spiked with the indicated recombinant caspase-10 proteins featuring C-terminal hexahistidine tag and treated with the indicated compounds at the indicated concentrations or vehicle (DMSO) and subjected to immunoblot analysis for using both anti-His and anti-caspase-10 antibodies to visualize recombinant and endogenous protein, respectively. Loading was visualized both with the BioRad Chemidoc Stain-Free imaging technology^59^ and with anti-β-actin. For ‘A, B, and D,’ recombinant proCASP10 and for ‘C,’ recombinant proCASP10_C401A were analyzed. (E) Relative activity of proCASP10TEV Linker protein (333 nM concentration in PBS buffer) exposed to **PFTµ** at the indicated concentrations for 1h followed by TEV protease (667 nM) and analysis with Ac-VDVAD-AFC fluorogenic substrate (10 µM) in PBS supplemented with 333 mM sodium citrate. (F-J) Relative activity of recombinant active caspases in PBS buffer analyzed with Ac-VDVAD-AFC fluorogenic substrate (10 µM) after treatment with the indicated concentrations of **PFTµ** for 1h. For activeCASP10, fluorogenic substrate (10 µM) in PBS was supplemented with 333 mM citrate. Recombinant proteins used in ‘F’ activeCASP10 (333 nM), ‘G’ activeCASP-2 (0.3 µM), ‘H’ activeCASP-3 (0.3 µM), and ‘I’ activeCASP9 (1 µM). For A–D, data is representative of two biological replicates. For E–J, data represent mean values ± standard deviation for four biological replicates. Statistical significance was calculated with unpaired Student’s t-tests, * p<0.05, ***p<0.001, ****p<0.0001, ns, not significant p>0.05.

### Pifithrin-µ is a promiscuous caspase inhibitor that prevents FasL-mediated apoptosis

In contrast with the complex activation mechanism of **SO265s**, which likely could complicate assessing in-cell activity, we expected **PFTµ** to retain caspase inhibitory activity in complex cell environments. This expectation is further supported by **PFTµ**’s well-documented anti-apoptotic activity, which has been ascribed to its function as a TP53 inhibitor^47^. Therefore, we prioritized **PFTµ** for further analysis. Activity assay analysis revealed that **PFTµ** showed some preferential inhibition across a panel of analyzed caspases; consistent with our screening data, we observed preferential inhibition of procaspase-10 when compared to active caspase-10 (**Figure 5E, F** and **Figure S14, S15**). Similarly, **PFTµ** also preferentially labeled procaspase-2, as analyzed using our caspase-2 directed click probe^24^ and ABPP analysis (**Figure S41**). No detectable inhibition of active caspase-2 was observed, either by ABPP gel (**Figure S41**) or activity assay (**Figure 5G**). **PFTµ** also significantly inhibited active caspase-3 and active caspase-8, with slight, albeit significant, inhibition observed for active caspase-9 (**Figure 5H-J**). These data provided evidence that **PFTµ** is a promiscuous caspase inhibitor.

We next turned to chemoproteomics to more broadly assess whether PFTµ labels endogenous caspases. We deployed our established cysteine chemoproteomic platform^60^ in which covalent labeling sites are identified in a competitive manner using the pan cysteine-reactive probe iodoacetamide alkyne (**IAA**) and isotopically enriched “light” and “heavy” biotin-azide capture reagents. Following the workflow shown in **Figure 6A**, out of 8925 total cysteines quantified, we find that 1070 total unique cysteines showed log2(H/L) values greater than 1.5, indicative of labeling by **PFTµ**. Included in this list were several known targets, including PARP1, HSP70 (HSPA1A), and PRDX proteins (PRDX1, PRDX2, PRDX5), which were previously identified via chemoproteomics using a clickable **PFTµ** analog^61^, providing evidence of the robustness of our approach (**Table S2**). Quite strikingly, several caspases stood out as having high log2(H/L) ratios (**Figure 6B**), including caspase-8, which aligns with our aforementioned activity assay and gel-based ABPP analysis (**Figure 5I** and **Figure S40**). Aligning with our prior discovery^24^ of structurally-related phenylpropiolate molecules that label caspase-2 at C370, we also observe that C370 is highly sensitive to **PFTµ**, which is also consistent with our gel-based analysis (**Figure S41**). The noncatalytic cysteine residue, C264, in caspase-6^26^, was also observed to be labeled by **PFTµ**, as was the near-active site cysteine (C170) in caspase-3. We did not detect the catalytic cysteine nucleophile of caspase-3 (C163). Beyond these pro-apoptotic caspases, the catalytic cysteine in the pro-inflammatory caspase-4 (C258) was also labeled by **PFTµ**. The catalytic cysteine in caspase-10 was not identified, likely due to the long tryptic peptide that flanks this residue. Taken together, these data provide further evidence that **PFTµ** promiscuously labels many human caspases in addition to its previously reported targets.

**Figure 6.**
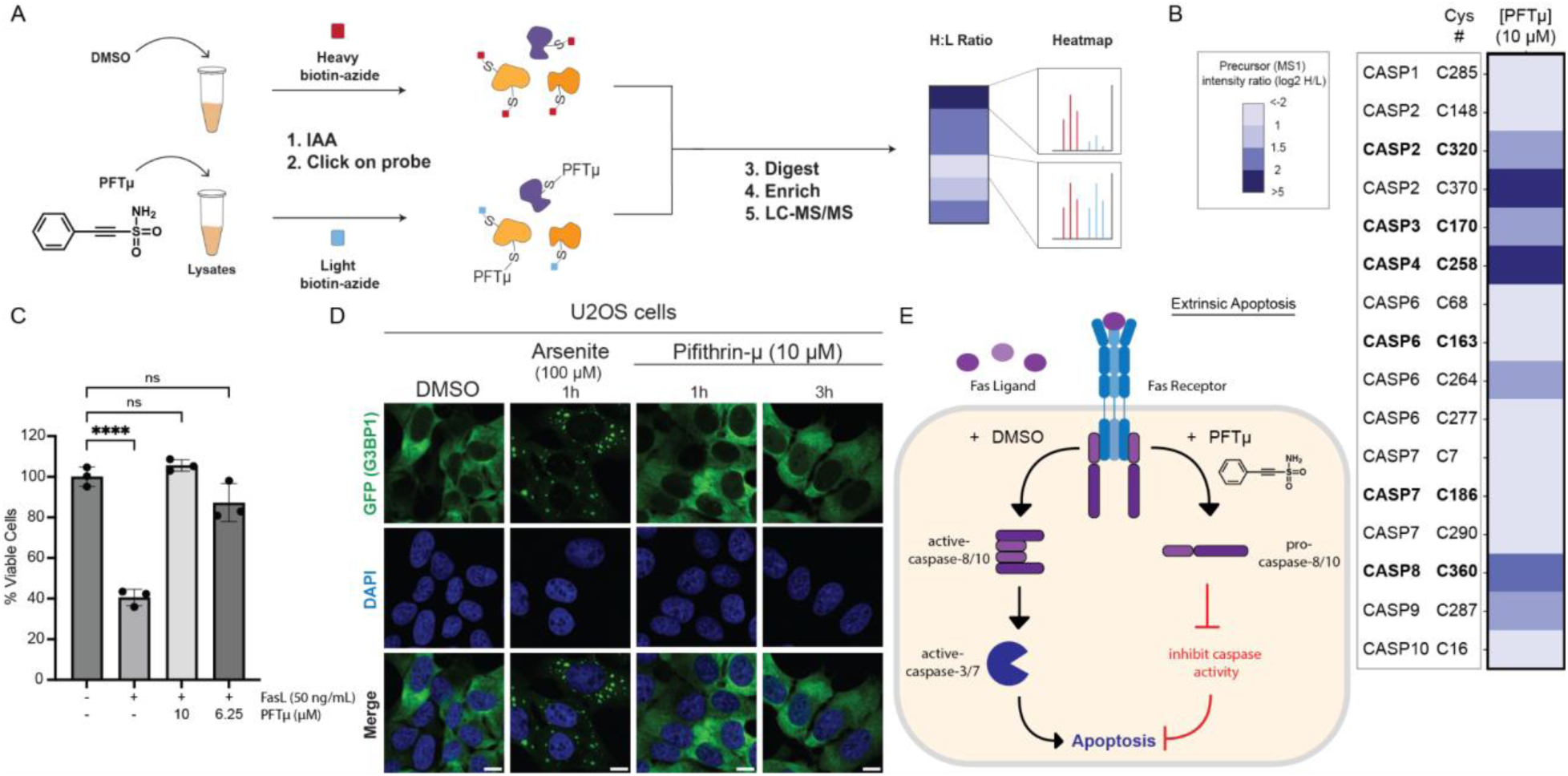
PFTµ labels initiator caspases and can protect Jurkat cells from FasL-mediated apoptosis. (A, B) Cysteine chemoproteomic analysis identifies protein targets of **PFTµ**. ‘A’ shows the general workflow in which Jurkat cell lysates were treated with either **PFTµ** (25 µM) or DMSO for 1h, followed by IAA (200 µM) cysteine capping, conjugated via click chemistry to isotopically differentiated biotin-azide reagents^73, 74^, Single-Pot Solid-Phase-enhanced sample-preparation (SP3)^60, 75, 76^ with on-resin tryptic digestion, enriched, and analyzed by LC-MS/MS. Heatmap in ‘B’ shows the mean quantified precursor intensity (log2H/L) for DMSO (H, heavy) versus compound treatment (L, light) for all caspase cysteines identified. Catalytic cysteine residues are annotated in bold. (C) CellTiter-Glo® measurement of relative viability of Jurkat cells subjected to **PFTµ** at the indicated concentrations for 1h followed by FasL (50 ng/mL, 3h)-induced apoptosis. (D) Fluorescent microscopy of U2OS cells expressing GFP-G3BP1 and treated with **PFTµ** (10 µM) or positive control sodium arsenite. All scale bars = 10 µm. (E) Proposed model where **PFTµ** functions as a promiscuous initiator caspase inhibitor (labeling caspase-2, -8, and -10) and blocks FasL-mediated extrinsic apoptosis. For B, experiments were conducted in four biological replicates, with two samples additionally analyzed as technical replicates. For C, data represent mean values ± standard deviation for three biological replicates. All MS data can be found in **Table S2**. Statistical significance was calculated with unpaired Student’s t-tests, ****p<0.0001, ns, not significant p>0.05.

Guided by these findings, we next asked whether **PFTµ** could protect cells from extrinsic apoptosis induced by Fas ligand (FasL). While **PFTµ**’s anti-apoptotic activity has been reported in other contexts^47, 62–65^, we selected FasL-mediated apoptosis as a model system due to the central role that caspase-8 and -10 play in this process^12, 66, 67^ and the less central role of TP53^68, 69^, with the goal of helping to define better the biologically active target(s) of **PFTµ**’s anti-apoptotic activity. Using the CellTiter-Glo assay, we find that **PFTµ** affords near-complete protection from FasL (**Figure 6C**) without inducing cytotoxic effects (**Figure S42**). These data align with caspase inhibition as contributing to **PFTµ**’s activity.

As our recent work has shown that electrophile stress can lead to the formation of stress granules^70^, we also opted to rule out **PFTµ**-induced stress granules as a potential confounding variable. Using a ΔΔG3BP1/2 KO cell line that stably expresses GFP-G3BP1^71, 72^, we find that **PFTµ** does not induce G3BP1 condensates using concentrations that are protective from apoptosis (**Figure 6D**) and are consistent with prior studies using **PFTµ** as a putative TP53 inhibitor^47^. Thus, we put forth a model (**Figure 6E**) that caspases are likely biological targets of **PFTµ** that contribute to the reported anti-apoptotic activity.

## Discussion

To enable high throughput procaspase-10 inhibitor discovery, here we developed and applied an engineered TEV-activatable caspase-10 protein to high throughput screening. Our high quality (an average Z’ of 0.58) ∼100,000 compound semi-automated screen yielded 237 total hits with z-score < -3. Subsequent rescreening and counter-screening delineated bona fide procaspase-10 inhibitors from those with activity against active caspase-10 or TEV protease. Orthogonal mode of action studies confirmed target engagement for both recombinant and endogenous procaspase-10. Thus, we expect that the TEV-activatable screening platform should prove broadly useful in the discovery of inhibitors targeting precursor proteases.

Our work also highlights several intriguing opportunities for future screening efforts. As TEV protease is a cysteine protease, we found that several of our hits functioned via engaging TEV rather than procaspase-10. We expect that these newly identified TEV protease inhibitors could serve as useful tools for synthetic biology studies that rely on TEV protease activity to control engineered circuits. To favor caspase rather than TEV inhibitor discovery, future studies could also consider using alternative non-cysteine protease enzymes for activation, which we expect would prove more substantially orthogonal to caspase cysteine protease activity. In addition to uncovering candidate TEV inhibitors, our screen also revealed that the thiadiazine chemotype is likely prone to disulfide rearrangement, as exemplified by our characterization of **SO265s**, which we found was the active species. Thus, our work both highlights the value of resynthesis in confirming screening hits and hints at the likely amenability of procaspase-10 to a disulfide trapping/tethering-style inhibitor discovery^77–79^ using additional disulfide-containing compounds.

Beyond the **SO265** thiadiazine chemotype, **PFTµ**, a reported TP53 inhibitor, proved to be a key hit from our screen. Comparing active and procaspase-10 inhibitor activity, we found that **PFTµ** inhibited both proteoforms to some degree, with increased inhibition observed for the proenzyme. Looking beyond caspase-10, the *in vitro*, cell-based, and proteomic analysis together confirmed that **PFTµ** engages caspase-8, -3, -10, -5, and -6. These data allow us to put forth a model whereby **PFTµ**’s reported activity as an anti-apoptotic agent is likely driven by caspase inhibition in addition to the previously reported TP53 inhibitory activity^47^. Therefore, we anticipate that the cysteine-reactive ethynesulfonamide chemotype will prove broadly useful as a starting point for caspase inhibitor development campaigns. We also expect our work to aid in phenotype interpretation for studies that have employed PFTµ to block p53 activity.

## Acknowledgements

This study was supported by National Institutes of Health (DP2 GM146246-02 to K.M.B.), Beckman Young Investigator Award (to K.M.B.), UCLA DOE Institute (DE-FC02-02ER63421 to K.M.B.), NSF GRFP (1000235263 to J.O.C), UCLA Chemistry Biology Interface Training Program (T32GM136614 to A.R.J. and M.F.P)), UCLA Cellular and Molecular Biology Training Program (T32GM145388-03 to N.P), Clinical and Translational Sciences Institute Award Core voucher (to K.M.B.) as part of the NIH/National Center for Advancing Translational Science (UL1TR001881 to UCLA). The UCLA Molecular Screening Shared Resource is supported by Jonsson Comprehensive Cancer Center, award number P30CA016042 by the National Cancer Institute of the National Institutes of Health. We thank all members of the Backus lab for their helpful suggestions, Michael Lenardo for providing plasmids encoding caspase-10 and Dennis Wolan for providing the **Rho-DEVD-AOMK** probe.

## DATA AVAILABILITY

The MS data have been deposited to the ProteomeXchange Consortium (http://proteomecentral.proteomexchange.org) via the PRIDE partner repository with the dataset identifier PXD053315 Username: Password: ut5UBouFBb4x

## References

(1) Horn, S.; Hughes, M. A.; Schilling, R.; Sticht, C.; Tenev, T.; Ploesser, M.; Meier, P.; Sprick, M. R.; Macfarlane, M.; Leverkus, M. Caspase-10 Negatively Regulates Caspase-8-Mediated Cell Death, Switching the Response to CD95L in Favor of NF-κB Activation and Cell Survival. Cell Reports 2017, *19* (4), 785-797. DOI: 10.1016/j.celrep.2017.04.010.

(2) Yang, C. Y.; Lien, C. I.; Tseng, Y. C.; Tu, Y. F.; Kulczyk, A. W.; Lu, Y. C.; Wang, Y. T.; Su, T. W.; Hsu, L. C.; Lo, Y. C.;, et al. Deciphering DED assembly mechanisms in FADD-procaspase- 8-cFLIP complexes regulating apoptosis. Nat Commun 2024, 15 (1), 3791. DOI: 10.1038/s41467-024-47990-2 From NLM Medline.

(3) Wang, J.; Chun, H. J.; Wong, W.; Spencer, D. M.; Lenardo, M. J. Caspase-10 is an initiator caspase in death receptor signaling. Proceedings of the National Academy of Sciences 2001, 98 (24), 13884–13888. DOI: 10.1073/pnas.241358198.

(4) Julien, O.; Wells, J. A. Caspases and their substrates. Cell Death & Differentiation 2017, 24 (8), 1380–1389. DOI: 10.1038/cdd.2017.44.

(5) Wang, L.; Main, K.; Wang, H.; Julien, O.; Dufour, A. Biochemical Tools for Tracking Proteolysis. J Proteome Res 2021, 20 (12), 5264–5279. DOI: 10.1021/acs.jproteome.1c00289 From NLM Medline.

(6) Eskandari, E.; Negri, G. L.; Tan, S.; Macaldaz, M. E.; Ding, S.; Long, J.; Nielsen, K.; Spencer, S. E.; Morin, G. B.; Eaves, C. J. Dependence of human cell survival and proliferation on the CASP3 prodomain. Cell Death Discovery 2024, 10 (1). DOI: 10.1038/s41420-024-01826-6.

(7) Alam, A.; Cohen, L. Y.; Aouad, S.; Sékaly, R.-P. Early Activation of Caspases during T Lymphocyte Stimulation Results in Selective Substrate Cleavage in Nonapoptotic Cells. The Journal of Experimental Medicine 1999, 190 (12), 1879–1890. DOI: 10.1084/jem.190.12.1879.

(8) Pistritto, G.; Jost, M.; Srinivasula, S. M.; Baffa, R.; Poyet, J. L.; Kari, C.; Lazebnik, Y.; Rodeck, U.; Alnemri, E. S. Expression and transcriptional regulation of caspase-14 in simple and complex epithelia. Cell Death & Differentiation 2002, 9 (9), 995–1006. DOI: 10.1038/sj.cdd.4401061.

(9) Lippens, S.; Kockx, M.; Knaapen, M.; Mortier, L.; Polakowska, R.; Verheyen, A.; Garmyn, M.; Zwijsen, A.; Formstecher, P.; Huylebroeck, D.;, et al. Epidermal differentiation does not involve the pro-apoptotic executioner caspases, but is associated with caspase-14 induction and processing. Cell Death & Differentiation 2000, 7 (12), 1218–1224. DOI: 10.1038/sj.cdd.4400785.

(10) Gorelick-Ashkenazi, A.; Weiss, R.; Sapozhnikov, L.; Florentin, A.; Tarayrah-Ibraheim, L.; Dweik, D.; Yacobi-Sharon, K.; Arama, E. Caspases maintain tissue integrity by an apoptosis-independent inhibition of cell migration and invasion. Nature Communications 2018, 9 (1). DOI: 10.1038/s41467-018-05204-6.

(11) Shao, W.; Yeretssian, G.; Doiron, K.; Hussain, S. N.; Saleh, M. The Caspase-1 Digestome Identifies the Glycolysis Pathway as a Target during Infection and Septic Shock. Journal of Biological Chemistry 2007, 282 (50), 36321–36329. DOI: 10.1074/jbc.m708182200.

(12) Consonni, F.; Moreno, S.; Vinuales Colell, B.; Stolzenberg, M.-C.; Fernandes, A.; Parisot, M.; Masson, C.; Neveux, N.; Rosain, J.; Bamberger, S.;, et al. Study of the potential role of CASPASE-10 mutations in the development of autoimmune lymphoproliferative syndrome. Cell Death & Disease 2024, 15 (5). DOI: 10.1038/s41419-024-06679-6.

(13) Krug, H. F. Caspase-10 is the key initiator caspase involved in tributyltin-mediated apoptosis in human immune cells. J Toxicol 2012, 2012, 395482. DOI: 10.1155/2012/395482 From NLM PubMed-not-MEDLINE.

(14) Philip, N. H.; Dillon, C. P.; Snyder, A. G.; Fitzgerald, P.; Wynosky-Dolfi, M. A.; Zwack, E. E.; Hu, B.; Fitzgerald, L.; Mauldin, E. A.; Copenhaver, A. M.;, et al. Caspase-8 mediates caspase-1 processing and innate immune defense in response to bacterial blockade of NF-κB and MAPK signaling. Proceedings of the National Academy of Sciences 2014, 111 (20), 7385–7390. DOI: 10.1073/pnas.1403252111.

(15) Rieux-Laucat, F.; Le Deist, F.; Fischer, A. Autoimmune lymphoproliferative syndromes: genetic defects of apoptosis pathways. Cell Death & Differentiation 2003, 10 (1), 124–133. DOI: 10.1038/sj.cdd.4401190.

(16) Olsson, M.; Zhivotovsky, B. Caspases and cancer. Cell Death & Differentiation 2011, 18 (9), 1441–1449. DOI: 10.1038/cdd.2011.30.

(17) Cui, Z.; Dabas, H.; Leonard, B. C.; Shiah, J. V.; Grandis, J. R.; Johnson, D. E. Caspase-8 mutations associated with head and neck cancer differentially retain functional properties related to TRAIL-induced apoptosis and cytokine induction. Cell Death & Disease 2021, 12 (8). DOI: 10.1038/s41419-021-04066-z.

(18) Jiang, M.; Qi, L.; Li, L.; Li, Y. The caspase-3/GSDME signal pathway as a switch between apoptosis and pyroptosis in cancer. Cell Death Discovery 2020, 6 (1). DOI: 10.1038/s41420-020-00349-0.

(19) Groborz, K. M.; Kalinka, M.; Grzymska, J.; Kołt, S.; Snipas, S. J.; Poręba, M. Selective chemical reagents to investigate the role of caspase 6 in apoptosis in acute leukemia T cells. Chemical Science 2023, 14 (9), 2289–2302. DOI: 10.1039/d2sc05827h.

(20) Shaulov-Rotem, Y.; Merquiol, E.; Weiss-Sadan, T.; Moshel, O.; Salpeter, S.; Shabat, D.; Kaschani, F.; Kaiser, M.; Blum, G. A novel quenched fluorescent activity-based probe reveals caspase-3 activity in the endoplasmic reticulum during apoptosis. Chemical Science 2016, 7 (2), 1322–1337. DOI: 10.1039/c5sc03207e.

(21) De Calignon, A.; Fox, L. M.; Pitstick, R.; Carlson, G. A.; Bacskai, B. J.; Spires-Jones, T. L.; Hyman, B. T. Caspase activation precedes and leads to tangles. Nature 2010, 464 (7292), 1201–1204. DOI: 10.1038/nature08890.

(22) Boxer, M. B.; Quinn, A. M.; Shen, M.; Jadhav, A.; Leister, W.; Simeonov, A.; Auld, D. S.; Thomas, C. J. A Highly Potent and Selective Caspase 1 Inhibitor that Utilizes a Key 3- Cyanopropanoic Acid Moiety. ChemMedChem 2010, 5 (5), 730–738. DOI: 10.1002/cmdc.200900531.

(23) Bosc, E.; Anastasie, J.; Soualmia, F.; Coric, P.; Kim, J. Y.; Wang, L. Q.; Lacin, G.; Zhao, K.; Patel, R.; Duplus, E.;, et al. Genuine selective caspase-2 inhibition with new irreversible small peptidomimetics. Cell Death & Disease 2022, 13 (11). DOI: 10.1038/s41419-022-05396-2.

(24) Castellón, J. O.; Ofori, S.; Burton, N. R.; Julio, A. R.; Turmon, A. C.; Armenta, E.; Sandoval, C.; Boatner, L. M.; Takayoshi, E. E.; Faragalla, M.;, et al. Chemoproteomics Identifies State-Dependent and Proteoform-Selective Caspase-2 Inhibitors. Journal of the American Chemical Society 2024, 146 (22), 14972–14988. DOI: 10.1021/jacs.3c12240.

(25) Tubeleviciute-Aydin, A.; Beautrait, A.; Lynham, J.; Sharma, G.; Gorelik, A.; Deny, L. J.; Soya, N.; Lukacs, G. L.; Nagar, B.; Marinier, A.;, et al. Identification of Allosteric Inhibitors against Active Caspase-6. Scientific Reports 2019, 9 (1). DOI: 10.1038/s41598-019-41930-7.

(26) Van Horn, K. S.; Wang, D.; Medina-Cleghorn, D.; Lee, P. S.; Bryant, C.; Altobelli, C.; Jaishankar, P.; Leung, K. K.; Ng, R. A.; Ambrose, A. J.;, et al. Engaging a Non-catalytic Cysteine Residue Drives Potent and Selective Inhibition of Caspase-6. Journal of the American Chemical Society 2023, 145 (18), 10015–10021. DOI: 10.1021/jacs.2c12240.

(27) Backus, K. M.; Correia, B. E.; Lum, K. M.; Forli, S.; Horning, B. D.; González-Páez, G. E.; Chatterjee, S.; Lanning, B. R.; Teijaro, J. R.; Olson, A. J.;, et al. Proteome-wide covalent ligand discovery in native biological systems. Nature 2016, 534 (7608), 570–574. DOI: 10.1038/nature18002.

(28) Bucur, O.; Gaidos, G.; Yatawara, A.; Pennarun, B.; Rupasinghe, C.; Roux, J.; Andrei, S.; Guo, B.; Panaitiu, A.; Pellegrini, M.;, et al. A novel caspase 8 selective small molecule potentiates TRAIL-induced cell death. Scientific Reports 2015, 5 (1), 9893. DOI: 10.1038/srep09893.

(29) Dhani, S.; Zhao, Y.; Zhivotovsky, B. A long way to go: caspase inhibitors in clinical use. Cell Death & Disease 2021, 12 (10). DOI: 10.1038/s41419-021-04240-3.

(30) Lopez-Hernandez, F. J.; Ortiz, M. A.; Bayon, Y.; Piedrafita, F. J. Z-FA-fmk inhibits effector caspases but not initiator caspases 8 and 10, and demonstrates that novel anticancer retinoid-related molecules induce apoptosis via the intrinsic pathway. Mol Cancer Ther 2003, 2 (3), 255–263. From NLM.

(31) Jänicke, R. U.; Sohn, D.; Totzke, G.; Schulze-Osthoff, K. Caspase-10 in mouse or not? Science 2006, 312 (5782), 1874. DOI: 10.1126/science.312.5782.1874a From NLM.

(32) Sprick, M. R.; Weigand, M. A.; Rieser, E.; Rauch, C. T.; Juo, P.; Blenis, J.; Krammer, P. H.; Walczak, H. FADD/MORT1 and Caspase-8 Are Recruited to TRAIL Receptors 1 and 2 and Are Essential for Apoptosis Mediated by TRAIL Receptor 2. Immunity 2000, 12 (6), 599–609. DOI: 10.1016/s1074-7613(00)80211-3.

(33) Bodmer, J.-L.; Holler, N.; Reynard, S.; Vinciguerra, P.; Schneider, P.; Juo, P.; Blenis, J.; Tschopp, J. TRAIL receptor-2 signals apoptosis through FADD and caspase-8. Nature Cell Biology 2000, 2 (4), 241–243. DOI: 10.1038/35008667.

(34) Juo, P.; Kuo, C. J.; Yuan, J.; Blenis, J. Essential requirement for caspase-8/FLICE in the initiation of the Fas-induced apoptotic cascade. Current Biology 1998, 8 (18), 1001–1008. DOI: 10.1016/s0960-9822(07)00420-4.

(35) Kawahara, A.; Ohsawa, Y.; Matsumura, H.; Uchiyama, Y.; Nagata, S. Caspase-independent Cell Killing by Fas-associated Protein with Death Domain. The Journal of Cell Biology 1998, 143 (5), 1353–1360. DOI: 10.1083/jcb.143.5.1353.

(36) Grotzer, M. A.; Eggert, A.; Zuzak, T. J.; Janss, A. J.; Marwaha, S.; Wiewrodt, B. R.; Ikegaki, N.; Brodeur, G. M.; Phillips, P. C. Resistance to TRAIL-induced apoptosis in primitive neuroectodermal brain tumor cells correlates with a loss of caspase-8 expression. Oncogene 2000, 19 (40), 4604–4610. DOI: 10.1038/sj.onc.1203816.

(37) Teitz, T.; Wei, T.; Valentine, M. B.; Vanin, E. F.; Grenet, J.; Valentine, V. A.; Behm, F. G.; Look, A. T.; Lahti, J. M.; Kidd, V. J. Caspase 8 is deleted or silenced preferentially in childhood neuroblastomas with amplification of MYCN. Nature Medicine 2000, 6 (5), 529–535. DOI: 10.1038/75007.

(38) Mühlethaler-Mottet, A.; Flahaut, M.; Bourloud, K. B.; Nardou, K.; Coulon, A.; Liberman, J.; Thome, M.; Gross, N. Individual caspase-10 isoforms play distinct and opposing roles in the initiation of death receptor-mediated tumour cell apoptosis. Cell Death & Disease 2011, 2 (3), e125–e125. DOI: 10.1038/cddis.2011.8.

(39) Chun, H. J.; Zheng, L.; Ahmad, M.; Wang, J.; Speirs, C. K.; Siegel, R. M.; Dale, J. K.; Puck, J.; Davis, J.; Hall, C. G.;, et al. Pleiotropic defects in lymphocyte activation caused by caspase-8 mutations lead to human immunodeficiency. Nature 2002, 419 (6905), 395–399. DOI: 10.1038/nature01063.

(40) Grønbaek, K.; Dalby, T.; Zeuthen, J.; Ralfkiaer, E.; Guidberg, P. The V410I (G1228A) variant of the caspase-10 gene is a common polymorphism of the Danish population. Blood 2000, 95 (6), 2184–2185. From NLM.

(41) Nagar, B.; Bornmann, W. G.; Pellicena, P.; Schindler, T.; Veach, D. R.; Miller, W. T.; Clarkson, B.; Kuriyan, J. Crystal Structures of the Kinase Domain of c-Abl in Complex with the Small Molecule Inhibitors PD173955 and Imatinib (STI-571)1. Cancer Research 2002, 62 (15), 4236–4243. (acccessed 7/26/2024).

(42) Capdeville, R.; Buchdunger, E.; Zimmermann, J.; Matter, A. Glivec (STI571, imatinib), a rationally developed, targeted anticancer drug. Nature Reviews Drug Discovery 2002, 1 (7), 493–502. DOI: 10.1038/nrd839.

(43) van Montfort, R. L. M.; Workman, P. Structure-based design of molecular cancer therapeutics. Trends in Biotechnology 2009, 27 (5), 315–328. DOI: 10.1016/j.tibtech.2009.02.003.

(44) Lin, Y.-L.; Meng, Y.; Jiang, W.; Roux, B. Explaining why Gleevec is a specific and potent inhibitor of Abl kinase. Proceedings of the National Academy of Sciences 2013, 110 (5), 1664–1669. DOI: 10.1073/pnas.1214330110.

(45) Xu, J. H.; Eberhardt, J.; Hill-Payne, B.; González-Páez, G. E.; Castellón, J. O.; Cravatt, B. F.; Forli, S.; Wolan, D. W.; Backus, K. M. Integrative X-ray Structure and Molecular Modeling for the Rationalization of Procaspase-8 Inhibitor Potency and Selectivity. ACS Chemical Biology 2020, 15 (2), 575–586. DOI: 10.1021/acschembio.0c00019.

(46) Zhang, J. H.; Chung, T. D.; Oldenburg, K. R. A Simple Statistical Parameter for Use in Evaluation and Validation of High Throughput Screening Assays. J Biomol Screen 1999, 4 (2), 67–73. DOI: 10.1177/108705719900400206 From NLM Publisher.

(47) Strom, E.; Sathe, S.; Komarov, P. G.; Chernova, O. B.; Pavlovska, I.; Shyshynova, I.; Bosykh, D. A.; Burdelya, L. G.; Macklis, R. M.; Skaliter, R.;, et al. Small-molecule inhibitor of p53 binding to mitochondria protects mice from gamma radiation. Nat Chem Biol 2006, 2 (9), 474–479. DOI: 10.1038/nchembio809 From NLM Medline.

(48) Gray, D. C.; Mahrus, S.; Wells, J. A. Activation of Specific Apoptotic Caspases with an Engineered Small-Molecule-Activated Protease. Cell 2010, 142 (4), 637–646. DOI: 10.1016/j.cell.2010.07.014.

(49) Oberst, A.; Pop, C.; Tremblay, A. G.; Blais, V.; Denault, J.-B.; Salvesen, G. S.; Green, D. R. Inducible Dimerization and Inducible Cleavage Reveal a Requirement for Both Processes in Caspase-8 Activation. Journal of Biological Chemistry 2010, 285 (22), 16632–16642. DOI: 10.1074/jbc.m109.095083.

(50) Kim, K.; Damoiseaux, R.; Norris, A. J.; Rivina, L.; Bradley, K.; Jung, M. E.; Gatti, R. A.; Schiestl, R. H.; McBride, W. H. High throughput screening of small molecule libraries for modifiers of radiation responses. International Journal of Radiation Biology 2011, 87 (8), 839–845. DOI: 10.3109/09553002.2011.560994.

(51) Sander, T.; Freyss, J.; von Korff, M.; Rufener, C. DataWarrior: an open-source program for chemistry aware data visualization and analysis. J Chem Inf Model 2015, 55 (2), 460–473. DOI: 10.1021/ci500588j From NLM.

(52) Pierce, J. W.; Schoenleber, R.; Jesmok, G.; Best, J.; Moore, S. A.; Collins, T.; Gerritsen, M. E. Novel Inhibitors of Cytokine-induced IκBα Phosphorylation and Endothelial Cell Adhesion Molecule Expression Show Anti-inflammatory Effects in Vivo. Journal of Biological Chemistry 1997, 272 (34), 21096–21103. DOI: 10.1074/jbc.272.34.21096.

(53) Kraft, M.; Radke, D.; Wieland, G. D.; Zipfel, P. F.; Horn, U. A fluorogenic substrate as quantitative in vivo reporter to determine protein expression and folding of tobacco etch virus protease in Escherichia coli. Protein Expression and Purification 2007, 52 (2), 478–484. DOI: 10.1016/j.pep.2006.10.019.

(54) Nam, H.; Hwang, B. J.; Choi, D. Y.; Shin, S.; Choi, M. Tobacco etch virus (TEV) protease with multiple mutations to improve solubility and reduce self-cleavage exhibits enhanced enzymatic activity. FEBS Open Bio 2020, 10 (4), 619–626. DOI: 10.1002/2211-5463.12828.

(55) Gerasimova, E. L.; Gazizullina, E. G.; Igdisanova, D. I.; Sidorova, L. P.; Tseitler, T. A.; Emelianov, V. V.; Chupakhin, O. N.; Ivanova, A. V. Antioxidant properties of 2,5-substituted 6H- 1,3,4-thiadiazines promising for experimental therapy of diabetes mellitus. Russian Chemical Bulletin 2022, *71* (12), 2730-2739. DOI: 10.1007/s11172-022-3702-0.

(56) Yiannios, C. N.; Karabinos, J. V. Oxidation of Thiols by Dimethyl Sulfoxide. The Journal of Organic Chemistry 1963, 28 (11), 3246–3248. DOI: 10.1021/jo01046a528.

(57) Wallace, T. J.; Mahon, J. J. Reactions of Thiols with Sulfoxides. II. Kinetics and Mechanistic Implications. Journal of the American Chemical Society 1964, 86 (19), 4099–4103. DOI: 10.1021/ja01073a039.

(58) Jafari, R.; Almqvist, H.; Axelsson, H.; Ignatushchenko, M.; Lundbäck, T.; Nordlund, P.; Molina, D. M. The cellular thermal shift assay for evaluating drug target interactions in cells. Nature Protocols 2014, 9 (9), 2100–2122. DOI: 10.1038/nprot.2014.138.

(59) Ghosh, R.; Gilda, J. E.; Gomes, A. V. The necessity of and strategies for improving confidence in the accuracy of western blots. Expert Review of Proteomics 2014, 11 (5), 549–560. DOI: 10.1586/14789450.2014.939635.

(60) Yan, T.; Desai, H. S.; Boatner, L. M.; Yen, S. L.; Cao, J.; Palafox, M. F.; Jami-Alahmadi, Y.; Backus, K. M. SP3-FAIMS Chemoproteomics for High-Coverage Profiling of the Human Cysteinome**. ChemBioChem 2021, 22 (10), 1841–1851. DOI: 10.1002/cbic.202000870.

(61) Yang, J.; Liu, Z.; Perrett, S.; Zhang, H.; Pan, Z. PES derivative PESA is a potent tool to globally profile cellular targets of PES. Bioorg Med Chem Lett 2022, 60, 128553. DOI: 10.1016/j.bmcl.2022.128553 From NLM Medline.

(62) Maj, M. A.; Ma, J.; Krukowski, K. N.; Kavelaars, A.; Heijnen, C. J. Inhibition of Mitochondrial p53 Accumulation by PFT-mu Prevents Cisplatin-Induced Peripheral Neuropathy. Front Mol Neurosci 2017, 10, 108. DOI: 10.3389/fnmol.2017.00108 From NLM PubMed-not-MEDLINE.

(63) Sekihara, K.; Harashima, N.; Tongu, M.; Tamaki, Y.; Uchida, N.; Inomata, T.; Harada, M. Pifithrin-mu, an inhibitor of heat-shock protein 70, can increase the antitumor effects of hyperthermia against human prostate cancer cells. PLoS One 2013, 8 (11), e78772. DOI: 10.1371/journal.pone.0078772 From NLM Medline.

(64) Yang, L. Y.; Greig, N. H.; Tweedie, D.; Jung, Y. J.; Chiang, Y. H.; Hoffer, B. J.; Miller, J. P.; Chang, K. H.; Wang, J. Y. The p53 inactivators pifithrin-mu and pifithrin-alpha mitigate TBI-induced neuronal damage through regulation of oxidative stress, neuroinflammation, autophagy and mitophagy. Exp Neurol 2020, 324, 113135. DOI: 10.1016/j.expneurol.2019.113135 From NLM Medline.

(65) Zhu, J.; Singh, M.; Selivanova, G.; Peuget, S. Pifithrin-alpha alters p53 post-translational modifications pattern and differentially inhibits p53 target genes. Sci Rep 2020, 10 (1), 1049. DOI: 10.1038/s41598-020-58051-1 From NLM Medline.

(66) Coe, G. L.; Redd, P. S.; Paschall, A. V.; Lu, C.; Gu, L.; Cai, H.; Albers, T.; Lebedyeva, I. O.; Liu, K. Ceramide mediates FasL-induced caspase 8 activation in colon carcinoma cells to enhance FasL-induced cytotoxicity by tumor-specific cytotoxic T lymphocytes. Scientific Reports 2016, 6 (1), 30816. DOI: 10.1038/srep30816.

(67) Milhas, D.; Cuvillier, O.; Therville, N.; Clavé, P.; Thomsen, M.; Levade, T.; Benoist, H.; Ségui, B. Caspase-10 Triggers Bid Cleavage and Caspase Cascade Activation in FasL-induced Apoptosis. Journal of Biological Chemistry 2005, 280 (20), 19836–19842. DOI: 10.1074/jbc.m414358200.

(68) Uriarte, S. M.; Joshi-Barve, S.; Song, Z.; Sahoo, R.; Gobejishvili, L.; Jala, V. R.; Haribabu, B.; McClain, C.; Barve, S. Akt inhibition upregulates FasL, downregulates c-FLIPs and induces caspase-8-dependent cell death in Jurkat T lymphocytes. Cell Death & Differentiation 2005, 12 (3), 233–242. DOI: 10.1038/sj.cdd.4401549.

(69) Mohr, A.; Deedigan, L.; Jencz, S.; Mehrabadi, Y.; Houlden, L.; Albarenque, S.-M.; Zwacka, R. M. Caspase-10: a molecular switch from cell-autonomous apoptosis to communal cell death in response to chemotherapeutic drug treatment. Cell Death & Differentiation 2018, 25 (2), 340–352. DOI: 10.1038/cdd.2017.164.

(70) Julio, A. R.; Shikwana, F.; Truong, C.; Burton, N. R.; Dominguez, E. R.; Turmon, A. C.; Cao, J.; Backus, K. M. Delineating cysteine-reactive compound modulation of cellular proteostasis processes. Nature Chemical Biology 2024. DOI: 10.1038/s41589-024-01760-9.

(71) Kedersha, N.; Panas, M. D.; Achorn, C. A.; Lyons, S.; Tisdale, S.; Hickman, T.; Thomas, M.; Lieberman, J.; McInerney, G. M.; Ivanov, P.;, et al. G3BP–Caprin1–USP10 complexes mediate stress granule condensation and associate with 40S subunits. Journal of Cell Biology 2016, 212 (7), 845–860. DOI: 10.1083/jcb.201508028.

(72) Götte, B.; Panas, M. D.; Hellström, K.; Liu, L.; Samreen, B.; Larsson, O.; Ahola, T.; McInerney, G. M. Separate domains of G3BP promote efficient clustering of alphavirus replication complexes and recruitment of the translation initiation machinery. PLOS Pathogens 2019, 15 (6), e1007842. DOI: 10.1371/journal.ppat.1007842.

(73) Shikwana, F.; Heydari, B.; Ofori, S.; Truong, C.; Turmon, A.; Darrouj, J.; Holoidovsky, L.; Gustafson, J.; Backus, K. CySP3-96 enables scalable, streamlined, and low-cost sample preparation for cysteine chemoproteomic applications. American Chemical Society (ACS): 2024.

(74) Yan, T.; Palmer, A. B.; Geiszler, D. J.; Polasky, D. A.; Boatner, L. M.; Burton, N. R.; Armenta, E.; Nesvizhskii, A. I.; Backus, K. M. Enhancing Cysteine Chemoproteomic Coverage through Systematic Assessment of Click Chemistry Product Fragmentation. Analytical Chemistry 2022, 94 (9), 3800–3810. DOI: 10.1021/acs.analchem.1c04402.

(75) Hughes, C. S.; Foehr, S.; Garfield, D. A.; Furlong, E. E.; Steinmetz, L. M.; Krijgsveld, J. Ultrasensitive proteome analysis using paramagnetic bead technology. Molecular Systems Biology 2014, 10 (10), 757. DOI: 10.15252/msb.20145625.

(76) Hughes, C. S.; Moggridge, S.; Müller, T.; Sorensen, P. H.; Morin, G. B.; Krijgsveld, J. Single-pot, solid-phase-enhanced sample preparation for proteomics experiments. Nature Protocols 2019, 14 (1), 68–85. DOI: 10.1038/s41596-018-0082-x.

(77) Hallenbeck, K. K.; Davies, J. L.; Merron, C.; Ogden, P.; Sijbesma, E.; Ottmann, C.; Renslo, A. R.; Wilson, C.; Arkin, M. R. A Liquid Chromatography/Mass Spectrometry Method for Screening Disulfide Tethering Fragments. SLAS Discovery 2018, 23 (2), 183–192. DOI: 10.1177/2472555217732072.

(78) Sadowsky, J. D.; Burlingame, M. A.; Wolan, D. W.; McClendon, C. L.; Jacobson, M. P.; Wells, J. A. Turning a protein kinase on or off from a single allosteric site via disulfide trapping. Proceedings of the National Academy of Sciences 2011, 108 (15), 6056–6061. DOI: 10.1073/pnas.1102376108.

(79) Erlanson, D. A.; Braisted, A. C.; Raphael, D. R.; Randal, M.; Stroud, R. M.; Gordon, E. M.; Wells, J. A. Site-directed ligand discovery. Proceedings of the National Academy of Sciences 2000, 97 (17), 9367–9372. DOI: 10.1073/pnas.97.17.9367.

