## Supporting Information for "An activation-based high throughput screen identifies caspase-10 inhibitors"

**Table of Contents**


| | | |
|---|---|---|
| (A) | Supplementary Schemes | 02 - 03 |
| (B) | Supplementary Figures | 03 - 44 |
| (C) | Supplementary Tables | 45 - 48 |
| (D) | Biology Methods | 48 - 57 |
| (E) | Chemistry Methods | 57 - 61 |
| (F) | NMR Spectra | 61 - 70 |
| (G) | References | 71 |

### (A) Supplementary Schemes

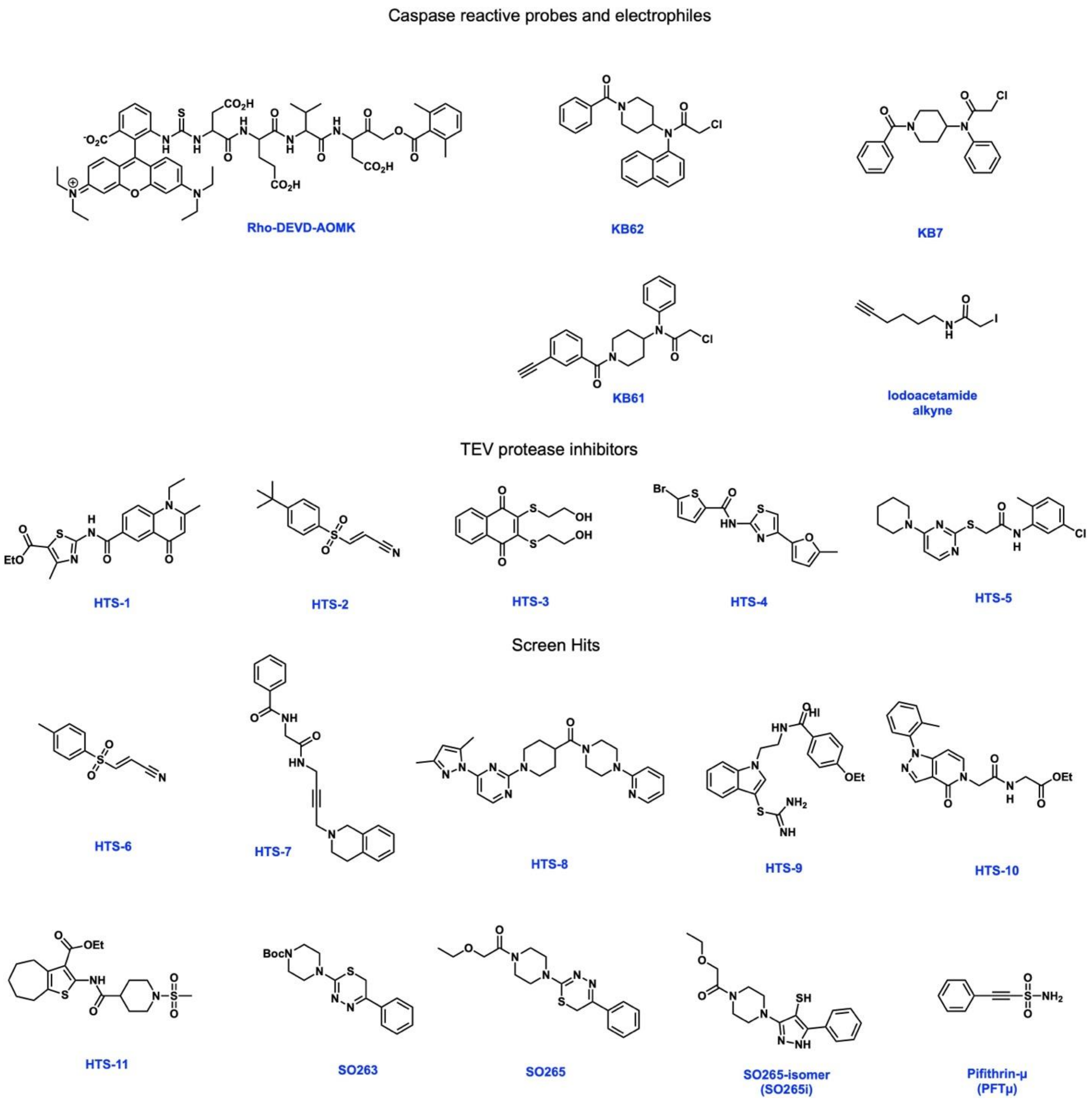


**Scheme S1. Structure of compounds used in this study,** including previously reported compounds KB7, KB61, and KB62[1].

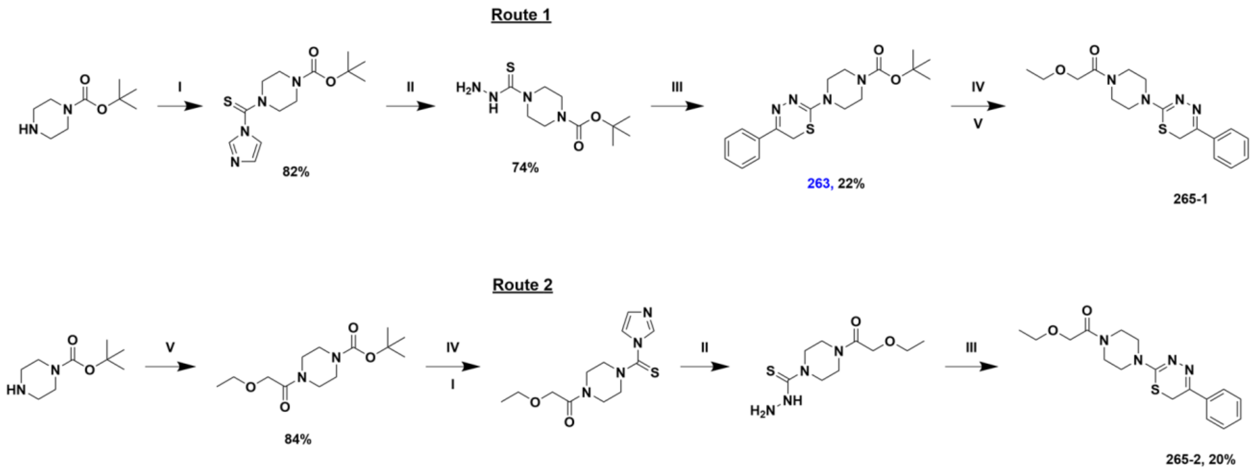


**Scheme S2: Synthesis of 263, 265-1, 265-2 :** (I)1,1′-Thiocarbonyldiimidazole, $CH_2Cl_2$, rt, 24 h (II) Hydrazine monohydrate, EtOH, 80$^oC$, 3 h (III) 2-Bromoacetophenone, EtOH, 80 $^oC$, 0.5 h (IV) TFA, $CH_2Cl_2$, rt, 1-2 h., (V) 2-Ethoxyacetyl chloride, $Et_3N$, $CH_2Cl_2$, 0$^oC$ - rt, 18 h.

### (B) Supplementary Figures

```
Wildtype                FGAVYSSDEALIPIREIMSHFTALQCPRLAEKPKLFFIQACQGEEIQPSVSIEAD-----    415
procASP10TEVx2          FGAVYSSDEALIPIREIMSHFTALQCPRLAEKPKLFFIQACQGEEIQPSVSIE--AENLY    418
proCAPS10 C401A         FGAVYSSDEALIPIREIMSHFTALQCPRLAEKPKLFFIQAAQGEEIQPSVSIEAD-----    415
proCASP10TEV            FGAVYSSDEALIPIREIMSHFTALQCPRLAEKPKLFFIQACQGEEIQPSVSIE--AENLY    418
proCASP10TEV Linker     FGAVYSSDEALIPIREIMSHFTALQCPRLAEKPKLFFIQACQGEEIQPSVSIEAAAENLY    420
proCASP10TEV D435A      FGAVYSSDEALIPIREIMSHFTALQCPRLAEKPKLFFIQACQGEEIQPSVSIE--AENLY    418
                        ****************************************.************

Wildtype                ---ALNPEQAPTSLQDSIPAEA------DFLLGLATVPGYVSFRHVEEGSWYIQSLCNHL    466
procASP10TEVx2          FQGALNPEQAPTSLQDSIPAEAENLYFQGFLLGLATVPGYVSFRHVEEGSWYIQSLCNHL    478
proCAPS10 C401A         ---ALNPEQAPTSLQDSIPAEA------DFLLGLATVPGYVSFRHVEEGSWYIQSLCNHL    466
proCASP10TEV            FQGALNPEQAPTSLQDSIPAEA------DFLLGLATVPGYVSFRHVEEGSWYIQSLCNHL    472
proCASP10TEV Linker     FQGALNPEQAPTSLQDSIPAEA------DFLLGLATVPGYVSFRHVEEGSWYIQSLCNHL    474
proCASP10TEV D435A      FQGALNPEQAPTSLQDSIPAE-------AFLLGLATVPGYVSFRHVEEGSWYIQSLCNHL    471
                           *******************       .********************************
```

**Figure S1. Sequences modified to generate the indicated engineered caspase-10 proteins**. Shows sequence alignments for the amino acid region of caspase-10 (Q92851-4, 10-L isoform) engineered to insert the indicated mutations and TEV cleavage motifs. Catalytic cysteine (C401) is highlighted in yellow. residue highlighted in green indicates proteolytic target aspartate residue, and pink residues correspond to the TEV protease proteolytic recognition sequence. The suspected additional cleavage site, D435, was replaced with an additional TEV cleavage motif (proCASP10TEVx2).

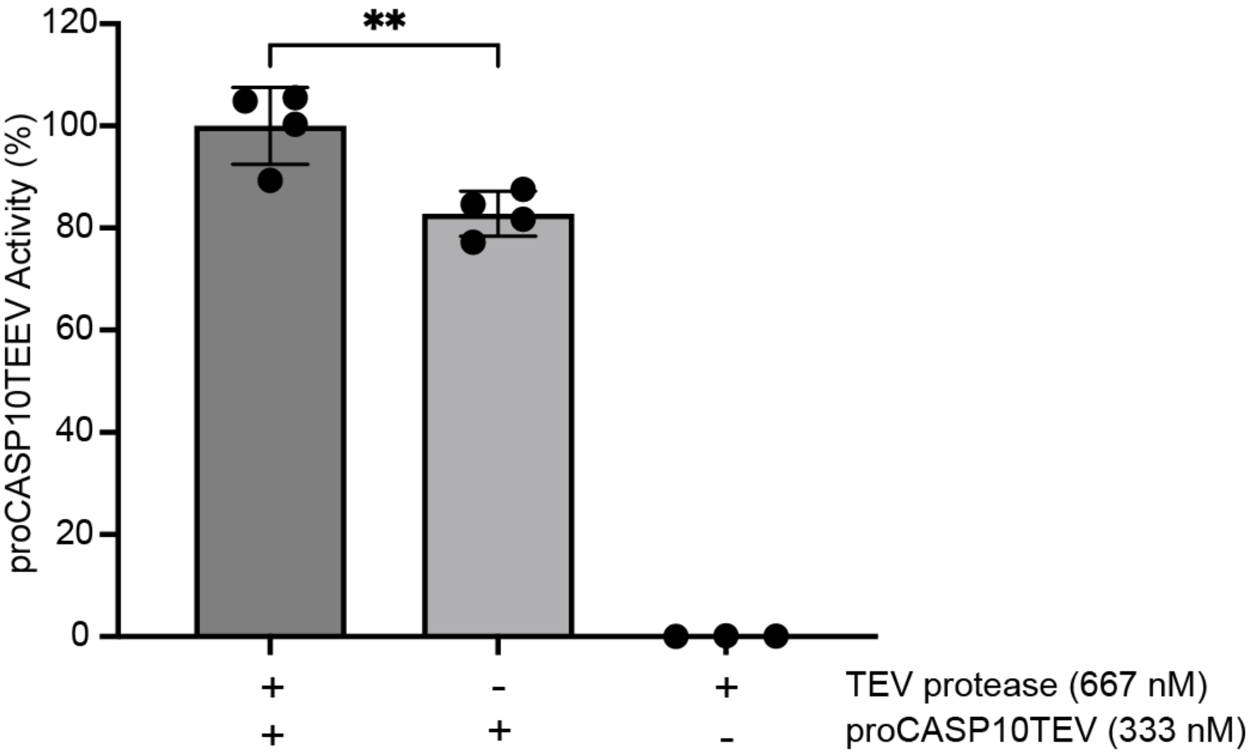


**Figure S2**. **proCASP10TEV has higher background activity in the absence of TEV protease**. Activity of a proCASP10TEV (333 nM final concentration) stored in the -80°C after treatment with TEV protease (667 nM) solution containing 333 mM citrate, 5 mM DTT, and 10 μM **Ac-VDVAC-AFC** substrate in PBS. Data represents mean values and standard deviation (n = 4 biological replicates). Statistical significance was calculated with unpaired Student's t-tests, **p<0.001.

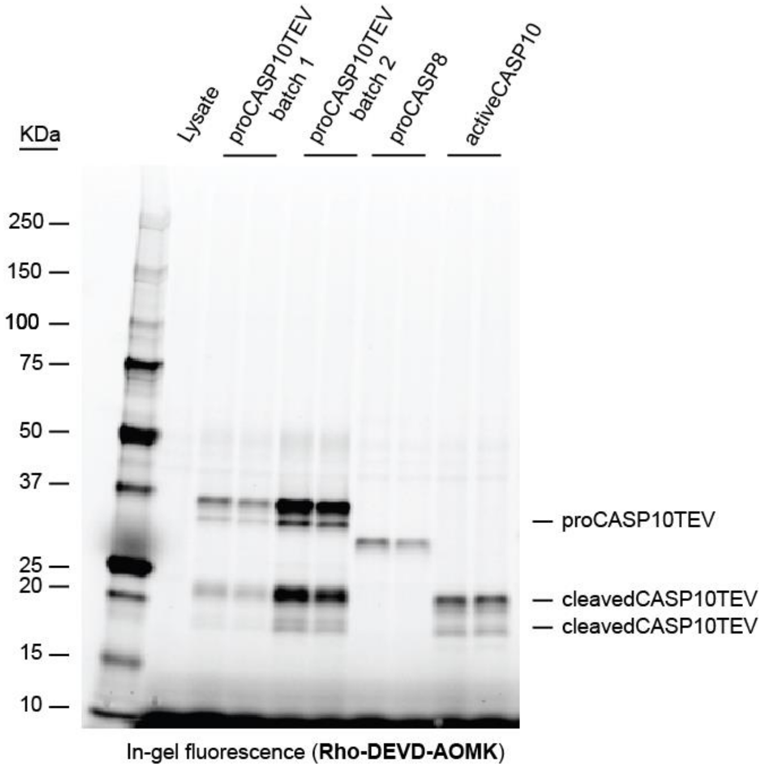


**Figure S3**. **proCASP10TEV self-activates in the absence of TEV protease**. **Rho-DEVD-AOMK** (1 µM) was used to visualize proCASP10TEV by gel-based ABPP. Both batches of recombinant protein show cleaved protein (~20 kDa band), consistent with the molecular weight of activeCASP10. proCASP10, proCASP8, and activeCASP10 were prepared in PBS buffer at a final concentration of 1 µM in duplicates and treated with **Rho-DEVD-AOMK** for 1h at ambient conditions.

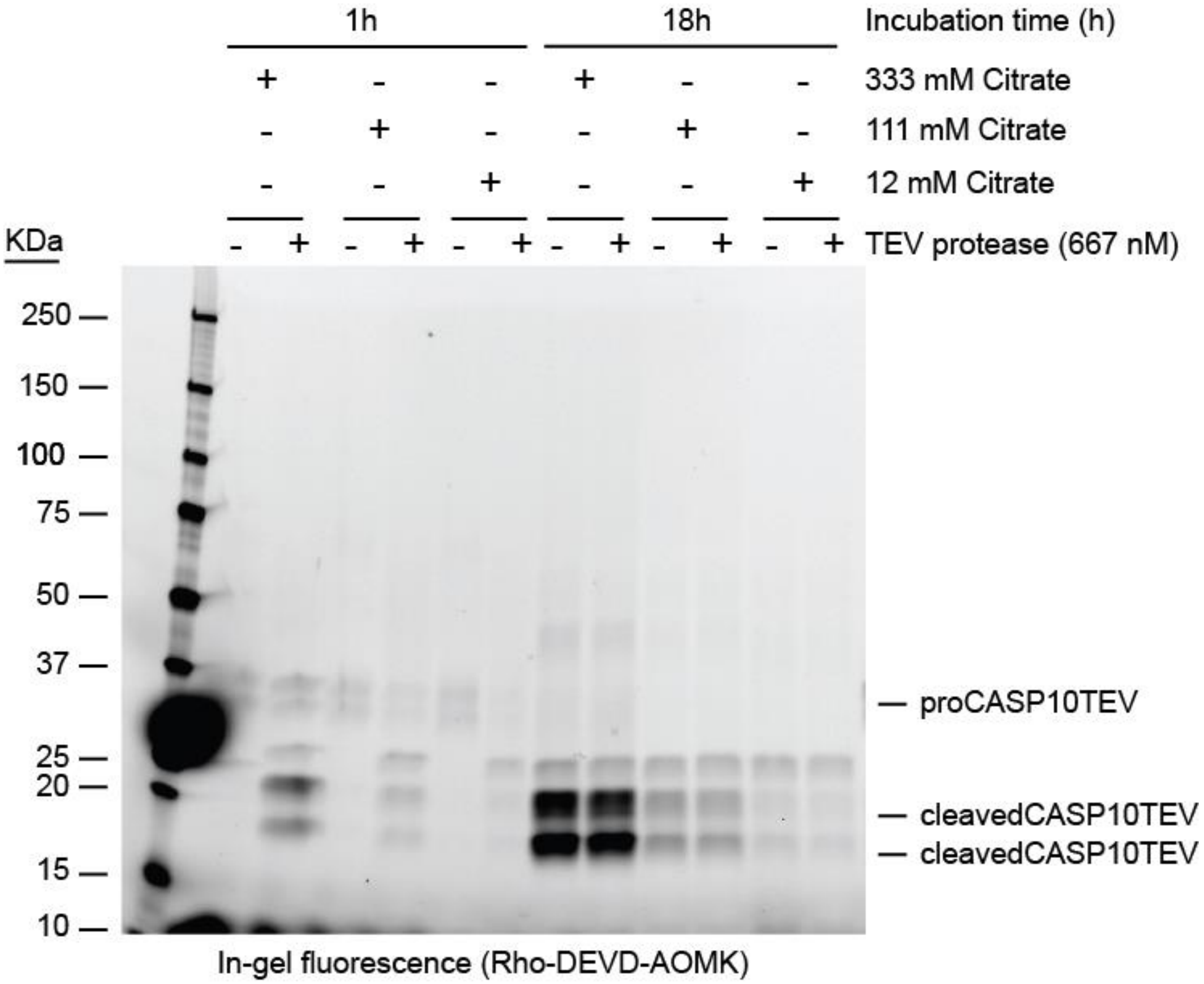


**Figure S4**. **proCASP10TEV self-activation increases over time and is enhanced by adding sodium citrate kosmotrope.** Recombinant proCASP10TEV (300 nM) was subjected to incubation in storage buffer (20 mM Tris-HCl pH 7.2, 50 mM NaCl, 1 mM DTT) for the indicated times with or without the addition of TEV protease (667 nM) and sodium citrate (pH 7.4) at the indicated concentrations (333 mM, 111 mM, and 12 mM). Protein activity was then visualized by gel-based ABPP after labeling with **Rho-DEVD-AOMK** (1 μM).

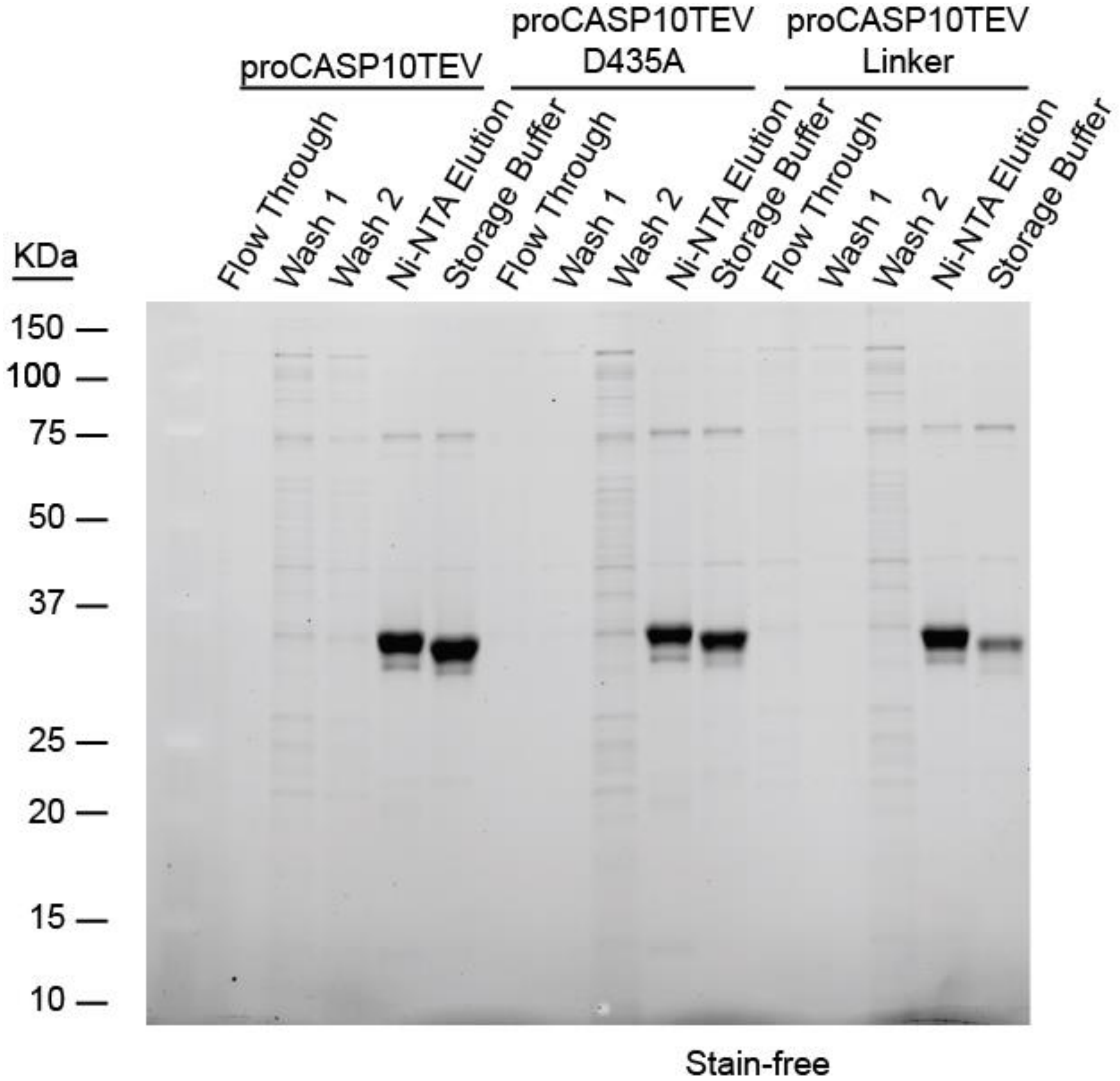


**Figure S5**. **Enhanced protein production conditions reduce caspase self-activation.** Recombinant proCASP10 construct purification stain-free gel for proCASP10TEV, proCASP10TEV D435A, proCASP10TEV linker, and proCASP10TEV delD435. Caspase storage buffer consists of 20 mM Tris pH 7.4, 50 mM NaCl, 1 mM DTT.

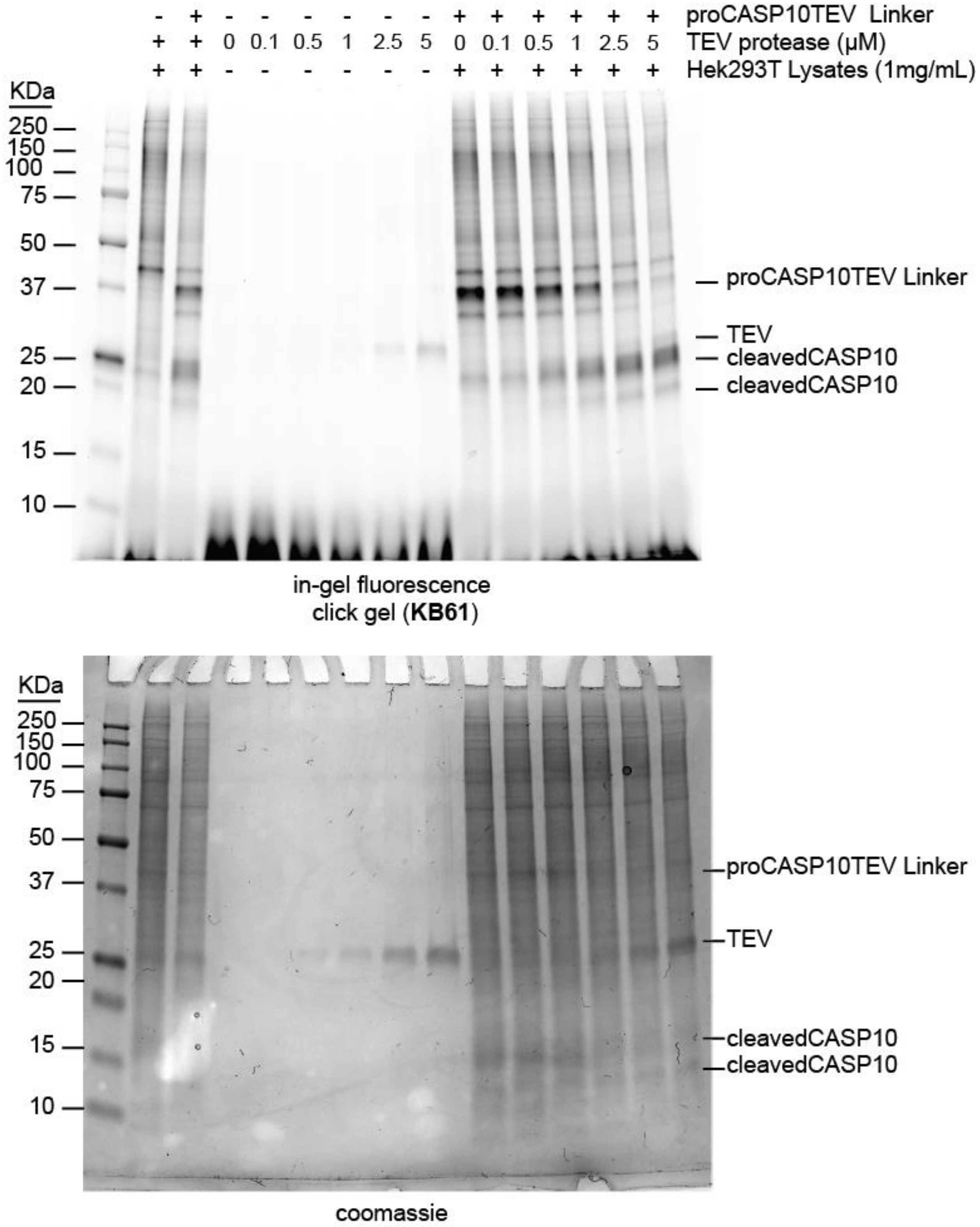


**Figure S6**. **proCASP10TEV Linker is cleaved with the addition of TEV protease in a dose-dependent manner**. 1 mg/mL HEK293T cell lysates were spiked with proCASP10TEV Linker (1 μM) followed by treatment with increasing TEV protease concentrations (0 nM, 100 nM, 500 nM, 1 μM, 2.5 μM, and 5 μM) for 1h at ambient conditions. All samples were then treated with KB61[1]

(10 µM final concentration) for 1h. Each sample was clicked with rhodamine azide click mix for 1 additional hour and visualized by in-gel fluorescence.

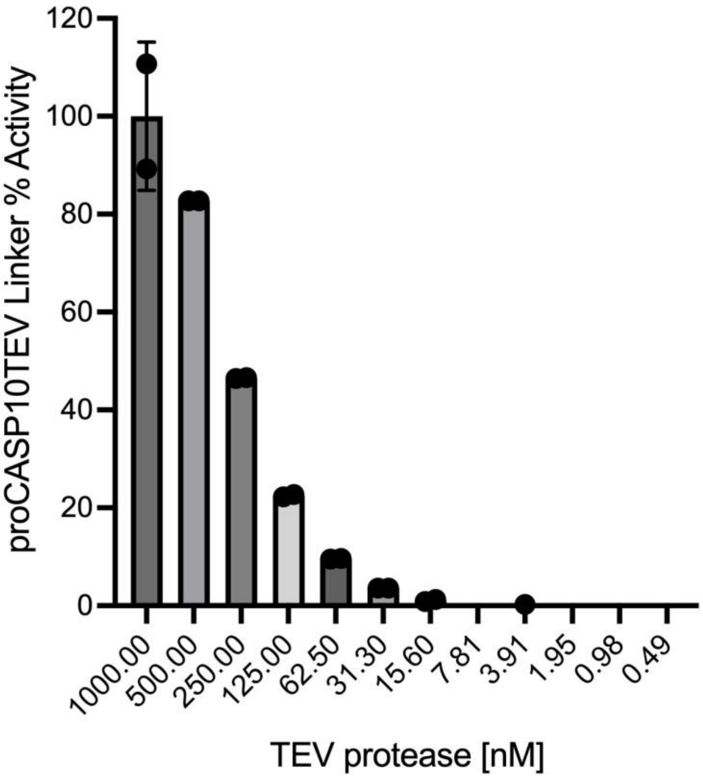


**Figure S7**. **Defining the minimal concentration of TEV protease required for screening.** Caspase substrate (Ac-VDVAD-AFC 10 µM) in citrate buffer (333 mM citrate, 5 mM DTT in PBS) was dispensed into 96-well plates containing TEV protease in PBS. TEV protease dilutions were prepared at the final concentrations: 1000 nM, 500 nM, 250 nM, 125 nM, 62.5 nM, 31.3 nM, 15.6 nM, 7.81 nM, 3.91 nM, 1.95 nM, 0.98 nM, and 0.49 nM. The activity of proCASP10TEV Linker was read immediately after mixing proCASP10TEV Linker (333 µM in PBS) with TEV protease samples. (400 nm excitation, 505 nm emission). Data represents mean values and standard deviations (n = 2 biological replicates).

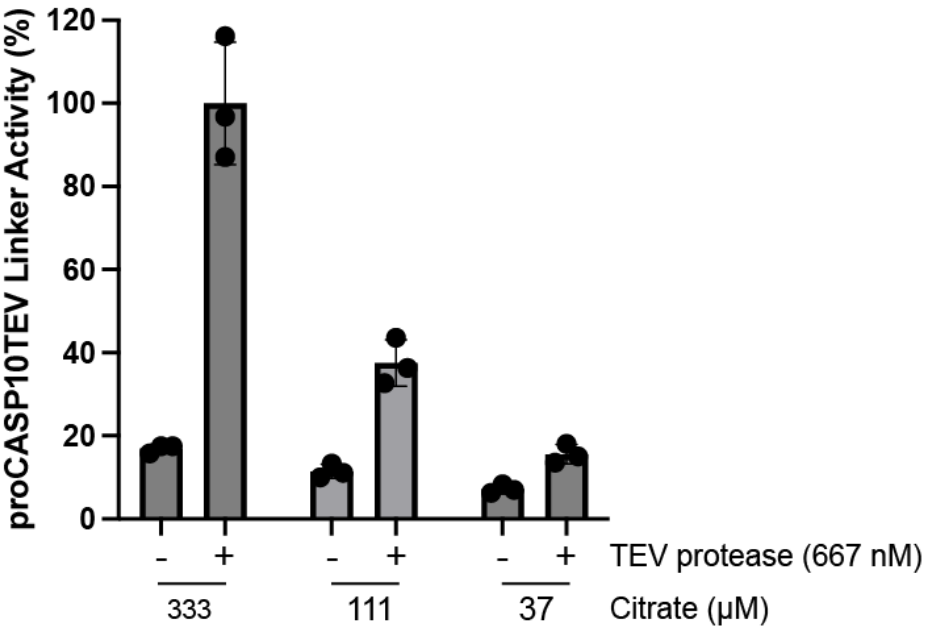


**Figure S8**. **Increasing concentration of sodium citrate enhances proCASP10TEV Linker activity in a TEV-dependent manner**. proCASP10TEV Linker (333 nM) was diluted in PBS buffer and mixed with substrate solution containing 5 mM DTT, 10 µM substrate (Ac-VDVAD-AFC), and the indicated final concentrations (333 nM, 111 nM, and 37 nM) of sodium citrate (pH 7.4) in the absence or presence of TEV protease. Data represent mean values and standard deviations (n = 2 biological replicates).

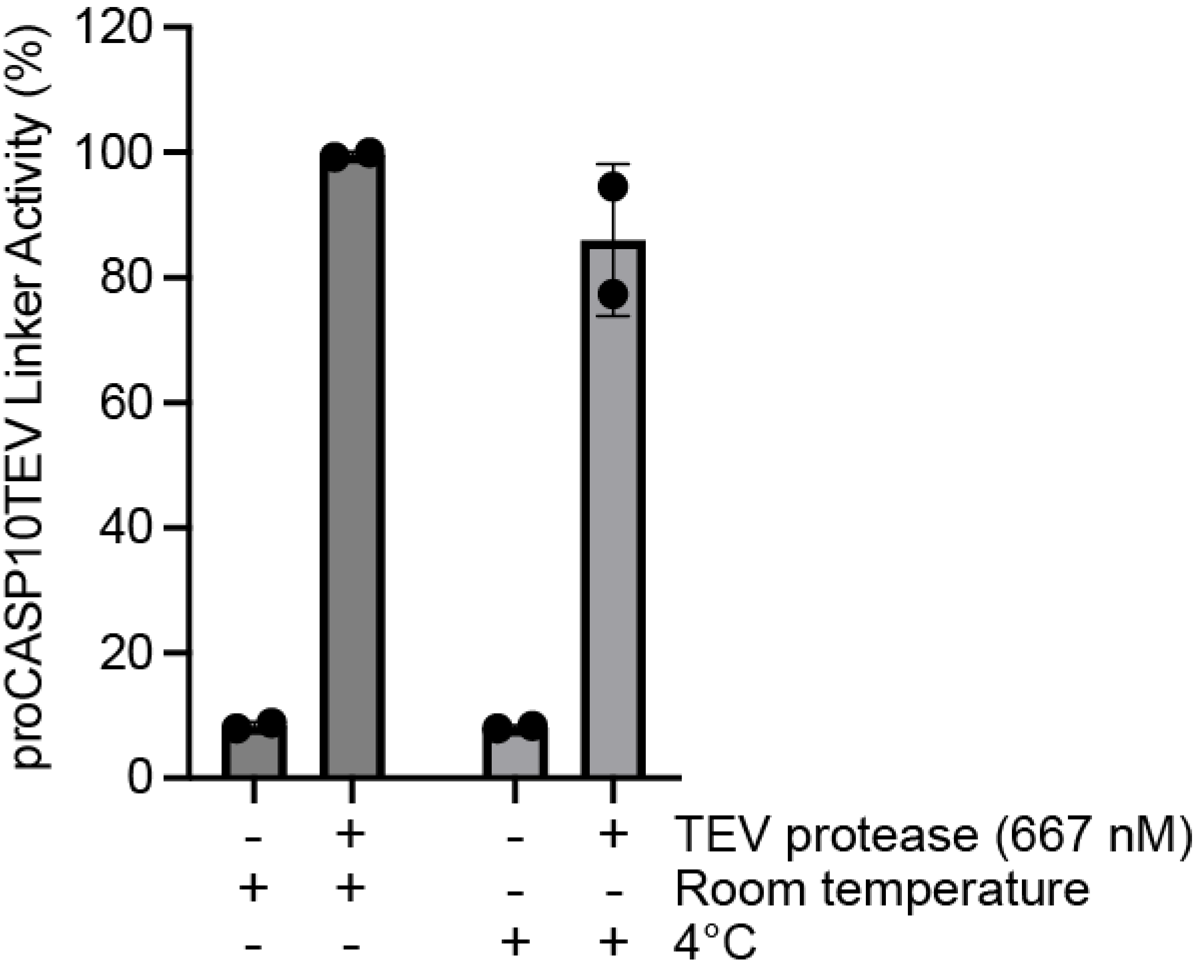


**Figure S9**. **ProCASP10TEV Linker is stable after 18h incubation in two different storage conditions (ambient, 4°C, and -80°C)**. proCASP10TEV Linker was diluted in PBS buffer at a final concentration of 333 mM and mixed with substrate solution containing 5 mM DTT, 10 µM of the caspase fluorogenic substrate, Ac-VDVAD-AFC, and 333 mM sodium citrate pH 7.4. TEV-treated and non-TEV-treated samples were also treated with the same substrate solutions containing citrate and DTT in PBS. Data represent mean values and standard deviations (n = 3 biological replicates).

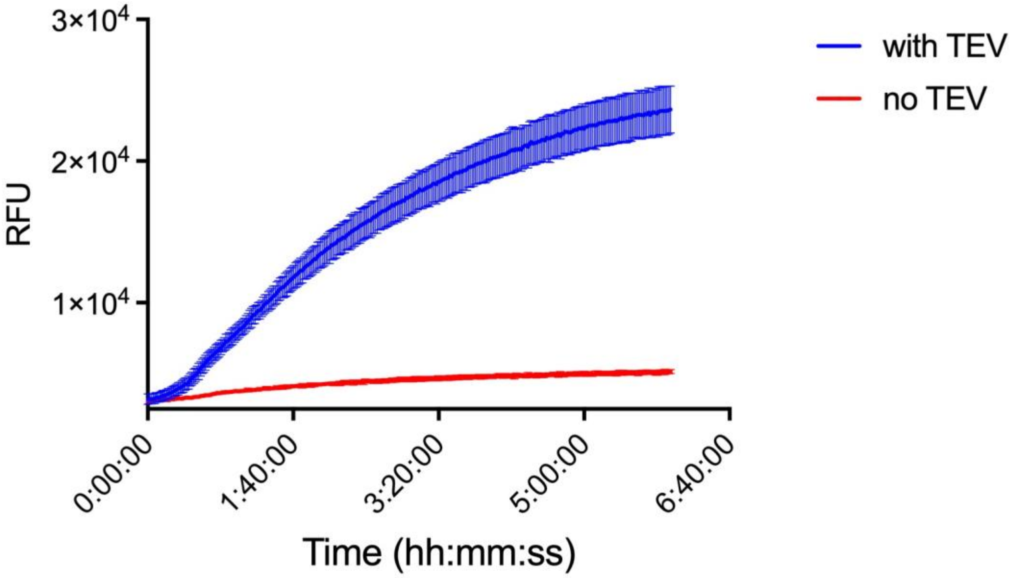


**Figure S10**. **Assaying the time-dependence of the proCASP10TEV Linker assay**. Caspase Fluorogenic substrate, Ac-VDVAD-AFC, can be cleaved by proCASP10TEV Linker and maintains activity after 6h at ambient conditions. 10 µL of proCASP10TEV Linker at 333 nM final concentration in PBS was dispensed using the Mantis liquid handler, followed by the addition of 10 µL of citrate buffer with and without TEV (333 mM sodium citrate, 5 mM DTT, 10 µM substrate, and 667 nM TEV protease in PBS). Data represent mean values and standard deviations (n = 5 biological replicates).

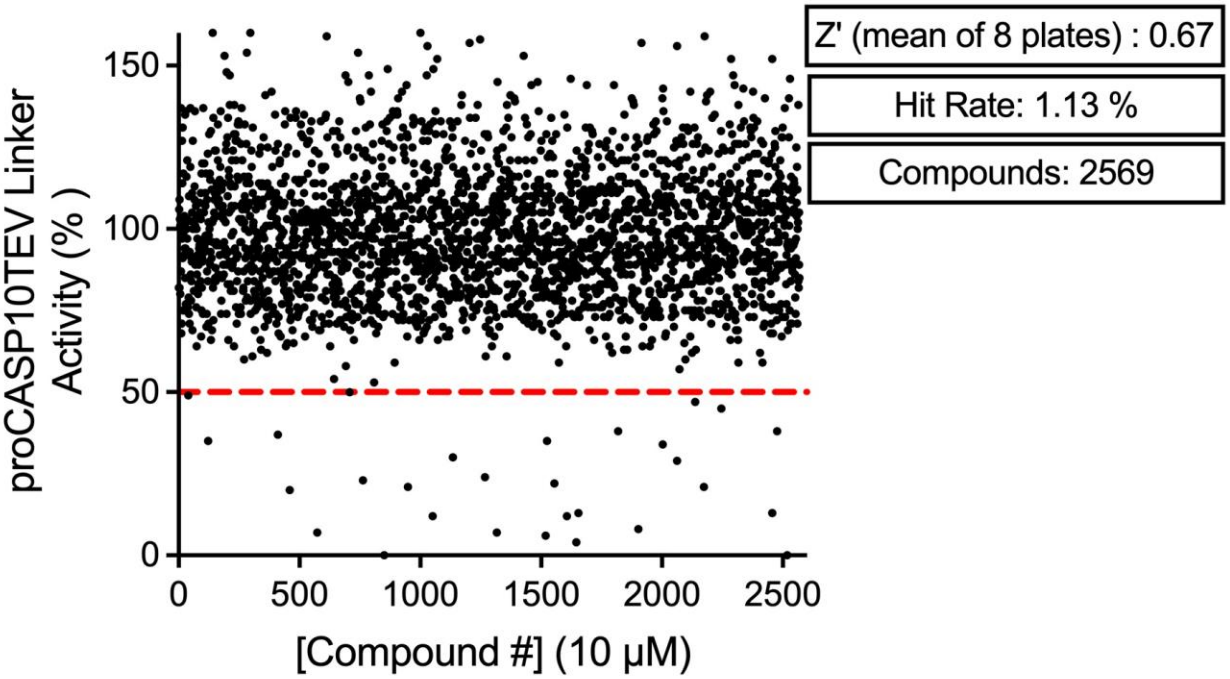


**Figure S11. Initial manual screening of proCASP10TEV Linker with LOPAC and NPW library identified 29 compounds (1.13% hit rate) with less than 50% proCASP10TEV Linker activity**.

A total of eight low-volume 384-well plates were prepared by dispensing 10 µL of proCASP10TEV Linker (333 nM) in PBS using the Mantis Liquid Dispenser (Formulatrix). Plates were manually pinned with compounds (Beckman Coulter Biomek FX system) from two different FDA-approved libraries and left to incubate for 1h at ambient conditions. In PBS, 10 µL of TEV solution was dispensed onto each plate containing 667 nM TEV protease, 333 mM citrate pH 7.2, 5 mM DTT, and 10 µM substrate (Ac-VDVAD-AFC). Plates were left to incubate for 1h. Endpoint reads were obtained for each well and percent activity was calculated to the average of DMSO positive controls (n = 16).

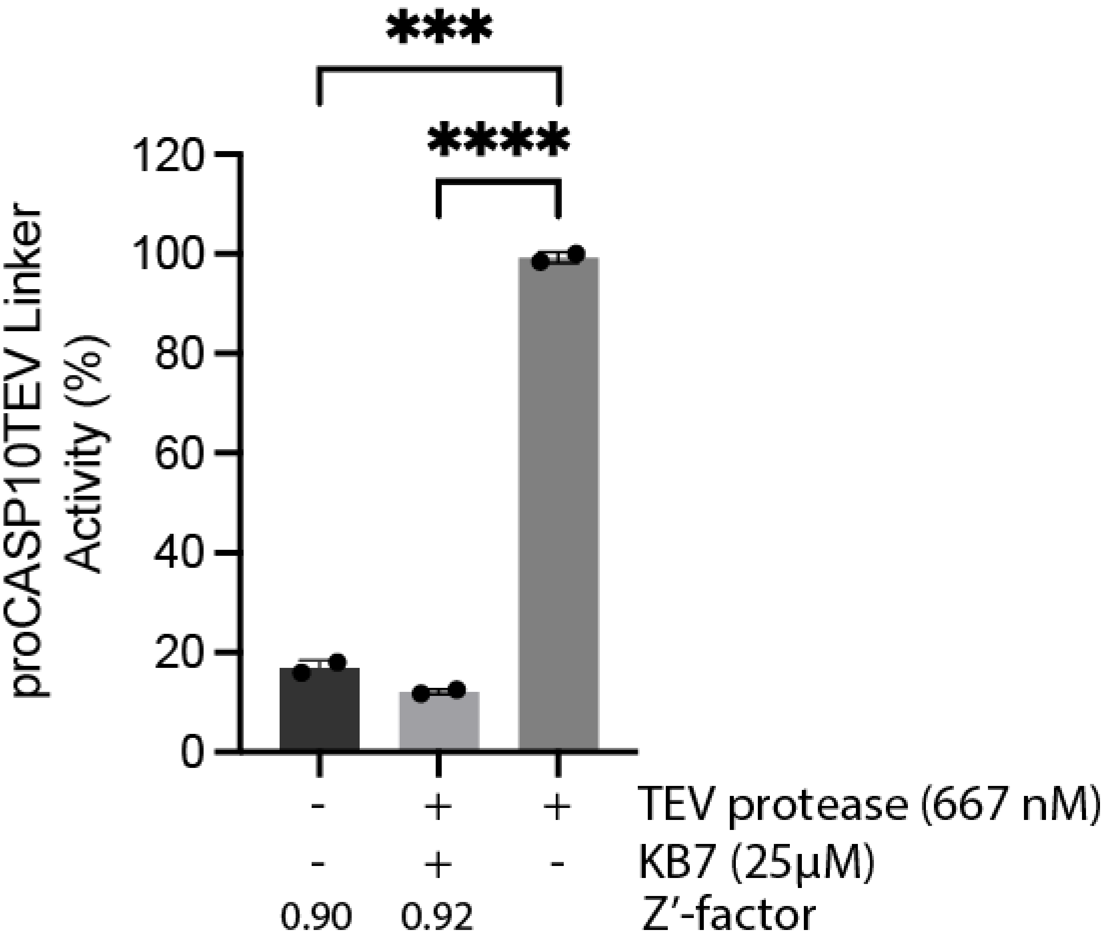


**Figure S12**. **Control plate confirms compatibility of proCASP10TEV Linker with HTS**. Z' for the no TEV protease control of 0.90 and the positive control, KB7 (25 µM), of 0.92 supports the compatibility of the proCASP10TEV Linker with high throughput screening. proCAPS10TEV Linker at 333 nM with citrate buffer in the absence of TEV protease was treated with KB7 for 1h followed by adding citrate buffer containing TEV protease at 667 nM and 333 mM sodium citrate. Data represent mean values and standard deviations (n = 2 biological replicates). Statistical significance was calculated with unpaired Student's t-tests, ****p<0.0001.

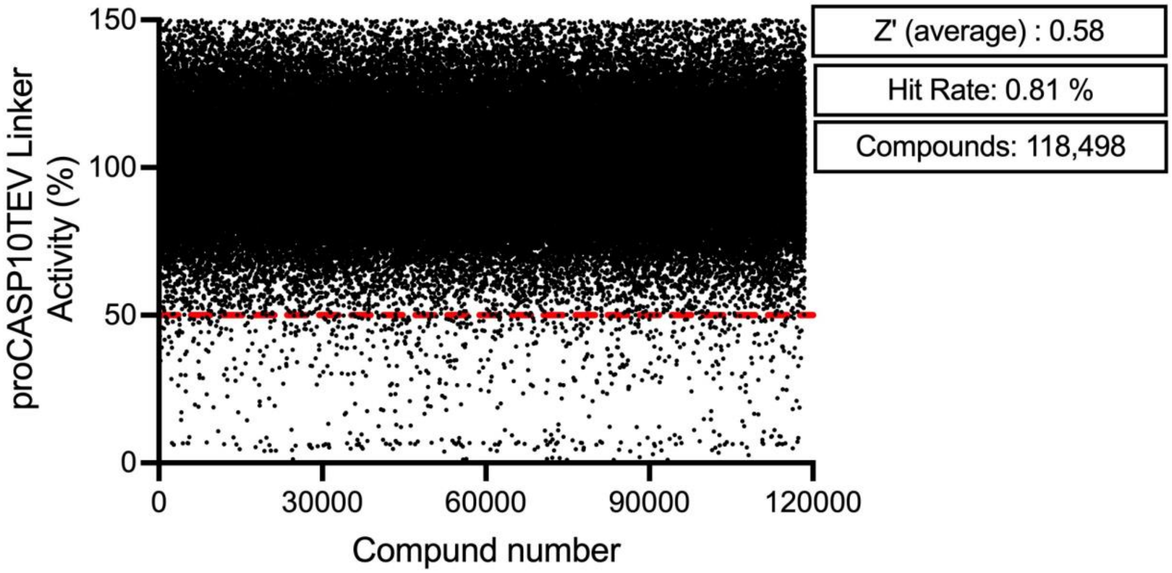


**Figure S13**. **A total of 963 initial hit compounds were found to have less than 50% proCASP10TEV Linker activity**. A total of 118,498 compounds were analyzed to determine whether they were hits based on proCASP10TEV Linker activity (0.81% hit rate). Endpoint values from each well of the 384-well plates (375 plates) were used to calculate the percent activity relative to the average of the DMSO treatments.

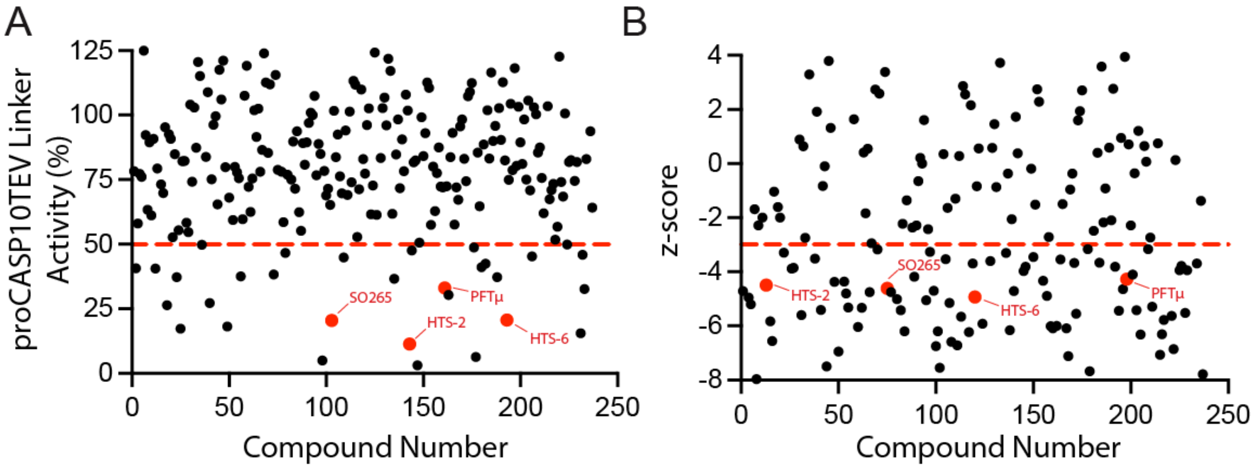


**Figure S14**. **Re-screen of proCASP10TEV Linker validated hit compounds**. A) Percent activity of proCASP10TEV Linker re-screen results showing approximately 32 of 237 compounds having ≤ 50% activity (red horizontal line). Percent activity of LOPAC hit compounds are represented as red dots (compound number 143 for **HTS-2**, compound number 193 for **HTS-6,** and compound number 163 for **PFTμ**) B) z-score of proCASP10TEV Linker re-screen results showing 38 compounds with a z-score ≤ -3 (red horizontal line) calculated using the mean of the control lanes. The z-score of LOPAC hit compounds are represented as red dots (compound number 13 for **HTS-2**, compound number 120 for **HTS-6,** and compound 198 for **PFTμ**).

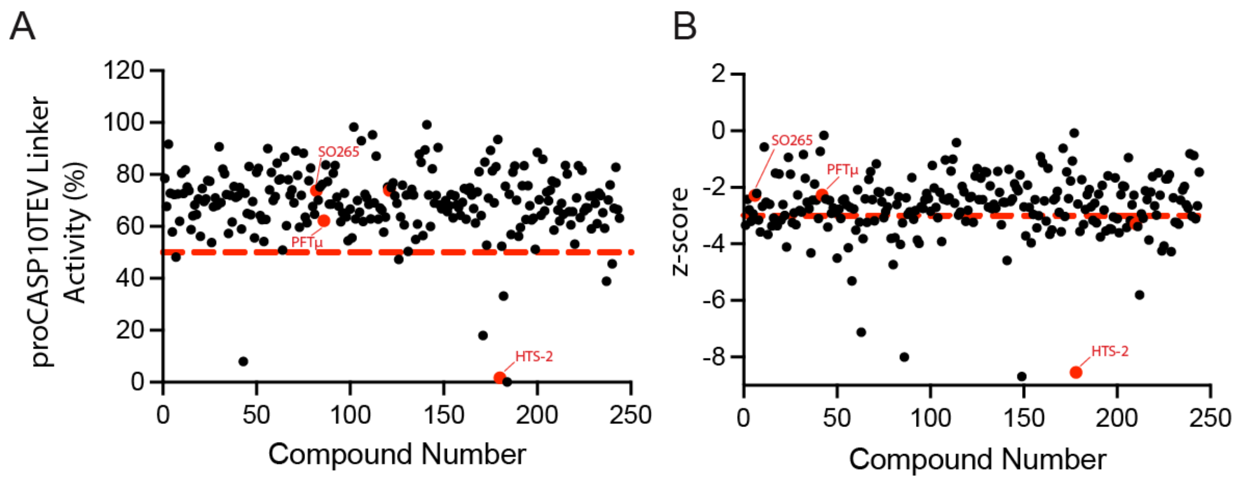


**Figure S15**. **Caspase-10 counter-screen identified inhibitors for the active form (activeCASP10)**. A) Percent activity of activeCAP10 counter screen results showing 40 of 237 compounds having ≤ 60% activity (red horizontal line). B) z-score of activeCASP10 counter screen results showing 78 compounds with a z-score of ≤ -3 (red horizontal line) calculated using the mean and standard deviation of the DMSO wells (n = 12). The z-score of LOPAC hit compounds is represented as red dots.

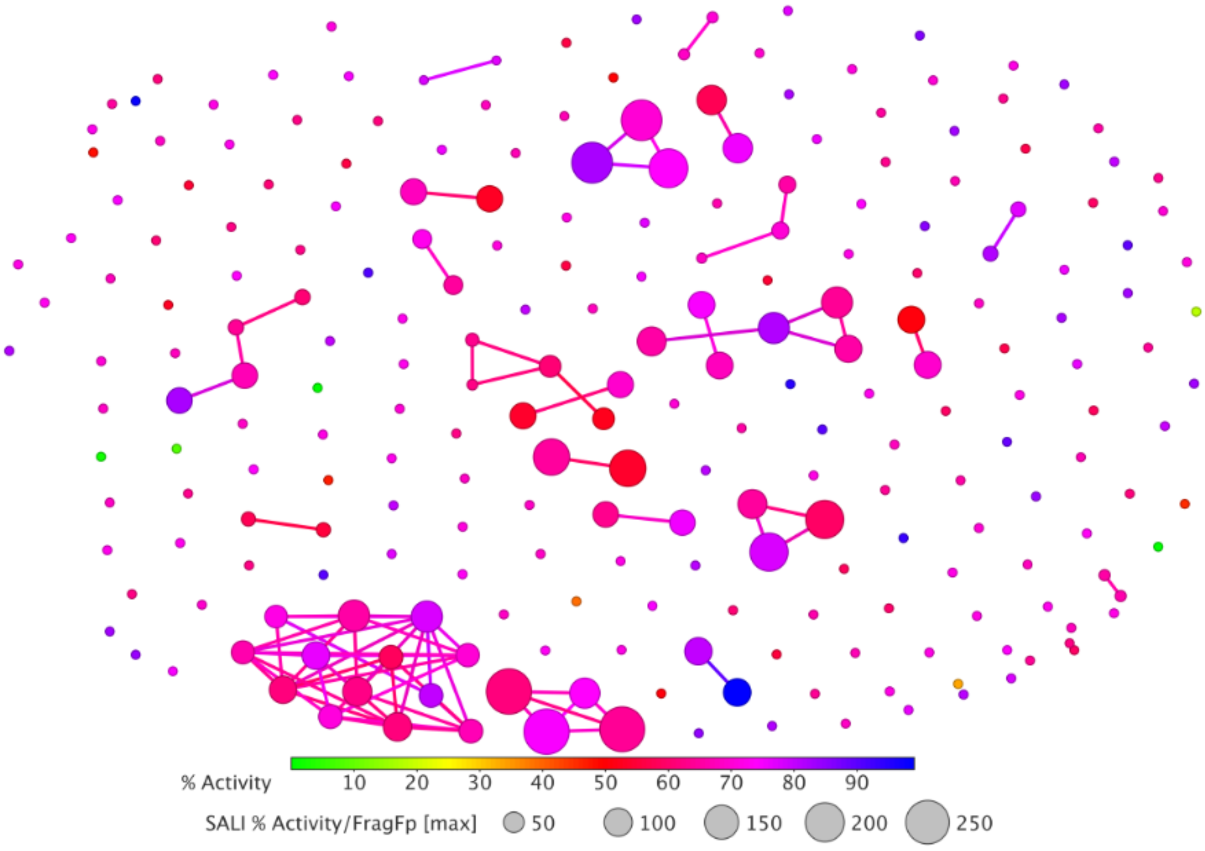

**Figure S16**. **SAR analysis of activeCASP10 counter screen with 19 clusters.** Analysis was performed as described in the DataWarrior SAR analysis methods section. The structure-activity landscape index (SALI) was first calculated using the percent activities based on the DMSO controls (n = 16) [2].

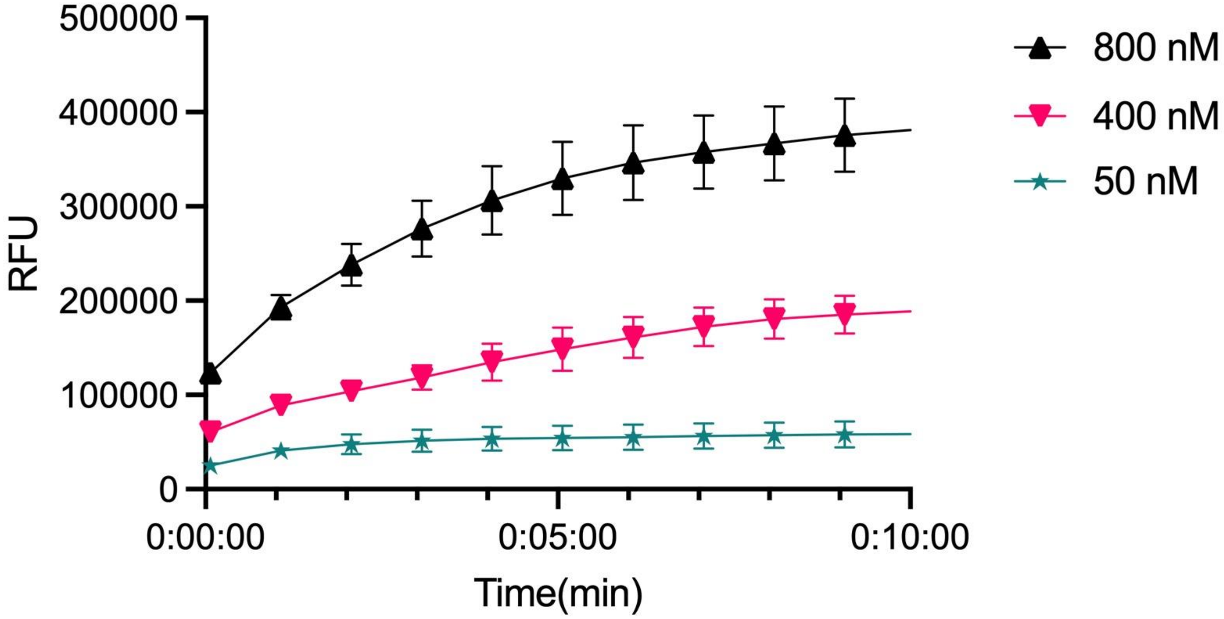


**Figure S17**. **Determining optimal concentration of TEV protease substrate to validate screen hits**. **ABCYL-ENLYFQSGTK-5-FAM** substrate was added to the citrate buffer containing 333 mM sodium citrate and 5 mM DTT in PBS. TEV protease (100 nM) in PBS was then mixed with the citrate buffer at a 1:1 mixture (100 µL of TEV protease with 100 µL of citrate buffer) and immediately analyzed on a plate reader. Data represents mean values and standard deviations (n = 4 biological replicates).

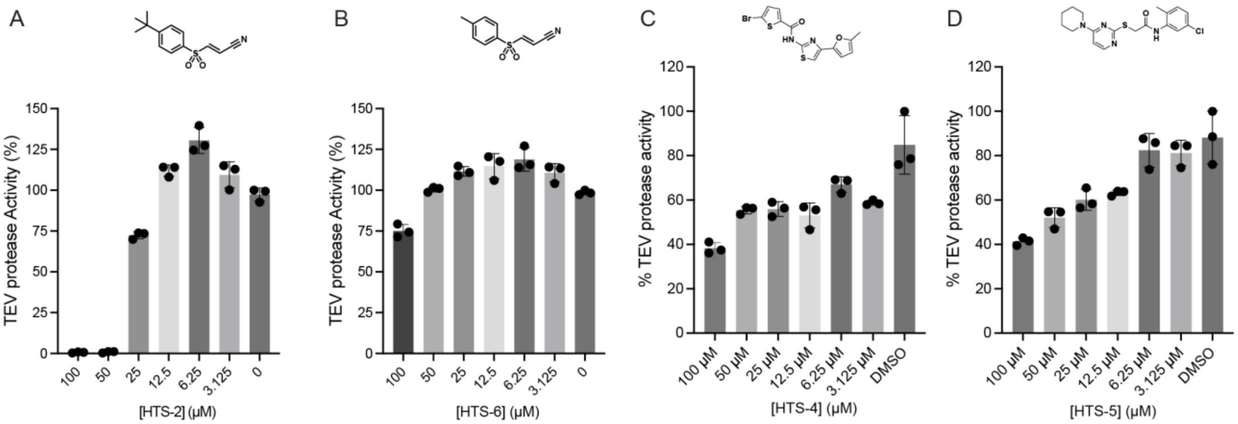


**Figure S18**. **proCASP10TEV Linker hit compounds have little to no effect on TEV protease activity at increasing concentration of compounds**. TEV protease at 100 nM was treated with (A) **HTS-2**, (B) **HTS-6**, (C) **HTS-4**, and (D) **HTS-5** at indicated concentrations for 1h followed by the addition of TEV substrate solution. Data represents mean values and standard deviations (n = 3 biological replicates).

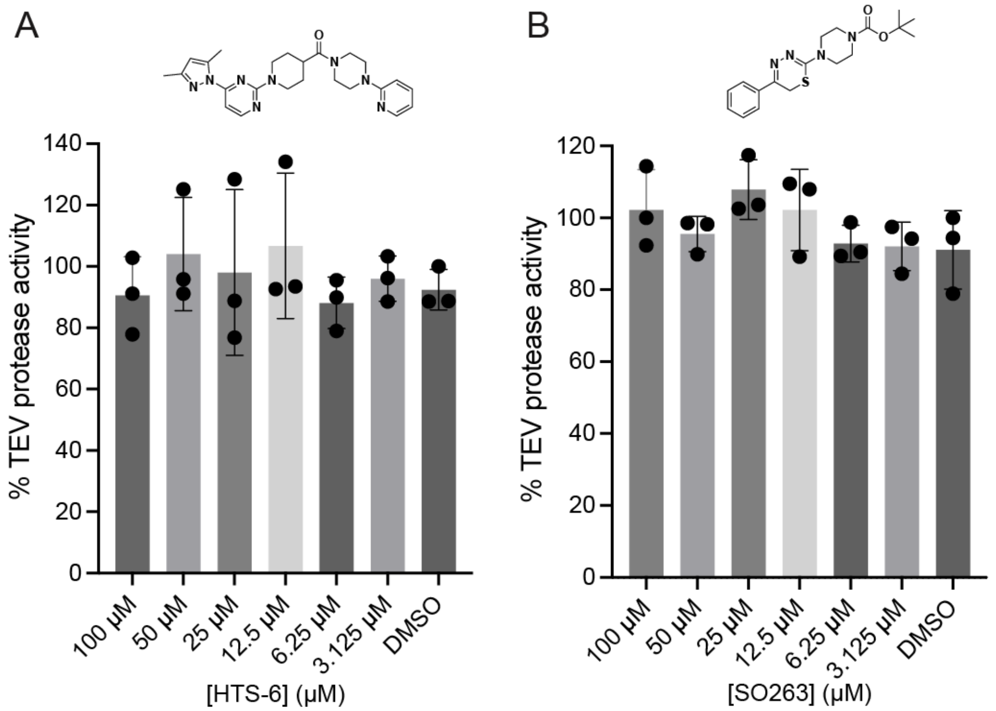


**Figure S19**. **Initial screen compounds can inhibit TEV protease at higher concentrations**. TEV protease (100 nM) was treated with (A) **HTS-8** and (B) **SO263** in DMSO at indicated final concentrations for 1h followed by the addition of TEV substrate solution. Data represents mean values and standard deviations (n = 3 biological replicates).

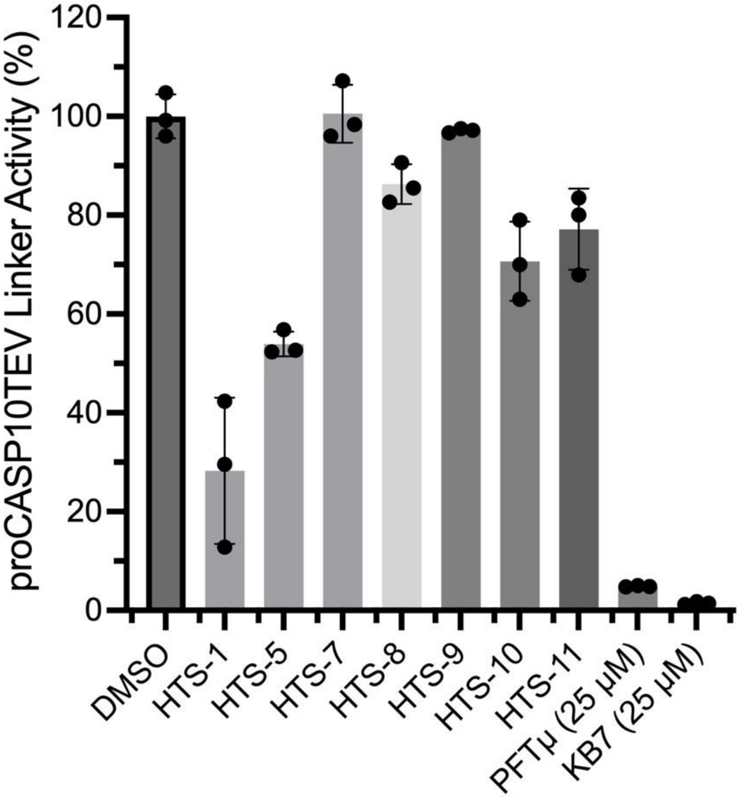


**Figure S20**. **Validation of proCASP10TEV Linker re-screen results**. proCASP10TEV Linker at 333 nM final concentration was treated with screen compounds at 100 µM, **KB7** positive control at 25 µM and **PFTµ** at 25 µM for 1h at room temperature. Data represents mean values and standard deviations (n = 3 biological replicates).

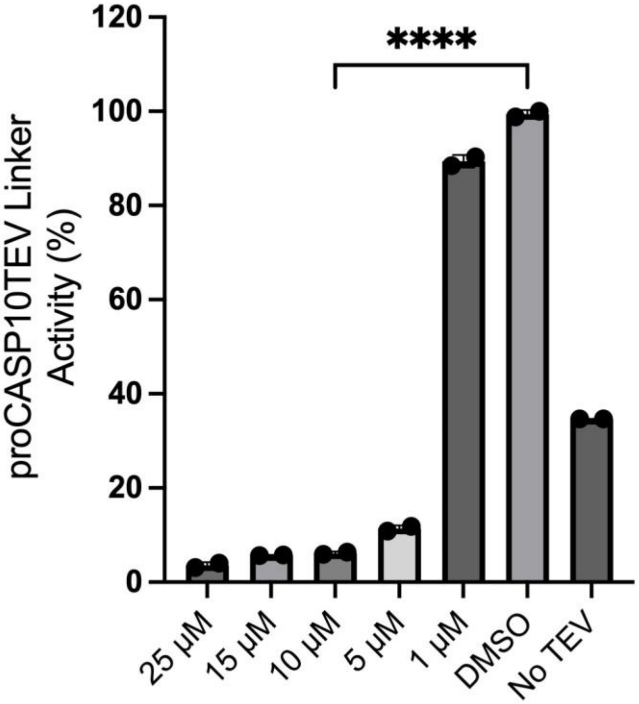


**Figure S21. procaspase-10 TEV Linker is inhibited by the highly reactive screen hit HTS-6 in phosphate buffer**. proCASP10TEV Linker in PBS (333 nM) was treated with **HTS-6** for 1h at decreasing concentrations (25 µM, 15 µM, 10 µM, 5 µM and 1 µM). Data represents mean values and standard deviations (n = 2 biological replicates). Statistical significance was calculated with unpaired Student's t-tests, ****p<0.0001.

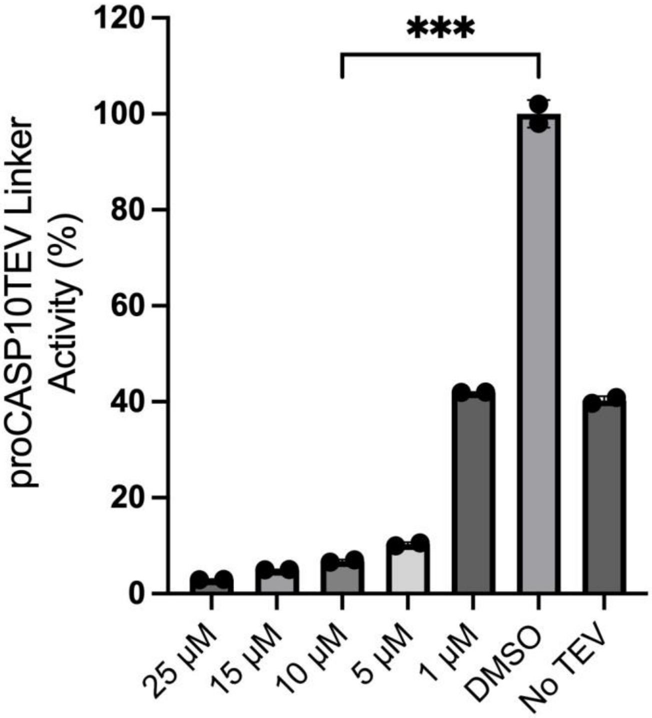


**Figure S22**. **procaspase-10 TEV Linker is inhibited by the highly reactive screen hit HTS-2 in phosphate buffer**. Data represents mean values and standard deviations (n = 2 biological replicates). Statistical significance was calculated with unpaired Student's t-tests, ***$p<0.001$.

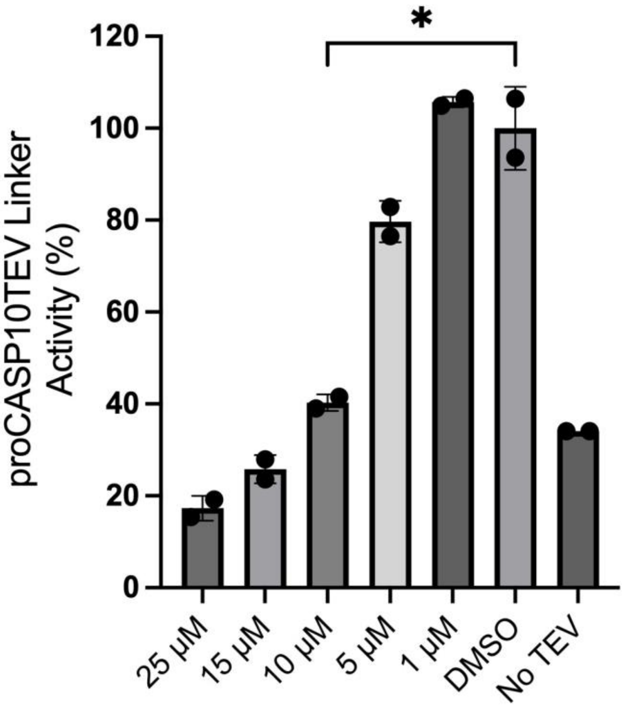


**Figure S23. procaspase-10 TEV Linker is inhibited by the highly reactive screen hit PFTµ in phosphate buffer.** proCASP10TEV Linker (333 nM) spiked in PBS was treated with **PFTµ** at varying concentrations (25, 15, 10, 5, and 1 µM) for 1h. Data represents mean values and standard deviations (n = 2 biological replicates). Statistical significance was calculated with unpaired Student's t-tests, ***p<0.001.

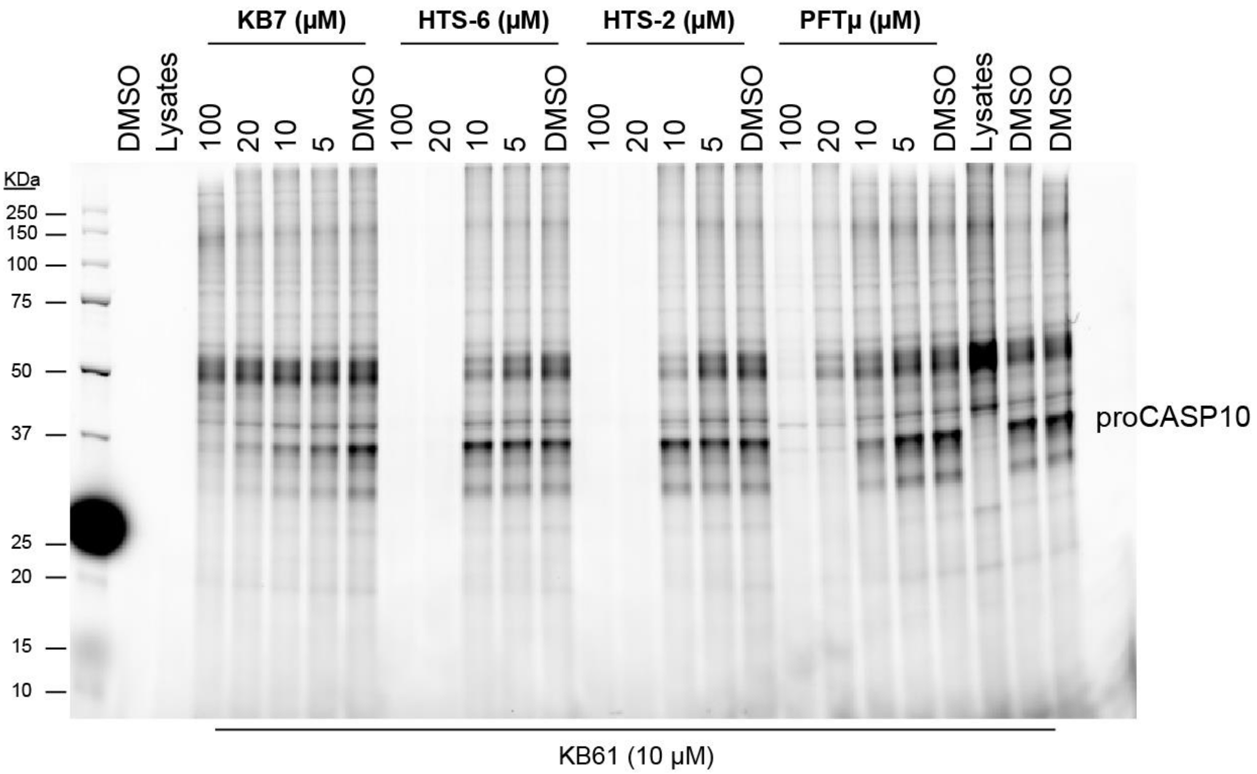

**Figures S24**. **Initial screen compounds can compete for procaspase-10 labeling against KB61**. 1 mg/mL HEK293T cell lysates were spiked with 1 µM proCASP10 and treated with LOPAC compounds at decreasing final concentrations (100 µM, 20 µM, 10 µM, and 5 µM) for 1h. Samples were then treated with 10 µM **KB61** for 1h, followed by clicking onto rhodamine azide (25 µM) for 1h. Protein labeling was then visualized by competitive gel-based ABPP and visualized by in-gel fluorescence.

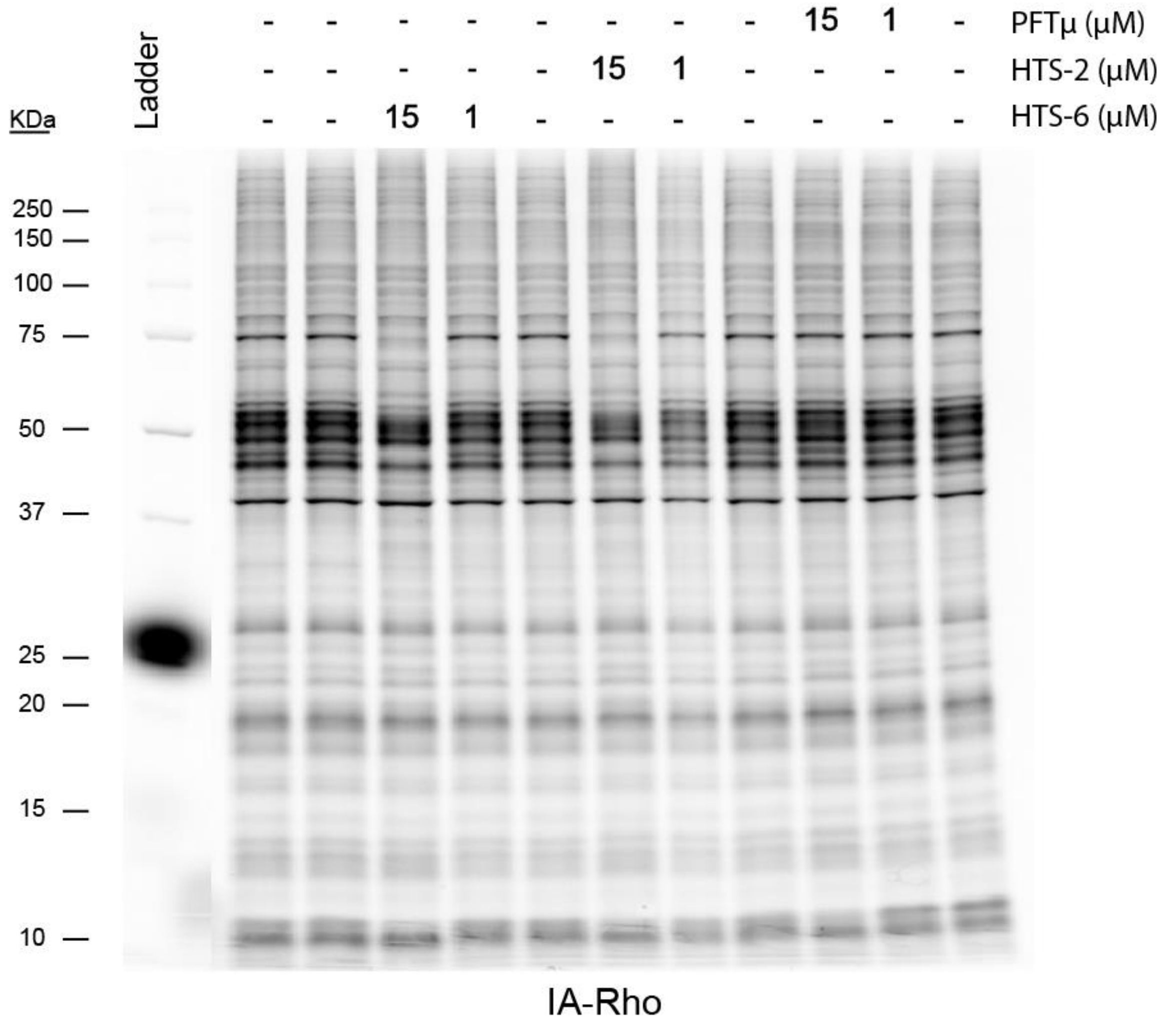


**Figure S25. Initial hit screen compounds are promiscuous**. 1 mg/mL HEK293T lysates were treated with LOPAC compounds at 15 µM and 1 µM concentrations for 1h followed by treatment with 1 µM IA-Rho for 1h under ambient conditions.

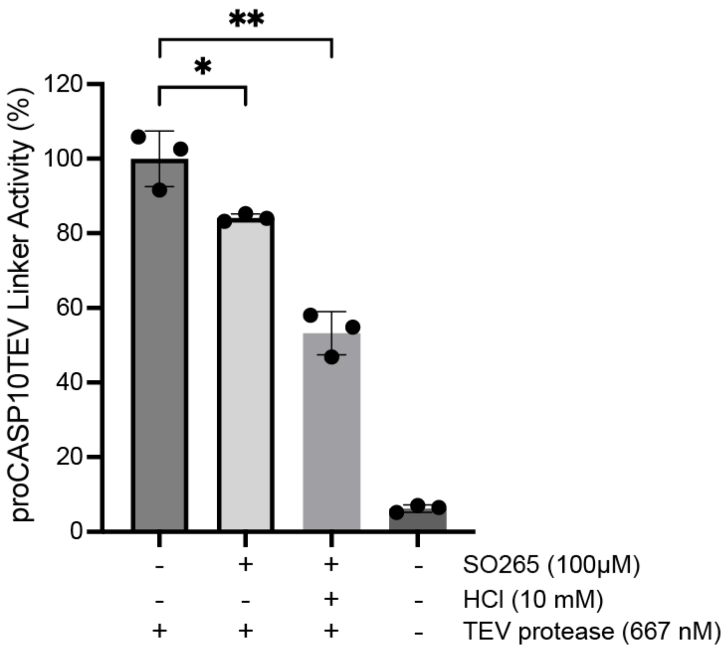


**Figure S26. Compound SO265 activates under acidic conditions, decreasing proCASP2TEV Linker activity.** Compound **SO265** was prepared as a 10 mM stock in DMSO, 333 mM sodium citrate, or 10 mM HCl and left to incubate at 37oC for 15 min. proCASP10TEV Linker (333 nM in PBS) was treated with 10 mM stocks (concentration of 100 µM) for 1 h. Samples were then treated with TEV protease and substrate solution (10 µM substrate, 667 nM TEV protease in PBS). Data represents mean values and standard deviations (n = 4 biological replicates). Statistical significance was calculated with unpaired Student's t-tests, **p<0.001, *p<0.05.

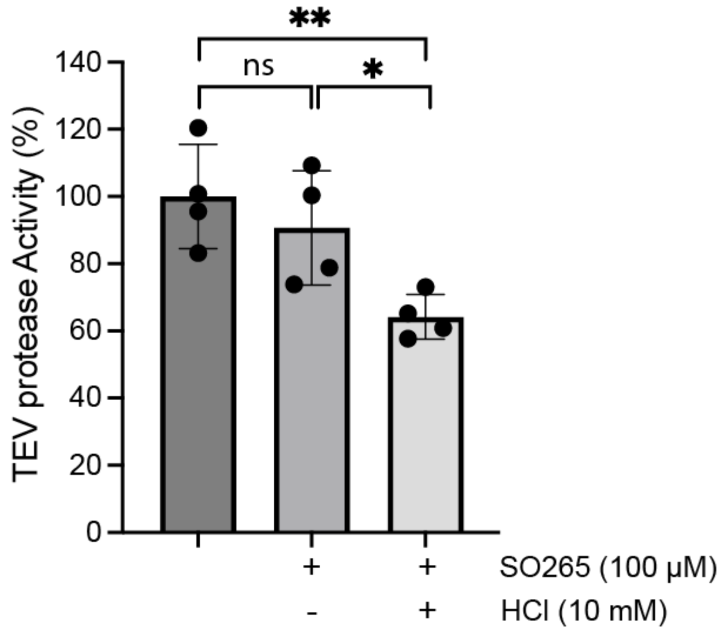


**Figure S27**. **SO265 becomes active under acidic conditions and slightly reduces TEV protease activity**. TEV protease treated with **SO265** stocks (100 uM) prepared under acidic conditions (stocks prepared in 10 mM HCl at 37°C for 15 min). Data represents mean values and standard deviations (n = 4 biological replicates). Statistical significance was calculated with unpaired Student's t-tests, *p<0.05, ns = not significant.

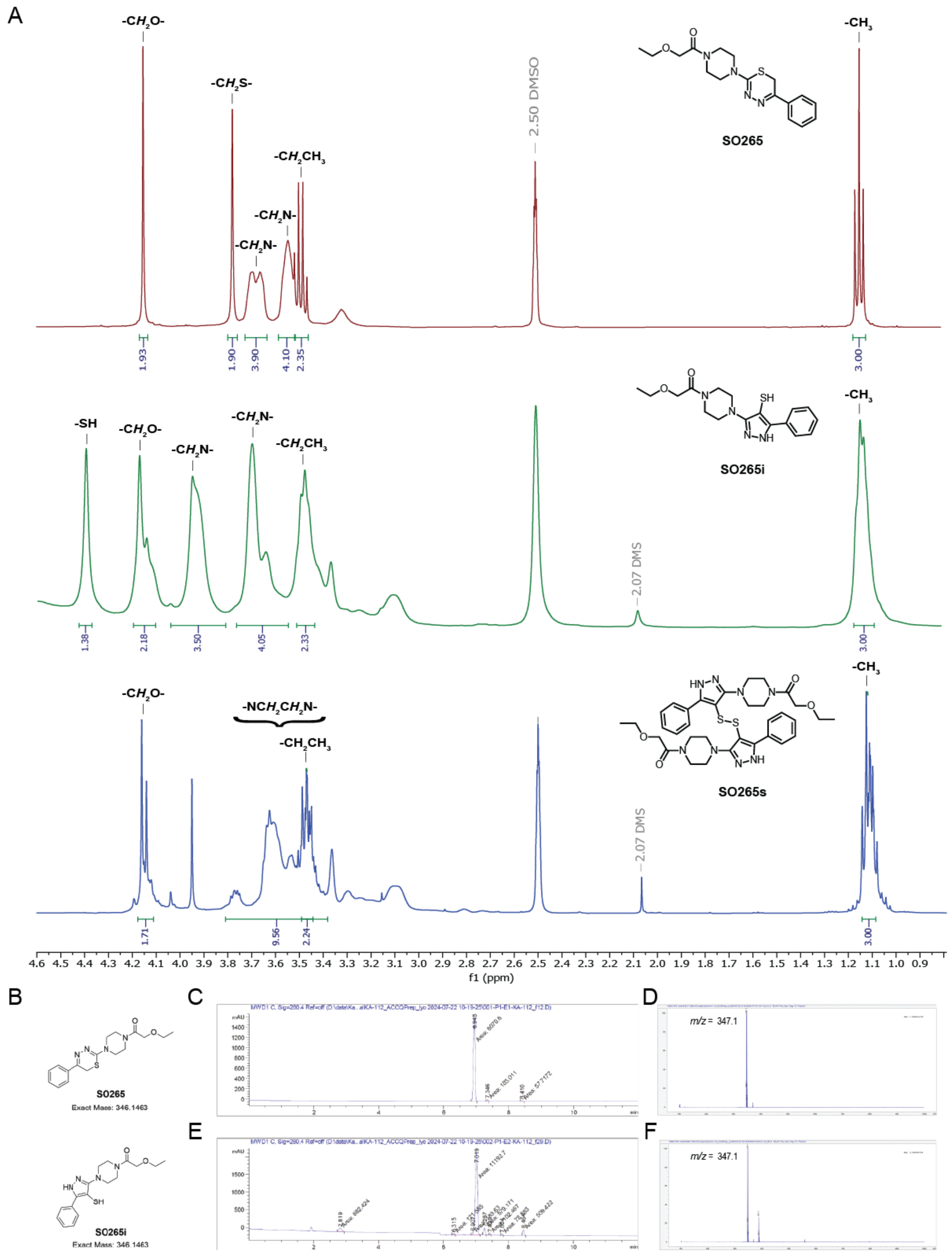


**Figure S28**. **Isomer SO265 shows a slight retention time shift in LC-MS compared to SO265**. (A) H NMR comparison of purified **SO265** compounds including **SO265** (red), the isomer **SO265i** (green), and the disulfide rearrange product **SO265s** (blue) (B) Structures of **SO265** and structural isomer **SO265i**. (C) LC trace of **SO265** (λ = 280 nm) with compound retention time = 6.945 min. (D) Mass spectrum (ESI+) for the **SO265** peak at 6.945 min showing *m/z* = 347.1 (**SO265** monoisotopic mass = 346.1). (E) LC trace of **SO265i** (λ = 280 nm) with compound retention time = 7.019 min. (F) Mass spectrum (ESI+) for the **SO265i** peak at 7.019 min showing *m/z* = 347.1 (**SO265i** monoisotopic mass = 346.1).

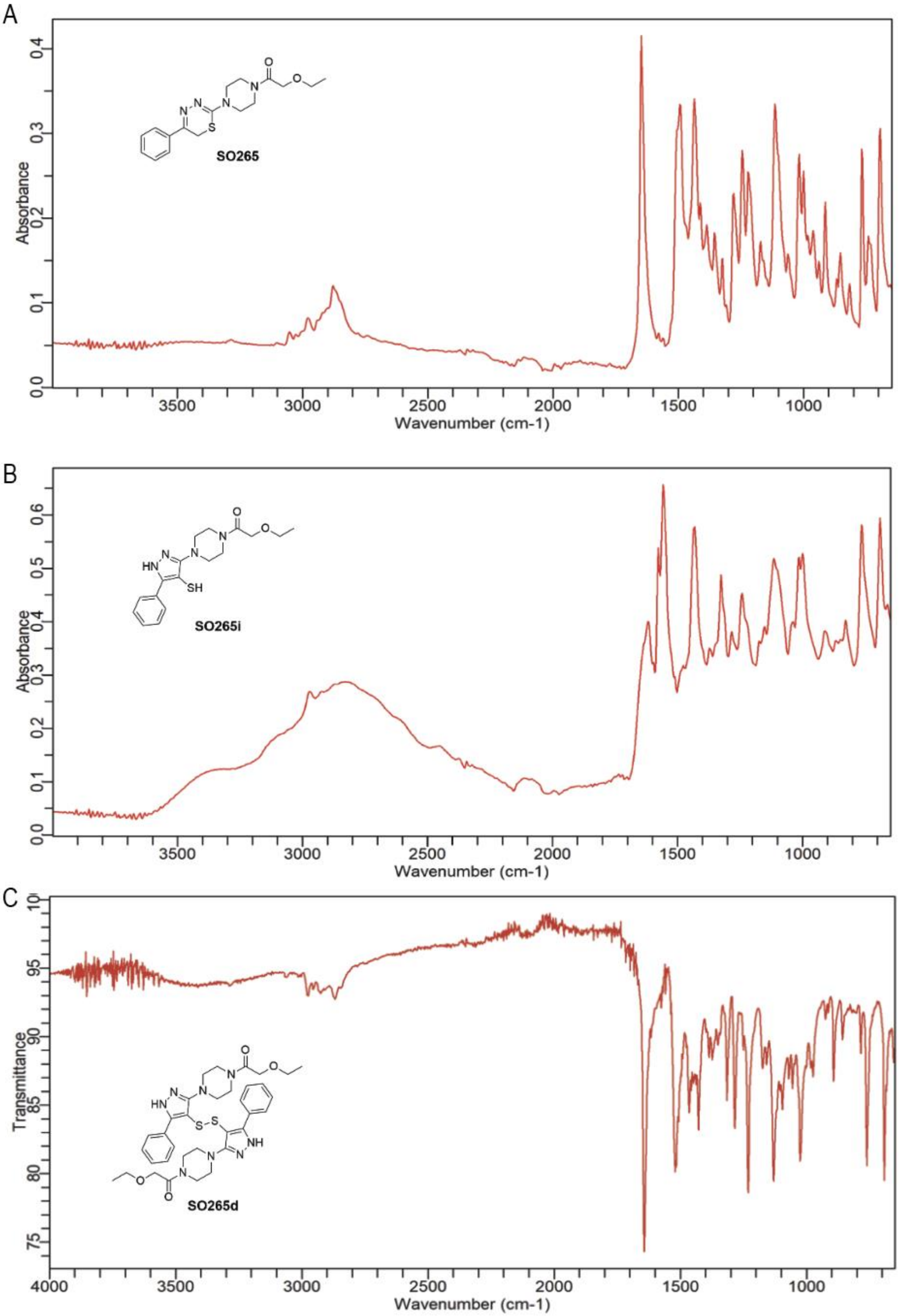
A
SO265
Absorbance
Wavenumber (cm-1)
B
SO265i
Absorbance
Wavenumber (cm-1)
C
SO265d
Transmittance
Wavenumber (cm-1)

**Figure S29. Purified SO265, purified isomer SO265i, and proposed disulfide SO265s exhibit different banding in IR spectroscopy.** (A) IR spectrum of **SO265**. (B) IR spectrum of **SO265i**. (C) IR spectrum of the **SO265i** NMR sample which had been left at room temperature for 3 days resulting in the formation of **SO265s**.

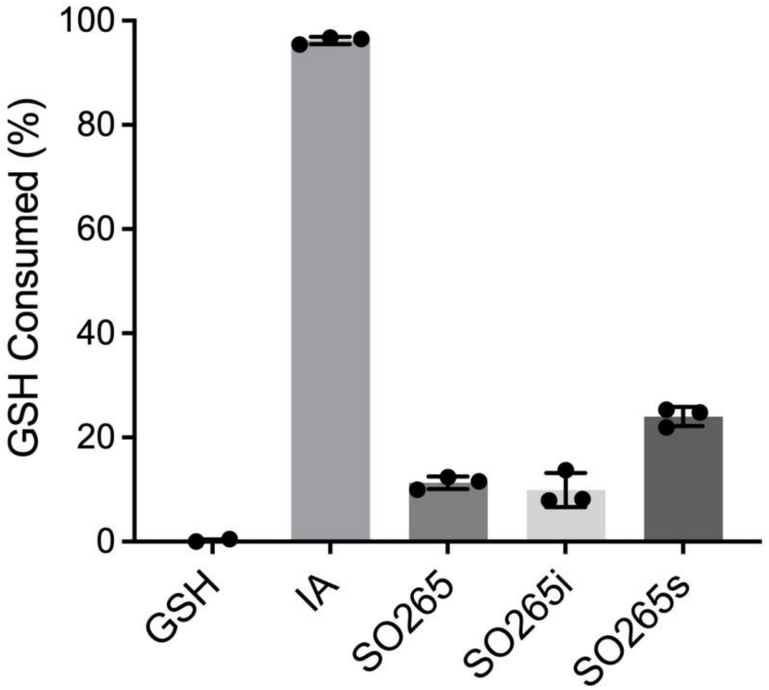


**Figure S30. The rearranged disulfide compound (SO265s) has higher activity with glutathione.** Consumed glutathione (GSH) was measured using Ellman's reagent and tested against SO265 compounds, and iodoacetamide (IA). Data represents mean values and standard deviations (n = 3 biological replicates).

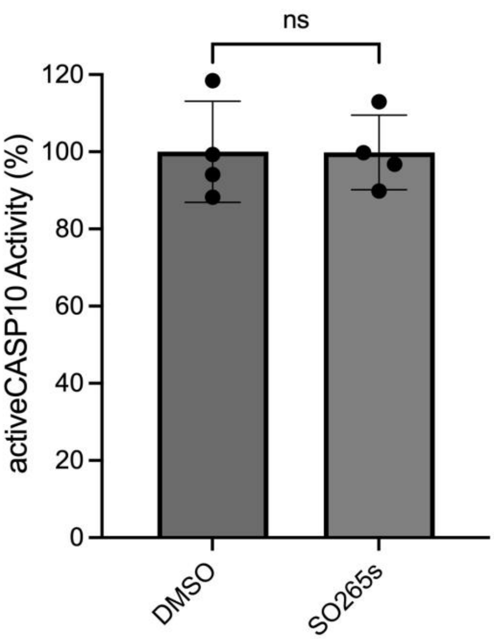


**Figure S31. ActiveCASP10 is not inhibited by SO265s**. ActiveCASP10 at 1 µM in PBS was treated with **SO265s** at final concentration of 100 µM for 1h at ambient conditions. Data represents mean values and standard deviations (n = 4 biological replicates). Statistical significance was calculated with unpaired Student's t-tests, ns = not significant.

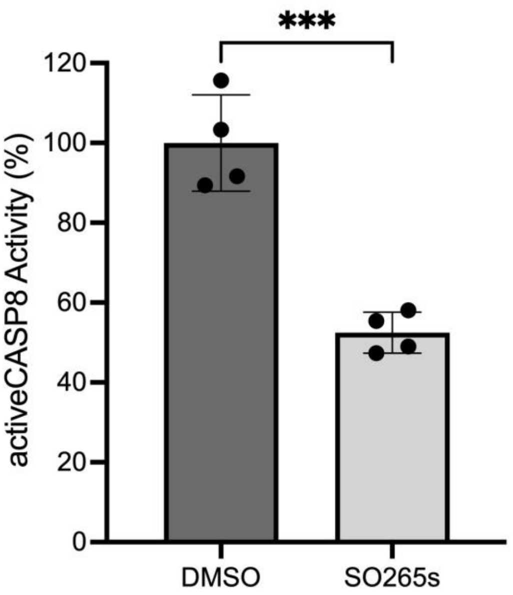


**Figure S32. ActiveCASP8 is partially inhibited by SO265s**. ActiveCASP8 at 1µM in PBS was treated with 100 µM **SO265s** for 1h at ambient conditions. Data represents mean values and standard deviations (n = 4 biological replicates). Statistical significance was calculated with unpaired Student's t-tests, ***p<0.001.

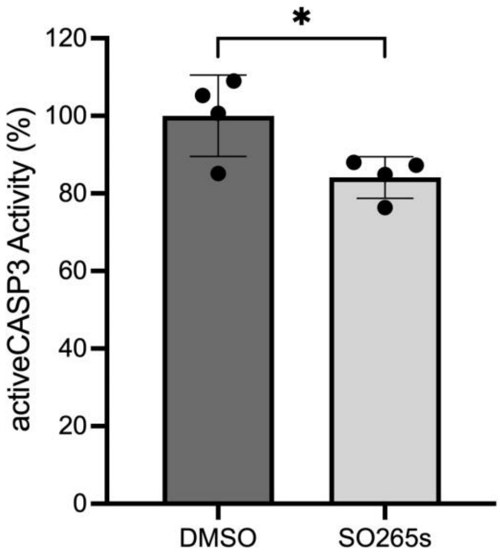


**Figure S33. ActiveCASP3 is partially inhibited by SO265s**. Active enzyme (0.1 µM) in PBS was treated with 100 µM of **SO265s** for 1h at ambient conditions. Data represents mean values and standard deviations (n = 4 biological replicates). Statistical significance was calculated with unpaired Student's t-tests, *$p<0.05$.

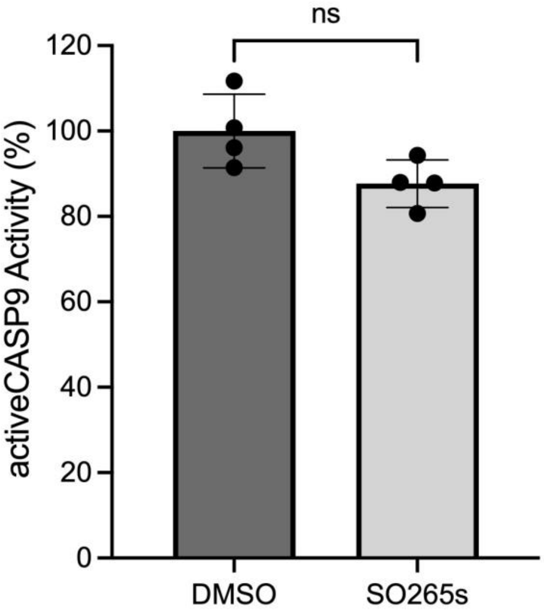


**Figure S34. SO265s modestly inhibit activeCASP9**. Active enzyme (1 µM) in PBS was treated with 100 µM of **SO265s** for 1h at ambient conditions. Data represents mean values and standard deviations (n = 4 biological replicates). Statistical significance was calculated with unpaired Student's t-tests, ns = not significant.

A

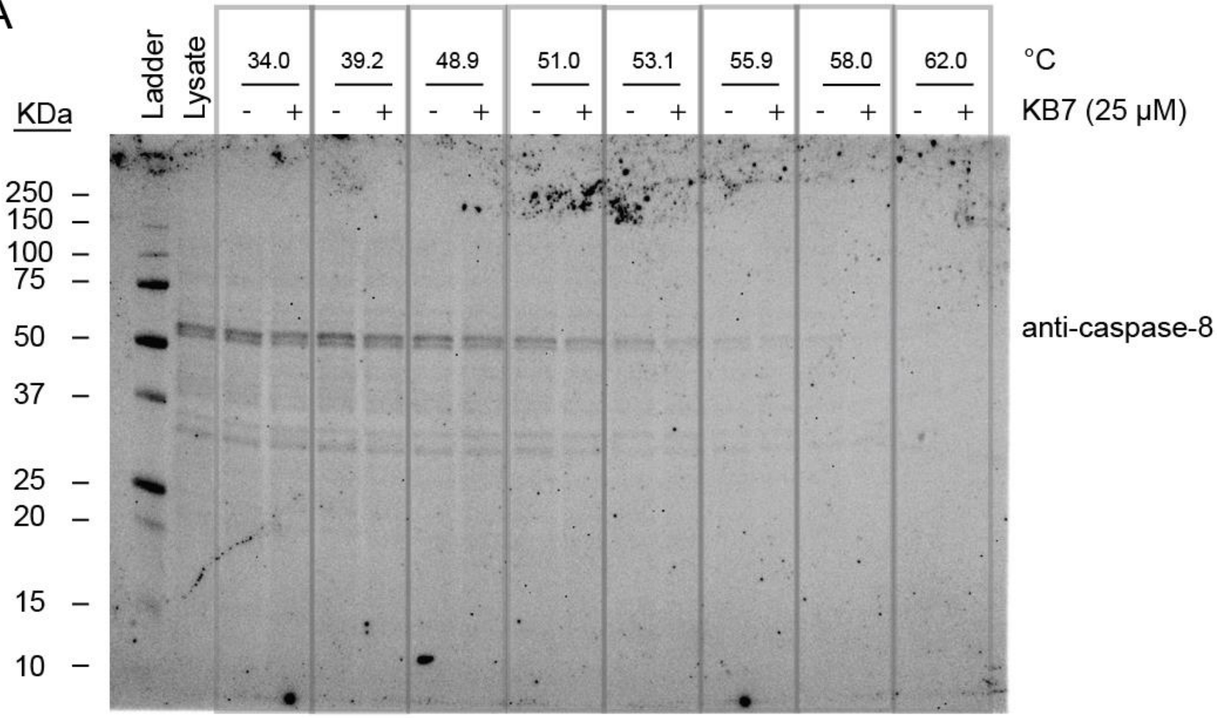
Ladder
Lysate
34.0
39.2
48.9
51.0
53.1
55.9
58.0
62.0
°C
KB7 (25 µM)
- +
KDa
250
150
100
75
50
37
25
20
15
10
anti-caspase-8

B

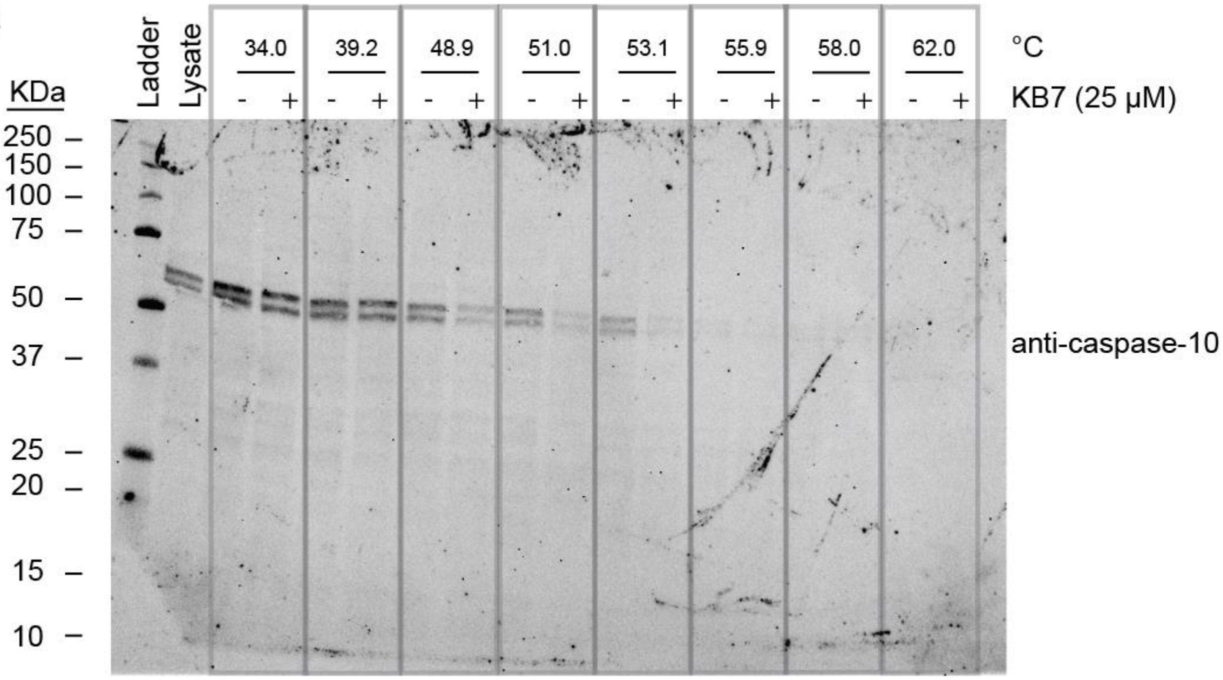
Ladder
Lysate
34.0
39.2
48.9
51.0
53.1
55.9
58.0
62.0
°C
KB7 (25 µM)
- +
KDa
250
150
100
75
50
37
25
20
15
10
anti-caspase-10

C

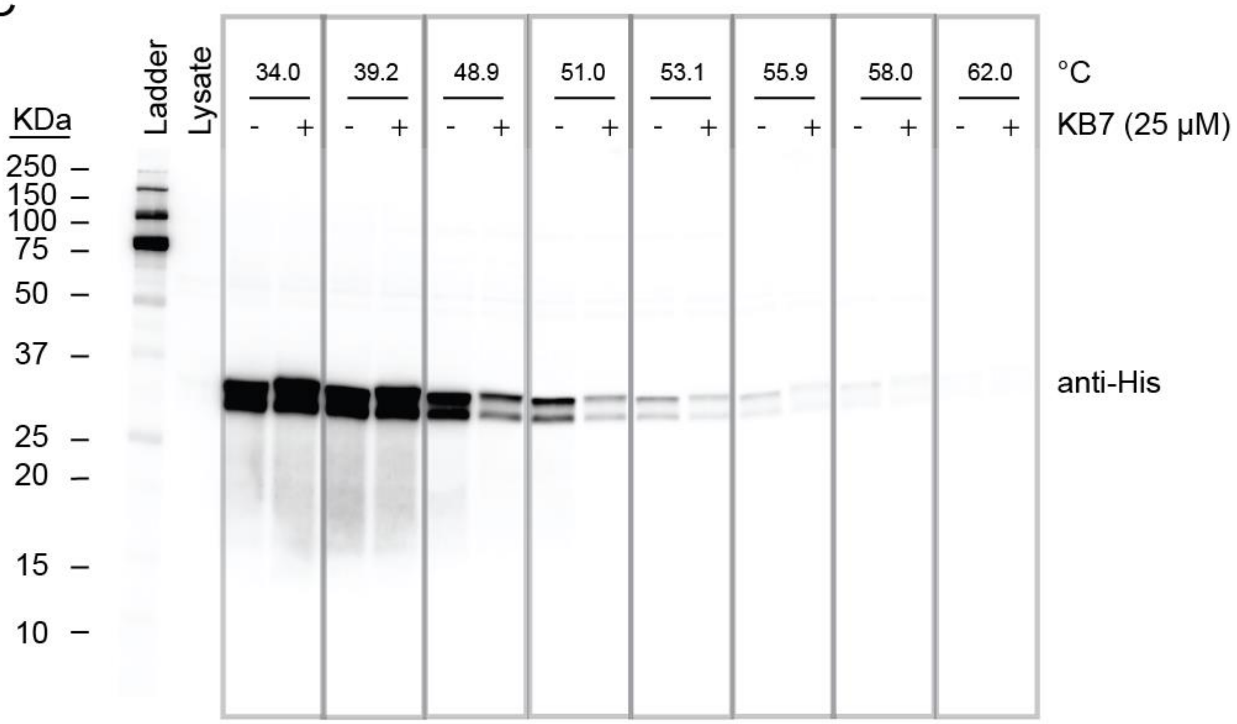

KDa
Ladder
Lysate
34.0
39.2
48.9
51.0
53.1
55.9
58.0
62.0
°C
- + - + - + - + - + - + - + - +
KB7 (25 µM)
250
150
100
75
50
37
25
20
15
10
anti-His


D

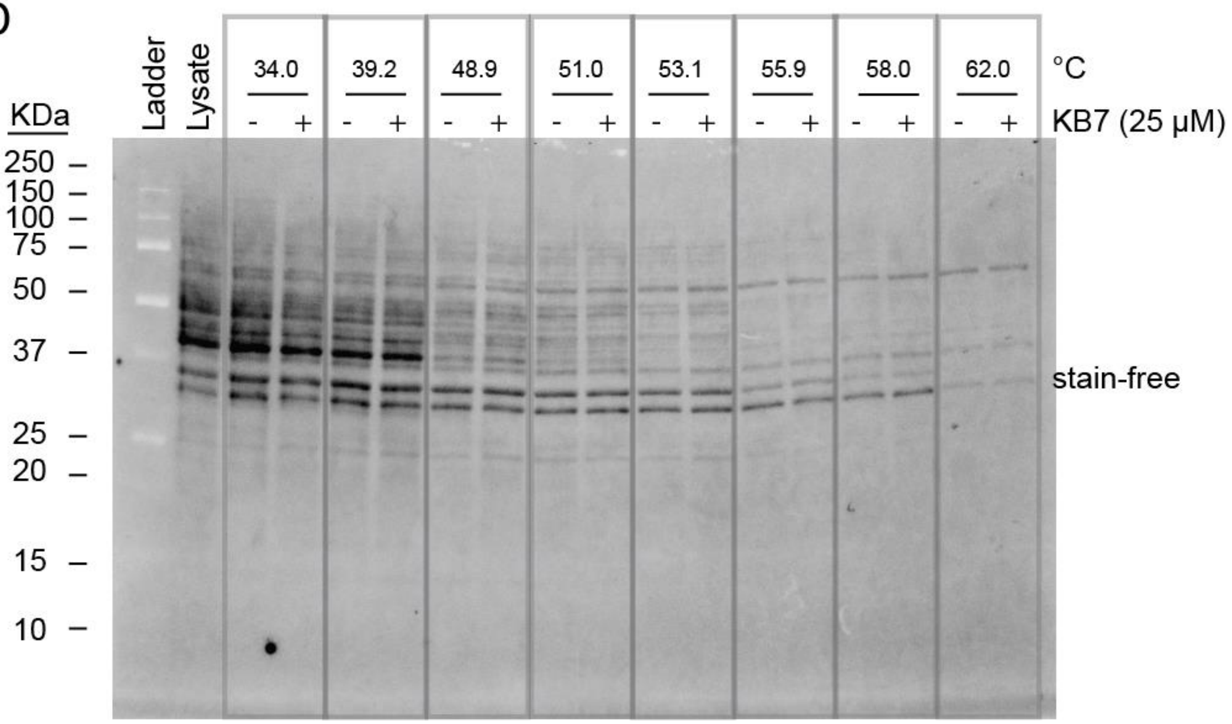

KDa
Ladder
Lysate
34.0
39.2
48.9
51.0
53.1
55.9
58.0
62.0
°C
- + - + - + - + - + - + - + - +
KB7 (25 µM)
250
150
100
75
50
37
25
20
15
10
stain-free

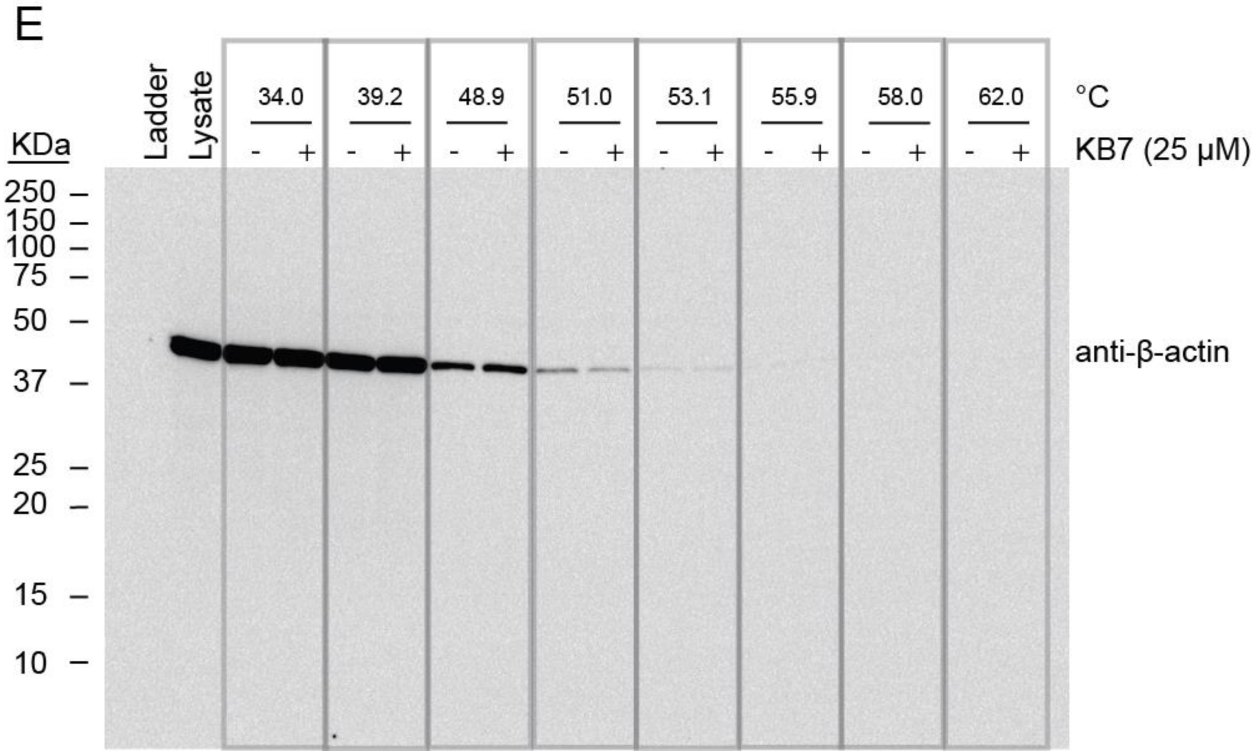


**Figure S35. Establishing CETSA assay using positive control KB7**. Recombinant caspase-10 (proCASP10) was spiked into 3.1 mg/mL Jurkat lysates and treated with KB7 (25 µM) for 1h at ambient conditions. Samples were treated at indicated temperatures for 5 min in a Thermal Cycler. Samples were blotted onto a nitrocellulose membrane followed by imaging for pro-caspase-8 using an (A) anti-caspase-8 primary antibody, (B) blotted for pro-caspase-10 using an anti-caspase-10 primary antibody, (C) blotted for His tag using an anti-His primary antibody. Loading was imaged using (D) stain-free and (E) by blotting for ß-actin using an anti-ß-actin primary antibody.

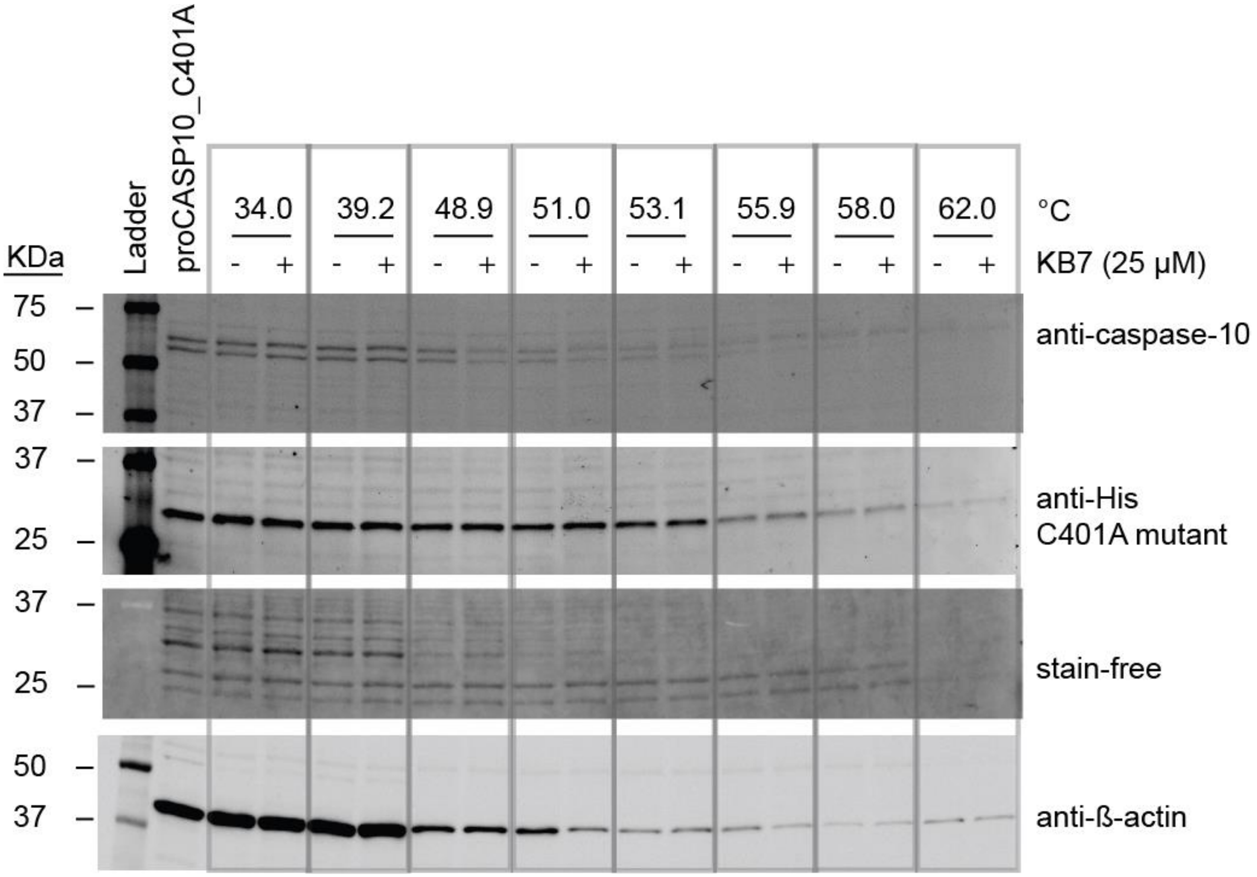


**Figure S36. KB7 treatment has little effect on thermal stability of recombinant proCASP10_C401A**. Recombinant caspase-10 catalytic mutant (proCASP10_C401A) was spiked into 2.4 mg/mL Jurkat lysates and treated with KB7 (25 µM) for 1h at ambient conditions. Samples were treated at indicated temperatures for 5 min in a Thermal Cycler. Samples were blotted onto a nitrocellulose membrane followed by imaging for the indicating antibodies.

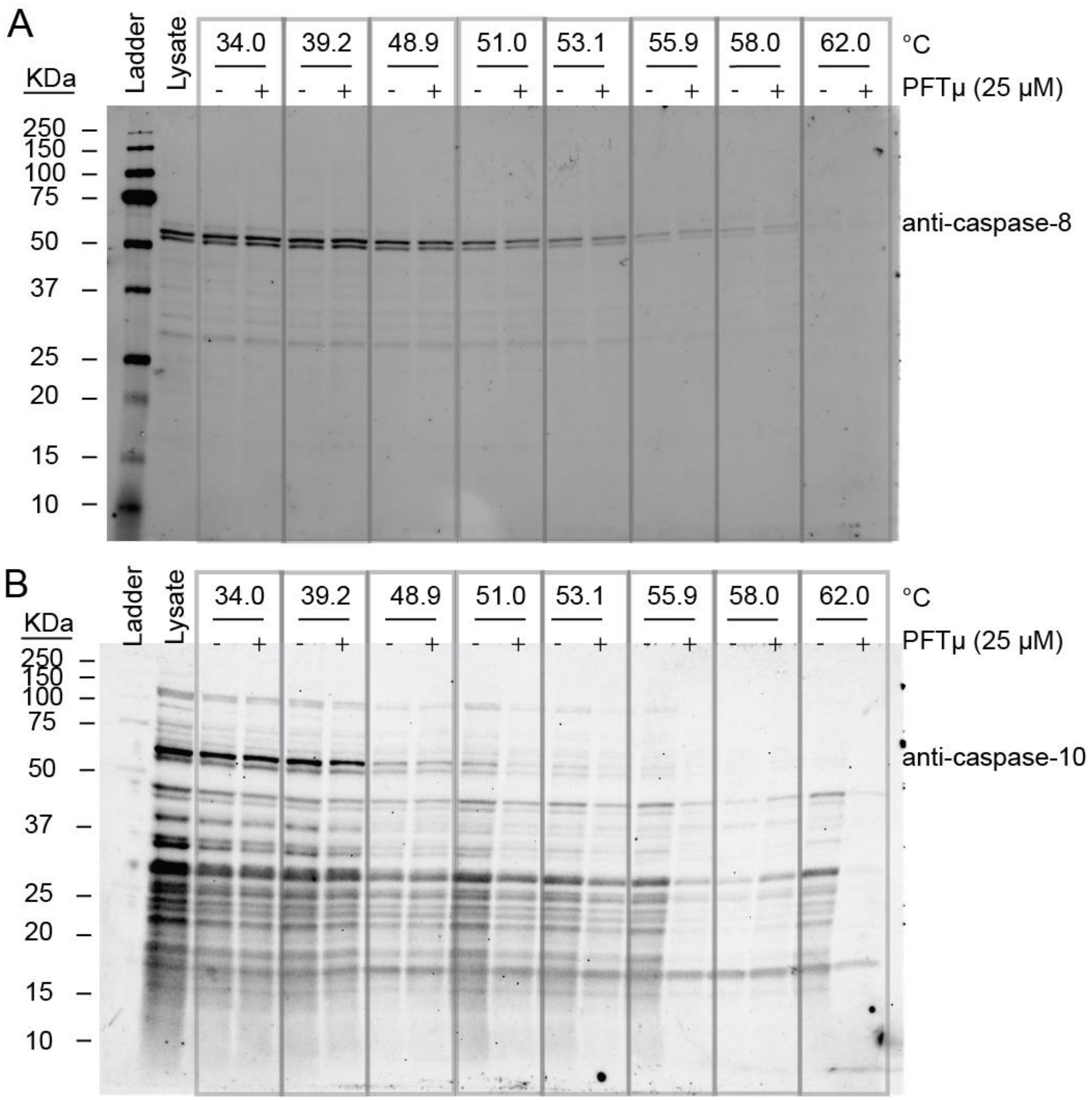
A
KDa
Ladder
Lysate
34.0
39.2
48.9
51.0
53.1
55.9
58.0
62.0
°C
PFTμ (25 μM)
250
150
100
75
50
37
25
20
15
10
anti-caspase-8
B
anti-caspase-10

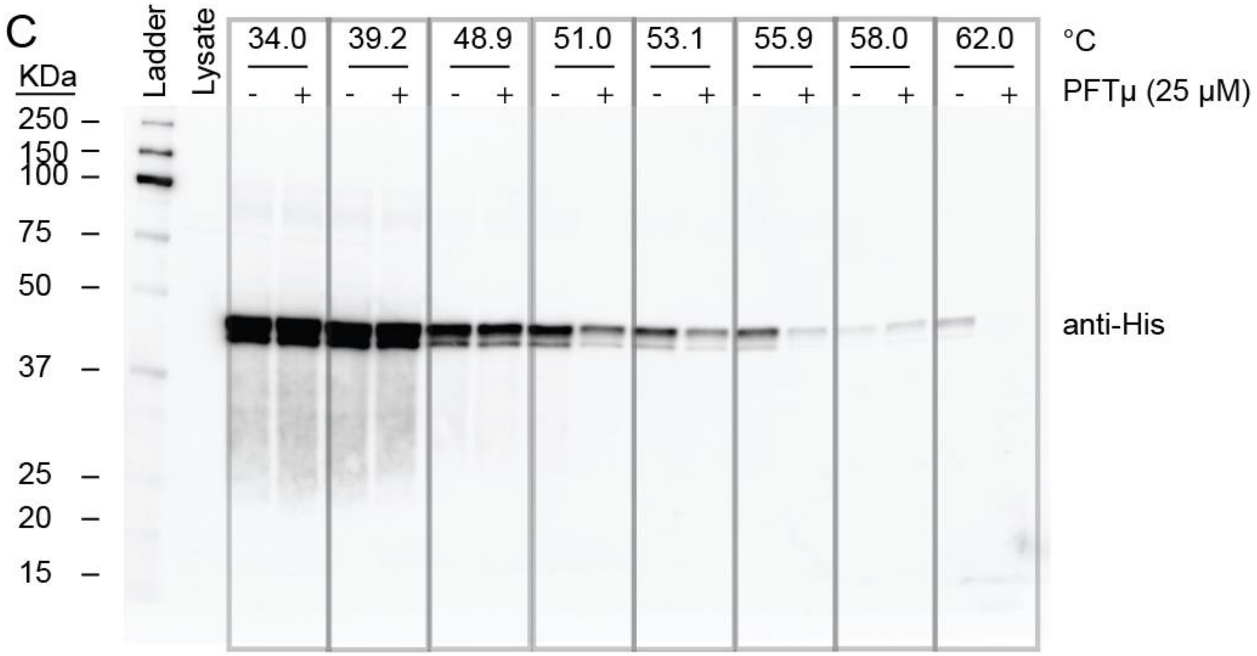
C
KDa
250
150
100
75
50
37
25
20
15
Ladder
Lysate
34.0
39.2
48.9
51.0
53.1
55.9
58.0
62.0
°C
- + - + - + - + - + - + - + - +
PFTμ (25 μM)
anti-His

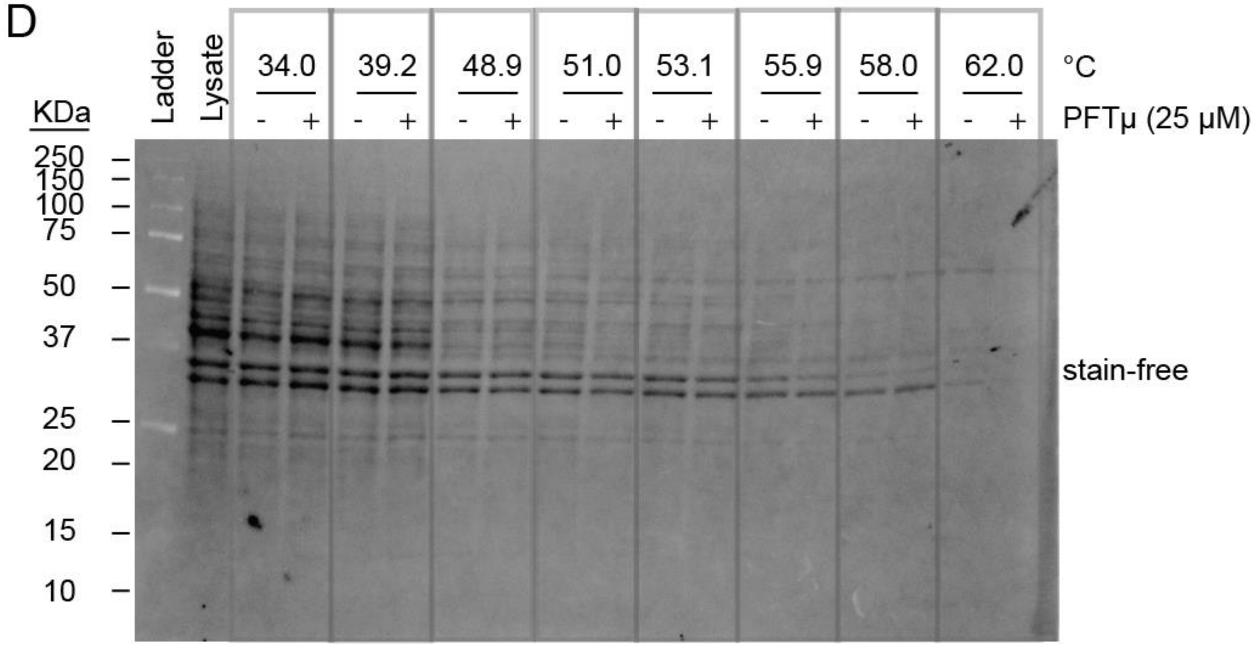
D
KDa
250
150
100
75
50
37
25
20
15
10
Ladder
Lysate
34.0
39.2
48.9
51.0
53.1
55.9
58.0
62.0
°C
- + - + - + - + - + - + - + - +
PFTμ (25 μM)
stain-free

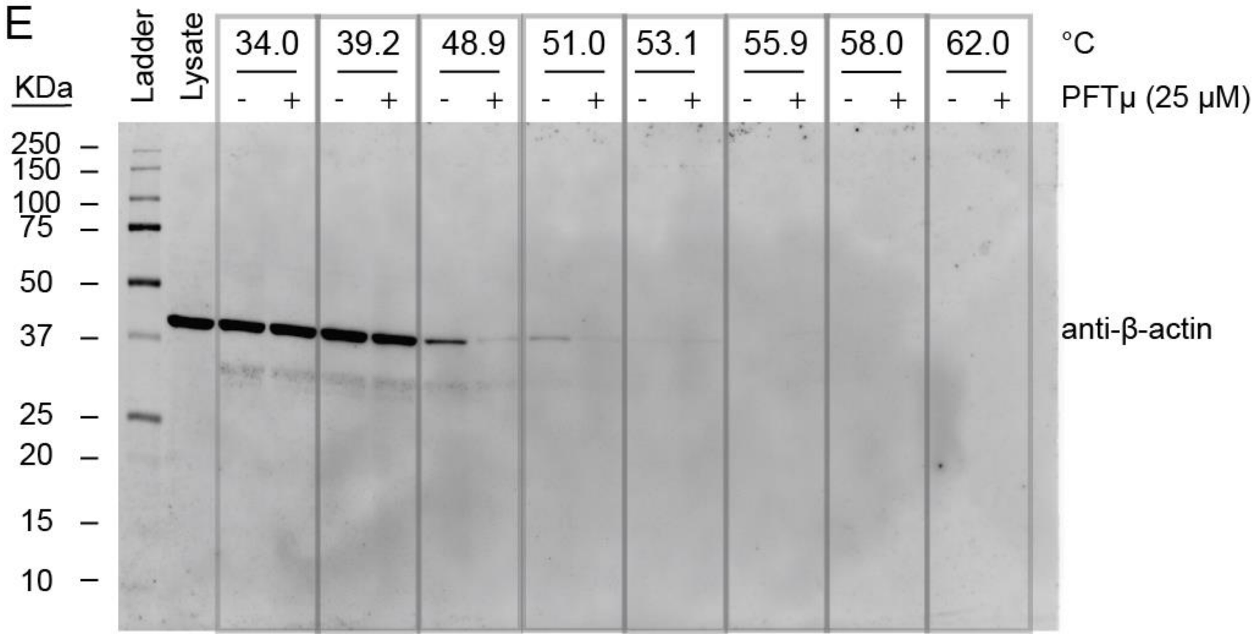


**Figure S37. CETSA analysis reveals that PFTμ destabilizes caspase-10.** Recombinant caspase-10 (proCASP10) was spiked into 2.8 mg/mL Jurkat lysates and treated with PFTμ (25 μM) for 1h under ambient conditions. Samples were treated at the indicated temperatures for 5 min in a Thermal Cycler. Samples were blotted onto a nitrocellulose membrane followed by imaging for procaspase-8 using an (A) anti-caspase-8 primary antibody, (B) blotted for pro-caspase-10 using the anti-caspase-10 primary antibody, (C) blotted for His tag using the anti-His primary antibody. (D) Loading was imaged using stain-free and (E) by blotting for ß-actin using the anti-ß-actin primary antibody.

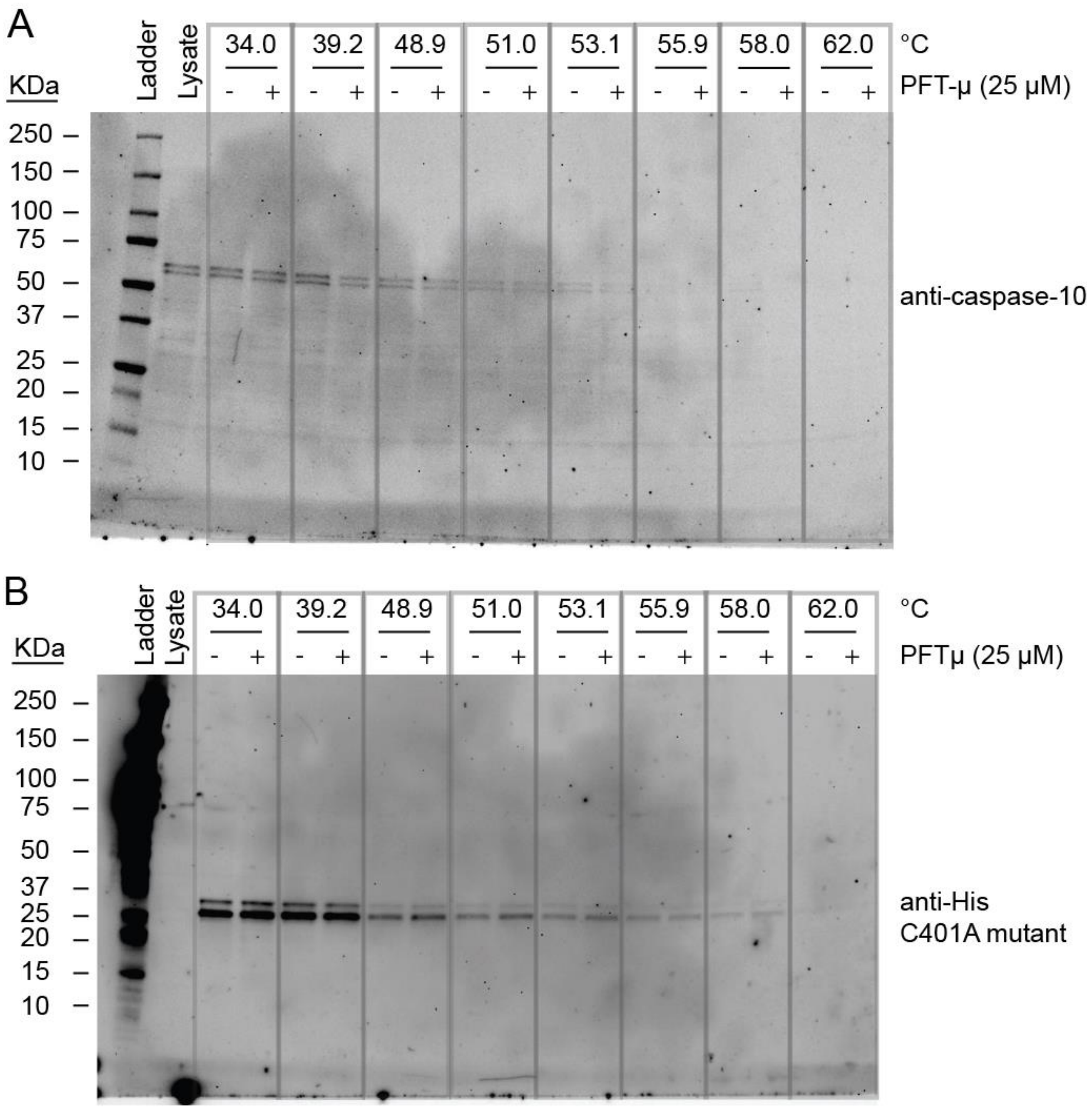
A
KDa
Ladder
Lysate
34.0
39.2
48.9
51.0
53.1
55.9
58.0
62.0
°C
PFT-μ (25 μM)
250
150
100
75
50
37
25
20
15
10
anti-caspase-10
B
KDa
Ladder
Lysate
34.0
39.2
48.9
51.0
53.1
55.9
58.0
62.0
°C
PFTμ (25 μM)
250
150
100
75
50
37
25
20
15
10
anti-His
C401A mutant

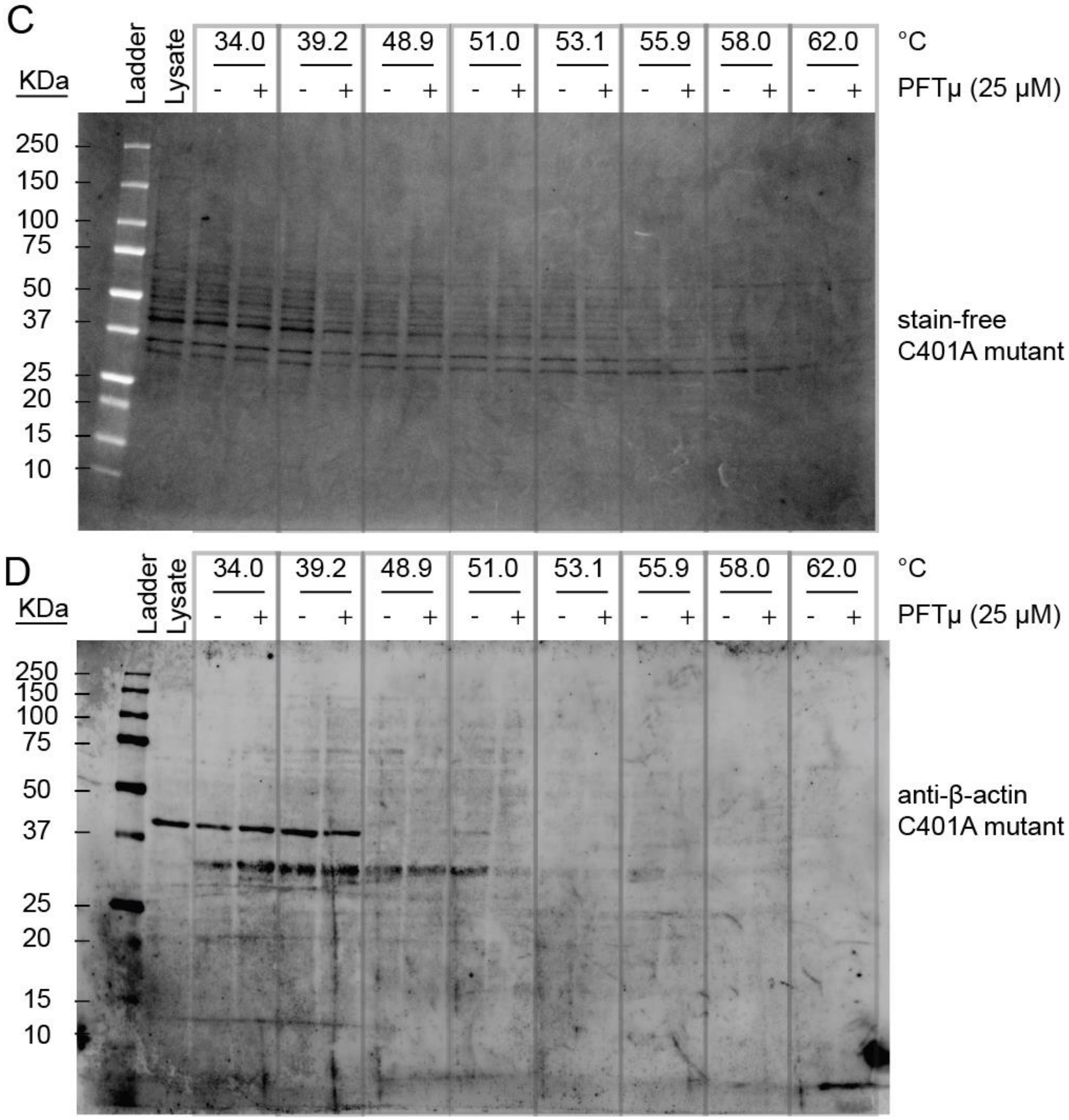


**Figure S38. CETSA analysis reveals that PFTμ does not destabilize recombinant pro-caspase-10_C401A.** Recombinant pro-caspase-10 C401A mutant (1 μM) spiked into 3.3 mg/mL Jurkat lysates treated with PFTμ (25 μM) for 1h at ambient conditions. Samples were treated at the indicated temperatures for 5 min in a Thermal Cycler. Samples were blotted onto a nitrocellulose membrane followed by (A) imaging for pro-caspase-10 using the procaspase-10 using the anti-caspase-10 primary antibody, (B) blotted for His tag using the anti-His primary antibody. (C) Loading was imaged using stain-free and (D) by blotting for ß-actin using the anti-ß-actin primary antibody.

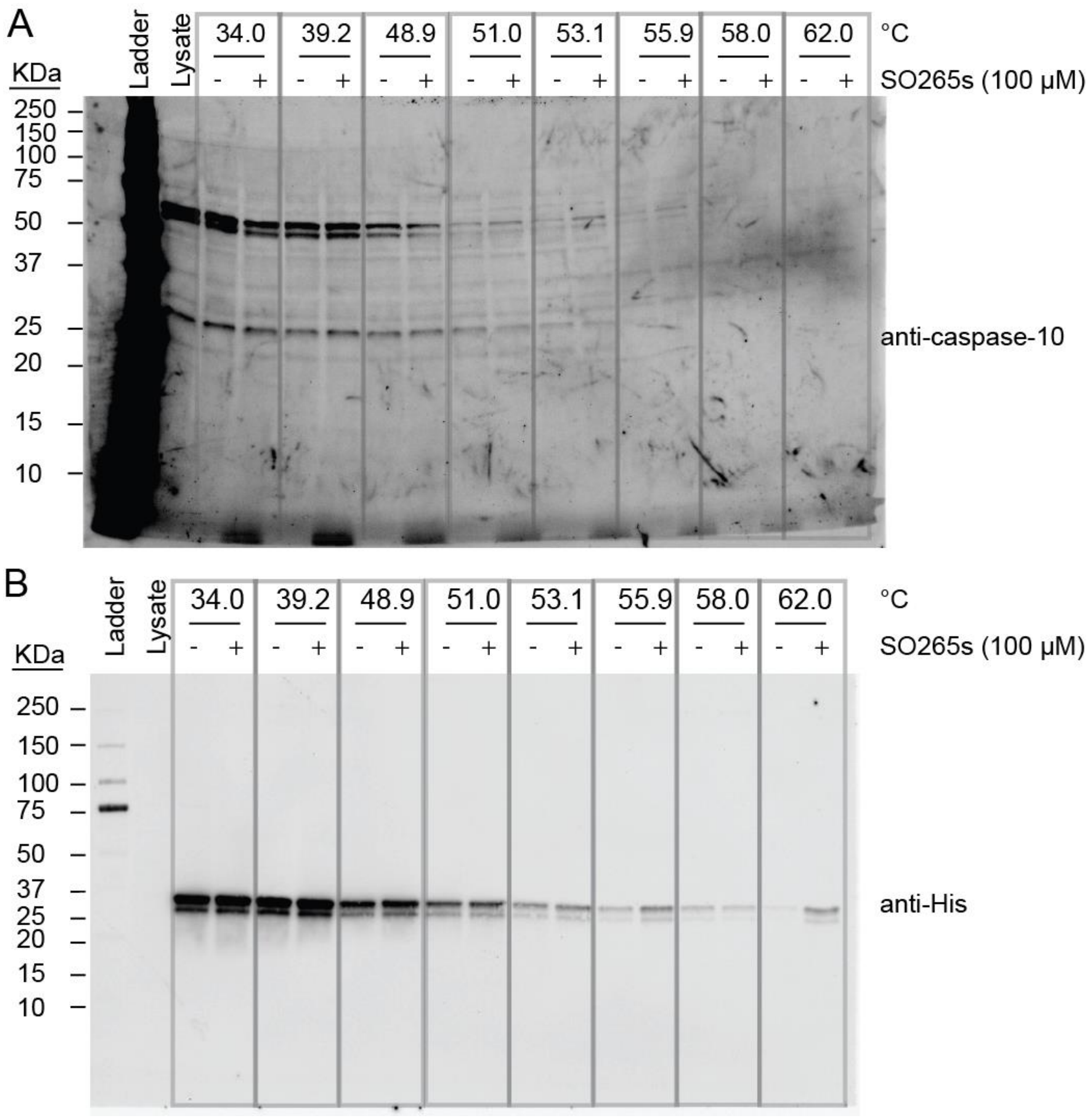
A
KDa
Ladder
Lysate
34.0
39.2
48.9
51.0
53.1
55.9
58.0
62.0
°C
SO265s (100 μM)
250
150
100
75
50
37
25
20
15
10
anti-caspase-10
B
anti-His

C

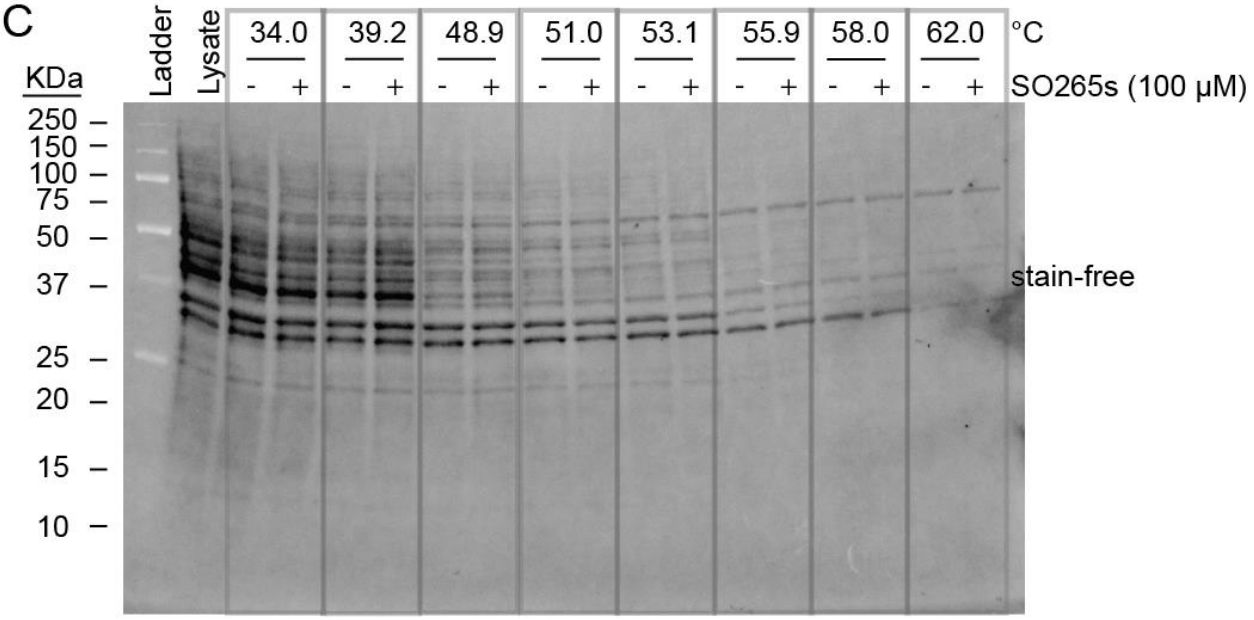


D

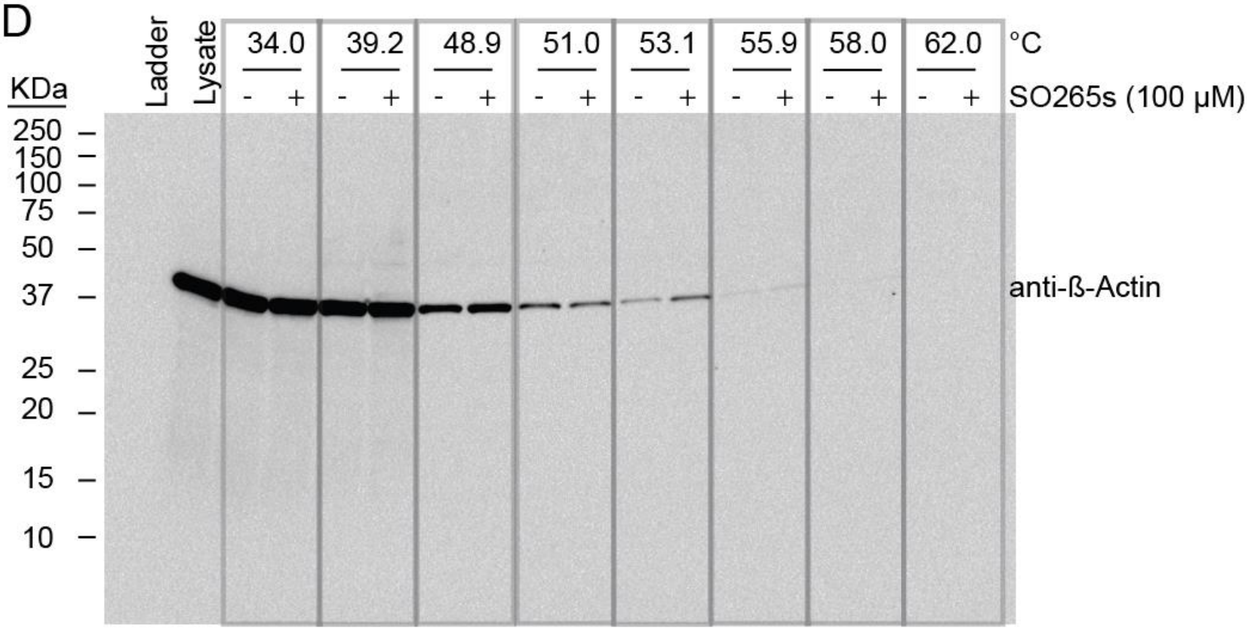


**Figure S39**. **CETSA analysis reveals slight stabilization of endogenous caspase-10 and recombinant caspase-10 by SO265s.** Recombinant procaspase-10 (proCASP10) (1 μM) was spiked into 3.0 mg/mL Jurkat lysates followed by treatment with **SO265s** (100 μM) for 1h at ambient conditions. Samples were treated at indicated temperatures for 5 min in a Thermal Cycler. Samples were blotted onto a nitrocellulose membrane followed (A) by imaging for pro-caspase-10 using the anti-caspase-10 primary antibody, (B)blotted for His tag using the anti-His primary antibody. (C) Loading was imaged using stain-free and (D) by blotting for ß-actin using the anti-ß-actin primary antibody.

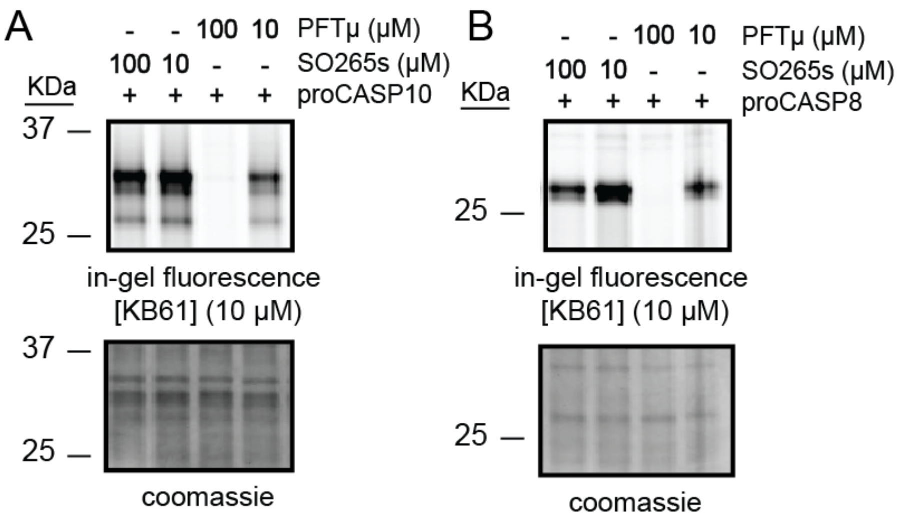


**Figure S40. PFTμ competes for labeling of caspase-10 and caspase-8 against KB7 but SO265s only modestly competes for labeling of caspase-8.** Competitive ABPP gel of Jurkat lysates (1 mg/ mL) spiked with (A) proCASP10 at 1 μM or (B) proCASP8 at 1 μM followed by treatment with **PFTμ** or **SO265** at the indicated final concentrations (100 μM and 10 μM) for 1h followed by treatment with **KB61** (10 μM) for 1h. Samples were then clicked to rhodamine azide (25 M) for 1h and visualized using in-gel fluorescence.

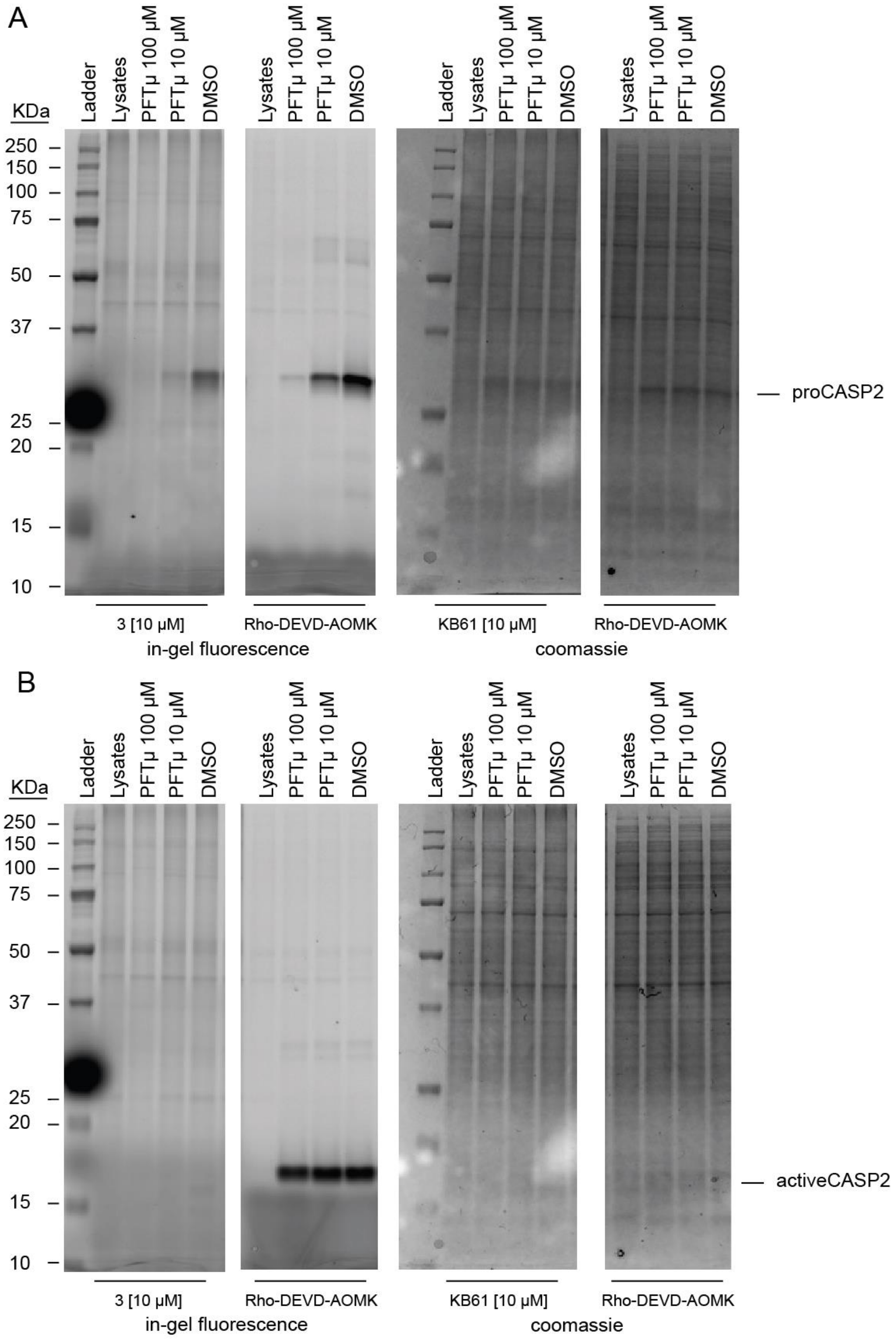


**Figure S41**. **PFTμ competes for labeling of proCASP2 but not activeCASP2**. Recombinant procaspase-2 (proCASP2) and active caspase-2 (activeCASP2) were spiked into 1 mg/mL HEK293T lysates at a final concentration of 1 μM. Samples were treated first with **PFTμ** at 100 μM and 10 μM or vehicle control DMSO for 1h followed by treatment with the click probe **3** [3] for an additional hour at ambient conditions. Samples were then clicked onto rhodamine-azide and visualized by gel-fluorescence. Gels were stained with Coomassie.

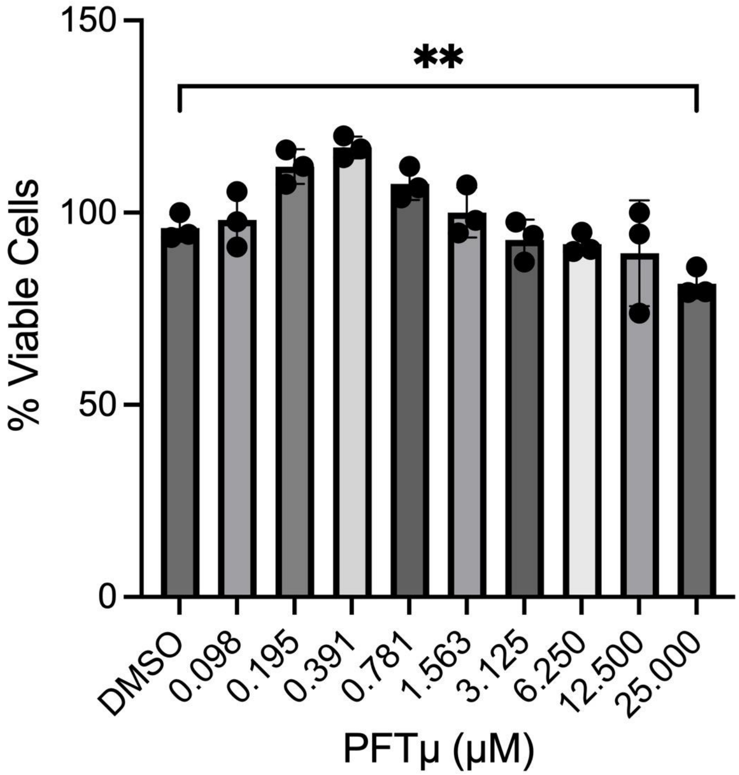


**Figure S42**. **PFTμ treatment results in minimal cytotoxicity in Jurkat cells**. Cell viability assay with PFTμ serial dilutions in Jurkat cells treated with compound for 1h followed by CellTiter Glo. Data represents mean values and standard deviations (n = 3 biological replicates). Statistical significance was calculated with unpaired Student's t-tests, **$p<0.001$.

### (C) Supplementary Tables

**Table S1** and **Table S2.** Datasets corresponding to each figure, provided in the attached supplementary files (.xlsx).

**Table S3.** Summary list of recombinant caspase-8 and caspase-10 constructs used in this study.

| Construct | Mutation | Source |
|---|---|---|
| activeCASP8 | n/a | [1] |
| proCASP8 | D347A, D384A, C409S, C433S | [1] |
| activeCASP3 | n/a | Abcam (ab52101) |
| activeCASP2 | n/a | [3] |
| activeCASP9 | n/a | Sigma-Aldrich (218807) |
| proCASP10 | D415A | [1] |
| activeCASP10 | n/a | [1] |
| proCASP10_C401A | C401A | This study |
| proCASP10TEV | D415 → ENLYFQG | This study |
| proCASP2xTEV | D415 and D428 → ENLYFQG | This study |
| proCASP10TEV Linker | D415 → AAENLYFQG | This study |

**Table S4.** List of caspase-10 primer sequences used in this study.

| **Purpose** | **Primer Sequence** | |
|---|---|---|
| TEV insertion 1 | GAAAATCTCTACTTCCAGGGCGGTAAAGAAAGTTGCCG | Forward |

| | | |
|---|---|---|
| D415 to AAENLYFQG | CAGGTTTTCTGCTGCTTCGATGGATACGGAAGGCTGTATCTC | Reverse |
| TEV insertion 2 D428 to AAENLYFQG | GGCACCCACTTCCCTGCAGGCAGCAGAAAACCTGTATTTTCAG GGCAG | Forward |
| | CGGCAGGAATACTGCCCTGAAAATACAGGTTTTCTGCTGCCTGC AGGG | Reverse |
| TEV insertion No Linker D415 to ENLYFQG | GCCTTCCGTATCCATCGAAAACCTGTATTTTCAGGGCGCTCTGA A | Forward |
| | CAGAGCGCCCTGAAAATACAGGTTTTCTGCTTCGATGGATACGG AAGG | Reverse |
| pET23B subcloning (NdeI) | CAGATCATATGGTTAAGACATTCTTGGAAGCCTTACC | forward |

| | | |
|---|---|---|
| pET23B subcloning (XhoI) | CGGCCTCGAGTAATGAAAGTGCATCCAGGGGC | reverse |
| D428A | CCCACTTCCCTGCAGGCCAGTATTCCTGCCGAGGCTGACTTCC | Forward |
| | GCAGGAATACTGGCCTGCAGGGAAGTGGGTGCCTGCTCAGGG | Reverse |
| D415A | CCTTCCGTATCCATCGAAGCAGCTGCTCTGAACCCTGAGCAGG | Forward |
| | CCTGCTCAGGGTTCAGAGCAGCTGCTTCGATGGATACGGAAGG | Reverse |
| C401A | CCTAAACTCTTTTTCATCCAGGCCGCCCAAGGTGAAGAGATACAGC | forward |
| | GCTGTATCTCTTCACCTTGGGCGGCCTGGATGAAAAAGAGTTTAGGT | Reverse |

**Table S5.** Files in Proteomics Identification Database (PRIDE) datasets. PRIDE IDENTIFIER: PXD053315.

| Figure | File name | Compound | Experiment |
|---|---|---|---|
| Figure 5B | 2023_05_09_KB_noFAIMS_KB_JOC_chymo_tryp_double_5 | PFTμ | HL biotin azide |
| | 2023_05_09_KB_noFAIMS_KB_JOC_chymo_tryp_double_6 | PFTμ | HL biotin azide |
| | 2024_06_03_KB_NoFAIMS_JOC_biotinazide_PFT_T1_rerun_0523 | PFTμ | HL biotin azide |
| | 2024_06_03_KB_NoFAIMS_JOC_biotinazide_PFT_T2_rerun_0523 | PFTμ | HL biotin azide |
| | 2024_06_11_KB_FAIMS_JOC1_biotinazide_T1_rerun1_0523 | PFTμ | HL biotin azide |
| | 2024_06_11_KB_FAIMS_JOC2_biotinazide_T2_rerun2_0523 | PFTμ | HL biotin azide |

### (D) Biology Methods

*Cell lines, culture conditions*

Jurkat cells were cultured in Roswell Park Memorial Institute (RPMI) media (Fisher Scientific, 11875119) with 10% fetal bovine serum (Avantor Seradigm Lot # 214B17) and 100U/mL penicillin and 100U/mL streptomycin at 37oC and 5% $CO_2$. Jurkat cells were obtained from ATCC (TIB-152). ΔΔG3BP1/2 KO U2OS cells stably expressing GFP-G3BP1 were a generous gift from the lab of Dr. Melody Li and were originally generated in the lab of Dr. Paul Anderson[4]. U2OS cells were cultured in Dulbecco's Modified Eagle Medium (DMEM) media (Fisher Scientific, 11995073) supplemented with 10 % fetal bovine serum (Avantor Seradigm Lot # 214B17) and 1 % antibiotics (Pen/Strep, 100 U/mL). Cell culture reagents including RPMI 1640 and DMEM media, trypsin-EDTA and penicillin/streptomycin (Pen/Strep) were purchased from Fisher Scientific.

*Mycoplasma testing*

Mycoplasma testing was conducted monthly using the MycoAlert® kit (LT07-703, Lonza Rockland, Rockland, ME) following the manufacturer's instructions.

*Cell Harvesting*

Suspension cells were centrifuged at 1,400 x *g* for 5 minutes and the supernatant was aspirated. The pellets were then washed in 10 mL PBS and centrifuged at 1,400 *g* for 5 minutes. PBS wash was repeated, and the subsequent pellet was then resuspended in 1 mL PBS in a microcentrifuge tube and centrifuged at 1,400 *g* for 5 minutes. The supernatant was aspirated, and the cell pellets were stored at -80 oC.

*Cell Lysis*

Jurkat lysates used for competitive ABPP gel analysis were lysed using an Ultrasonic Probe Sonicator at Power 2 for 10 pulses, 1-second pulse, and 1 second off on the ice. For western blotting samples, cells were reconstituted and lysed using 100 uL of cold 0.3% 3-[(3-Cholamidopropyl) dimethylammonio]-1-propanesulfonate (CHAPS) buffer in PBS. The reconstituted cell pellet was left to incubate with the CHAPS buffer on ice for 15 min. The samples were then harvested by centrifugation (1,400 x *g*, 10 min), and the clarified supernatant was then transferred to a new tube.

*Subcloning of caspase-10*

Caspase-10 (Q92851-4, 10-L isoform) expression vector was obtained as a kind gift from Dr. Michael Lenardo. The sequence encoding caspase-10, lacking the prodomain (residue numbers 220-521) was then subcloned into the pET23b(+) at ndeI and xhoI fused to a C-terminal hexahistidine-tag. Plasmids were propagated in chemically competent TOP10 cells.

*Mutagenesis*

Point mutations (C401A, D415A, D428, and insertions of TEV cleavage motifs at D415) were created by PCR-based site-directed mutagenesis using the primers in **Table S2**. Plasmids were propagated in TOP10 chemically competent cells and isolated by Zyppy Plasmid Miniprep following the manufacturer's protocol (Zymo Research, D4037).

*Protein expression and purification*

The sequence encoding caspase-10, lacking the prodomain (truncated at D219 and earlier residues) was subcloned into the pET23b(+) with a C-terminal hexaHis-tag. Point mutations (C401A, D415A, D428A) were created by PCR-based site-directed mutagenesis. TEV insertion mutations (ENLYFQG and AAENLYFQG) were created by PCR-based site-directed mutagenesis. Plasmids were propagated in TOP10 chemically competent cells. Single colonies from TOP10 grown cells were collected in 5 mL of LB supplemented with 100 µg/ml ampicillin and grown overnight (16h). Cells were harvested the following day and subjected to Zyppy Plasmid Miniprep following the manufacturer's protocol (Zymo Research, D4037). Following sequencing of plasmids, validated plasmids were transformed to BL21(DE3) *E. coli* cells. Single colonies were picked from an LB agar plate and grown in 10 mL of LB media supplemented with 100 µg/mL ampicillin at 37 oC for 18h. The cell culture was then transferred and grown in 1 L of Miller LB medium at 37°C to an optical density (OD600) of 0.5. The TEV cleavable construct cultures were then cooled to 18°C, induced with 1 mM isopropyl-β-D-galactopyranoside (IPTG), and incubated for an additional 4 h at 18°C. he other active- and pro-constructs were cooled to 18°C, induced with 1 mM IPTG, and incubated overnight (16-18 h) at 18°C. The cells were centrifuged at 8,000 x *rpm* for 45 min. All cell pellets were stored at -80°C except for the TEV cleavable constructs (i.e.

proCASP10TEV Linker) which was lysed and processed immediately after induction. The cells were resuspended in 10 mL lysis buffer (100 mM Tris pH 7.5, 100 mM NaCl, 25 mM Imidazole) per 1 g of cells. The resuspended cells were passed through a microfluidizer (Avestin Emulsiflex C3 Homogenizer; 8,000 psi x2 rounds) to ensure lysis. The cell debris was removed by centrifugation (20,000 x *g*, 45 min, at 4 oC) and the supernatant was resuspended with 1 mL of Hispur Ni-NTA agarose resin (Thermo Scientific$^{TM}$, PI88222). The sample was washed with lysis buffer (2 x 50 mL). His-tagged caspase-2 was eluted from the resin using an elution buffer with high imidazole concentration (100 mM Tris pH 7.5, 100 mM NaCl, 250 mM imidazole). The eluted sample was concentrated (Amicon Ultra Centrifugal Filter Unit, 4 mL 10 kDa, Fisher Scientific, UFC801024) and buffer exchanged via PD10 desalting column (Cytiva, GE17-0851-01) into storage buffer (20 mM Tris pH 7.5, 50 mM NaCl, 5 mM DTT). Recombinant TEV protease was purchased from the Berkeley MacroLab QB3, where it was was expressed as a double mutant (L56V / S135G) pRK793 plasmid in Rosetta2(DE3) pLysS cells (TEV-DM-Prk793 L56V/ S135G) and stored in 25 mM HEPES pH 7.5, 400 mM NaCl, 10% glycerol, 1 mM DTT. Recombinant proCASP8 and proCASP10 were purified , as reported previously[1].

*Recombinant TEV protease*

Recombinant TEV protease was purchased from the Berkeley MacroLab QB3, where it was was expressed as a double mutant (L56V / S135G) pRK793 plasmid in Rosetta2(DE3)pLysS cells (TEV-DM-Prk793 L56V/ S135G) and stored in 25 mM HEPES pH 7.5, 400 mM NaCl, 10% glycerol, 1 mM DTT.

*Recombinant Caspases*

Recombinant active, proCASP8, proCASP10, proCASP10TEV, and proCASP2TEV were purified as reported previously[1]. In brief, plasmids were transformed to BL21(DE3) *E. coli* cells (New England Biolabs). Starter cultures were grown from single colonies in 10 mL of LB media under ampicillin selection (100 µg/mL) at 37°C for 18 hours.  1 L of Miller LB medium was inoculated with each starter culture (10 mL) and grown with shaking (200 rpm) at 37°C to an optical density (OD600) of 0.5. The culture was then cooled (18°C) and induced for 4h at 18°C with 1 mM IPTG. The cells were centrifuged at 8,000 x *rpm* for 45 min, and the cell pellet mass was measured. The cells were resuspended in 10 mL lysis buffer (100 mM Tris pH 7.5, 100 mM NaCl, 25 mM imidazole) per 1 g of cells. The resuspended cells were passed through a microfluidizer (Avestin Emulsiflex C3 Homogenizer; 8,000 psi x2 rounds) to ensure lysis. The cell debris was removed by centrifugation (20,000 x *g*, 45 min*)*, and the supernatant was resuspended with 1 mL of Hispur Ni-NTA agarose resin (Thermo Scientific$^{TM}$, PI88222). The sample was washed with lysis buffer (2 x 50 mL). His-tagged recombinant caspase was eluted from the resin using an elution buffer with a high imidazole concentration (100 mM Tris pH 7.5, 100 mM NaCl, 250 mM imidazole). The eluted sample was concentrated (Amicon Ultra Centrifugal Filter Unit, 4 mL 10 kDa, Fisher Scientific, UFC801024), and buffer was exchanged via PD10 desalting column (Cytiva, GE17-0851-01) into storage buffer (20 mM Tris pH 7.5, 50 mM NaCl, 5 mM DTT).

*High throughput screening*

proCASP10TEV Linker activity was analyzed per well in a low-volume 384-well plate (Fisher Scientific, 784900). First, 10 µL of proCASP10TEV Linker (in PBS) was added to all wells using the Aquamax liquid handler. Each plate was then pinned with compounds (10 µM) or DMSO using a 250 nL pin (Beckman Coulter Biomek FX system) followed by a 1 h incubation (95% humidity, 25 oC). Substrate solution (5 mM DTT, 10 µM Ac-VDVAD-AFC substrate, 333 mM citrate in PBS with either 667 nM TEV protease or no TEV protease) was then dispensed into each well using an Aquamax liquid handler (Aquamax DW4, Surplus Solutions, 324757). A total of 16 wells were treated with DMSO, and sodium citrate solution containing TEV protease was used as the negative control. A total of 16 wells were treated with DMSO and substrate solution without adding TEV protease and used as the positive controls. The first and last columns of the plates were not used. Z'-factor was calculated per well using the mean and standard deviation of the negative and positive controls (endpoint data) as the first filtering step. From a total of 375 plates, the average Z'-factor calculated was 0.58. Each 384-well plate had a total of 320 wells that were pinned and treated with compounds. The total number of compounds was approximately 120,000 (320 wells x 375 plates). After filtering out plates outside the Z'-factor range (0.5 - 1.0), the resulting number of plates was 327. The average Z'-factor for these plates was 0.72 (from a total of 104,640 compounds) (**Table S1**). The next filtering step was determined with the help of z-scores (refer to *Z'-factor and z-score calculations* methods section). In our study, values with z-scores lower than -3 or those with 3 significant figures lower than the average of the DMSO controls per plate. A total of 237 hit compounds were obtained with a z-score of -3 or lower and used for re-screen, counter-screen, and validation assays.

*Re-screening and counter screening (proCASP10TEV and activeCASP10)*

proCASP10TEV Linker and activeCASP10 were analyzed per well in a low-volume 384-well plate each. First, 10 µL of proCASP10TEV Linker or activeCASP10 (in PBS) were added to all wells using the Mantis Precise Liquid Dispenser (Formulatrix). Each plate was then pinned with compounds (10 µM) or DMSO using a 250 nL pin (Beckman Coulter Biomek FX system) followed by a 1h incubation in ambient conditions. Substrate solution was then dispensed using the Mantis Liquid Dispenser. The substrate solution consisted of 5 mM DTT, 10 µM Ac-VDVAD-AFC substrate, 333 mM citrate in PBS for the counterscreen plate (activeCASP10), and the same substrate solution with the addition of TEV protease at 667 nM was used for the re-screen plate (proCASP10TEV Linker). A total of 16 wells were treated with DMSO and used as the positive control wells. For the re-screen plate, 16 wells were treated with a substrate solution in the absence of TEV protease (no TEV, negative controls). The first and last columns of the 384-well plate were not used. z-score was calculated per well using the mean and standard deviation of the positive controls (DMSO wells, n = 16) as shown in the *Z'-factor and z-score calculations* methods section.

*Gel-based ABPP*

Recombinant proteins were added to 1 mg/mL Jurkat cell lysates to a final concentration of 2 µM recombinant protein. 25 µL of the cell lysate-recombinant protein mixture was then treated with compounds (100 µM) or vehicle (DMSO) for 1h at ambient conditions. The samples were then incubated for 1h with 1 µL of 250 µM click probes (**IAA or KB61yne**) at a final concentration of 10 µM. Samples were then subjected to either click conjugation to rhodamine azide in 3 uL of click mix containing TBTA (1.5 µL of 1.7 mM for a final concentration of 100 µM), $CuSO_4$ (0.5 µL of 50 mM for a final concentration of 1 mM), Rhodamine-azide (0.5 µL of 1.25 mM for a final concentration of 25 µM), and TCEP (0.5 of µL 50 mM for a final concentration of 1 mM) or activity-based probe **Rho-DEVD-AOMK** (2 µM ). Next, 10 µL of 4x Laemmle loading dye (BioRad, 1610747) was added, and the samples were denatured at 95°C for 5 minutes. Samples were resolved by SDS-PAGE and imaged using a BioRad ChemiDoc Imaging System with visualization of protein loading with Coomassie InstantBlue (Fisher Scientific, ISB1L).

*TEV activation Gel-based ABPP*

Purified recombinant caspase-10-TEV cleavable construct at 2 µM final concentration in 1 mg/mL Jurkat lysates were first treated with either compound **KB61** or **Rho-DEVD-AOMK** at 10 µM final concentration for 1h at ambient conditions. proCASPTEV constructs were then activated with TEV protease (2 mg/mL stock) at increasing final concentrations (0 µM, 0.1 µM, 0.5 µM, 1.0 µM, 2.5 µM, 5.0 µM) for 1h at ambient conditions. Samples treated with compound **KB61** were then subjected to 'click' conjugation to rhodamine azide as prepared in the "Gel-based ABPP" section and visualized by SDS-PAGE in-gel fluorescence using a BioRad ChemiDoc Imaging System. Coomassie InstantBlue was used for the visualization of protein loading.

*Caspase activity assay*

Enzymatic activity of purchased active-caspase constructs (1 µM recombinant protein, 5 mM DTT, and 333 mM citrate in PBS) [caspase-3, Abcam, product number: ab52101; caspase-7, Abcam, product number: ab52173; caspase-9, Abcam, product number: ab52203] recombinant constructs (activeCASP10, proCASP2, proCASP8 and proCASP10), and TEV cleavable recombinant constructs (333 nM recombinant protein, 5 mM DTT, 333 mM citrate in PBS) were all determined using the fluorescent caspase-2 substrate Ac-VDVAD-AFC (Cayman Chemicals, item number 37351) at 10 µM final concentration. Activity assay is read on a BioTek Plate reader immediately after mixing active- or TEV-cleavable constructs with the substrate solution containing 5 mM DTT, 333 mM citrate pH 7.4, and 10 µM substrate. 7-amino-4-trifluoromethylcoumarin (AFC) released by substrate cleavage was detected at $\lambda_{ex}$ = 400 nm and $\lambda_{em}$ = 505 nm using a multimodal Synergy H1 microplate reader (BioTek). Reads were collected every minute after substrate addition. For the HCl acidic acid treatment, compound SO265 was prepared as a 10 mM stock in DMSO, 1 M citrate, 333 mM citrate, or 10 mM HCl and left to incubate at 37°C for 15 min. 10 mM stocks were directly used to treat proCASP10TEV Linker (333 nM in PBS) or TEV protease (667 nM in PBS) at a final concentration of 100 µM for 1h. Samples were then treated with substrate solution (10 µM substrate, 667 nM TEV protease, and 5 mM DTT in PBS). Percent activities were calculated from the slope of the linear range determined from the reaction progress curves. GraphPad Prism (Version Prism 10.2.3) was used to obtain $K_m$ and $k_{cat}$ values by fitting reaction velocities into the Michaelis-Menten equation using varied fluorogenic substrate Ac-VDVAD-AFC concentrations of 100 µM, 75 µM, 50 µM, 20 µM, 10 µM, 5 µM, 2.5 µM, and 1.0 µM.

*TEV protease activity assay*

TEV protease (100 nM final concentration) in PBS buffer was treated with a custom-ordered TEV protease substrate, **DABCYL-ENLYFQSGTK-5-FAM**, from GenScript. TEV substrate was prepared as 1 µM aliquot in DMSO, added to TEV protease samples at 50 nM final concentration, and detected at $\lambda_{ex}$ = 495 nm and $\lambda e_m$ = 550 nm. Activity assay is read on a BioTek Plate reader immediately after mixing TEV protease samples (first treated with DMSO or compounds at indicated concentrations for 1h) with the substrate in PBS buffer supplemented with 333 mM citrate and 5 mM DTT.

*DataWarrior SAR analysis (SALI Plots)*

Rescreen and counter-screen results were analyzed to determine the structure-activity relationship (SAR) using DataWarrior[2]. The linear region of progress curves from re-screen and counter-screen were used to calculate slopes for each treatment. The percent activity of caspase-10 was calculated to the DMSO control slopes. A list of generated percent activity values and SMILES corresponding to each compound were saved in a .csv file (**Table S1**). The generated file was opened with DataWarrior (v6.1.0). Data was decomposed to R-groups before running SAR analysis. The R-group decomposition chosen for our analysis was the “Most central ring system” to generate a new column of similar R-groups for our hit list. Similarity analysis using the structure-activity landscape index (SALI) was then used to generate the clustering SAR analysis based on the most central ring system.

*Cellular thermal shift assay (CETSA)*

Freshly lysed Jurkat cell lysates were spun down at 12,000 x *g* and 4°C for 10 min to remove any insoluble cell debris, then diluted to a final concentration of 3.0 mg/mL. These lysates were then labeled with either DMSO or compound (100 µM) for 1h at ambient temperature. Next, in 8 PCR tubes, DMSO treated lysates were added (50 µL each). In a different set of 8 tubes, compound-treated lysates (50 µL) were added to each. These lysates were then heated using a PCR block at the specified temperature for 3 min, immediately followed by an indefinite hold at 4°C. After heating, the lysates were transferred back to 1.5 mL microcentrifuge tubes and centrifuged at 21,100 x *g* and 4°C for 45 min. After centrifugation, 30 µL of the soluble portion of each sample was carefully pipetted into fresh microcentrifuge tubes, and 10 µL of 4x loading dye (Bio-Rad) was added. Samples were resolved by SDS-PAGE (Bio-Rad 4 - 20% Criterion Stain-Free™), and the gel was imaged using Bio-Rad Chemidoc Stain-Free setting for protein loading. The gel was then transferred to a nitrocellulose membrane (Bio-Rad) and blocked in 5% (w/v) milk in TBS (Tris-buffered saline) for 60 min at ambient conditions. Membranes were incubated with primary antibodies in 5% (w/v) milk in TBS overnight (16 - 18 hrs) at 4°C. Blots were then washed 3 times with TBS for 5 min. Membranes were then incubated with secondary antibodies (Goat anti-Rabbit IgG, DyLight 647 and DyLight 800; VWR, 102673-328 and 102673-412) in 5% (w/v) milk in TBST (Tris-buffered saline with 0.1% Tween20) for 1h at room temperature and washed 3 times with TBS. The membrane was then imaged using a Bio-Rad ChemiDoc imager. The primary antibodies and dilutions used are as follows: anti-CASP3 (Cell Signaling, 9662, 1:1,000), anti-caspase-8 (cleaved form, Cell Signaling, 9746, 1:1,000), anti-caspase-8 (pro-form, Cell Signaling, 4970, 1:1,000), anti-actin (Cell Signaling, 3700, 1:5,000), anti-caspase-2 (ProteinTech, 10436-1-AP, 1:1,000), and anti-caspase-9 (Cell Signaling, 9502, 1:1,000), anti-caspase-10 (MBL Life Science, M059-3, 1:1000).

*Confocal Microscopy.*

Sterilized coverslips were placed in each well of a 24-well plate and coated with poly-d-lysine for 30 minutes at 37°C, then washed 3 times with sterile water and allowed to dry. ΔΔG3BP1/2 KO U2OS cells stably expressing GFP-G3BP1 were seeded on coverslips overnight (80,000 cells/well) and then treated with the indicated compounds. Following treatment, media was aspirated, and each well was washed 1 time with 500 µL DPBS, followed by fixation with 3.7% formaldehyde in DPBS for 15 minutes at room temperature. Each well was then washed 2 times with 500 µL DPBS and permeabilized with 500 µL 0.1% TritonX in DPBS for 6 minutes at room temperature. Cells were then washed 2x with 500 µL of DPBS and blocked in 1% FBS in DPBS for 30 min at ambient conditions. Cells were then incubated with 500 µL 1 µg/mL DAPI stain in PBS for 5 minutes. Cells were then washed 2 times with 500 µL PBS and mounted onto glass slides using Aqua-Poly/Mount mounting media (Polysciences, Inc., 18606-20). Samples were left to sit in the dark for 24h. Slides were then imaged on a Zeiss LSM880 confocal microscope at 63X oil objective with 2X manual zoom.

*Compound treatment and biotinylation*

Competitive chemoproteomic analysis was performed as reported previously[5]. In brief, Jurkat lysates were prepared by Ultrasonic Probe Sonicator lysis (Power 2, 10x pulses; 1-second pulse, 1 second off) on ice. Lysate concentrations were then adjusted to 2 mg/mL) and 200 µL lysates (2 mg/mL) were treated with the pifithrin-µ (10 µM) or vehicle (DMSO) for 1h at ambient temperature. Samples were then labeled with 200 µM iodoacetamide alkyne (IAA) for 1h at ambient temperature. Samples were subjected to bioorthogonal copper(I)-catalyzed azide-alkyne cycloaddition (CuAAC)[11] or "click" conjugation to isotopically differentiated "light-" and "heavy-" biotin-azide. The samples were treated with a premixed cocktail of click reagents consisting of biotin-azide tags (4 µl of 5 mM stock), TCEP (4 µL of fresh 50 mM stock in water), TBTA (12 µl of 1.7 mM stock in DMSO/t-butanol 1:4), and $CuSO_4$ (4 µl of 50 mM stock in water). After 1h, the samples were then combined pairwise (400 µL total) and treated with 40 µL of 10% SDS (1% SDS final) followed by 0.5 µL of benzonase (Fisher Scientific, 707464). Samples were left to incubate for 30 min at 37°C. Following benzonase treatment, samples were subjected to Single-Pot Solid-Phase-enhanced sample preparation (SP3), as reported previously[6-9].

*Single-Pot Solid-Phase-enhanced Sample-Preparation (SP3)*

Following the previously reported protocol, for each 400 µL sample (400 µg protein), 40 µL Sera-Mag SpeedBeadsTM Carboxyl Magnetic Beads, hydrophobic (Thermo ScientificTM, 09-981-123) and 40 µL Sera-Mag SpeedBeadsTM Carboxyl Magnetic Beads, hydrophilic (Thermo ScientificTM, 09-981-121) were gently mixed and washed with 1 mL distilled water. Beads were combined using a magnetic rack (Sergi Lab Supplies, 1005a), and water was carefully aspirated. Washes were repeated 3x. Mixed beads were resuspended in 80 µL of double distilled water and then added to the 400 µL of combined samples. The bead-sample mixture was then incubated for 5 min at ambient conditions with shaking (1000 rpm). 200 proof ethanol was added to each sample such that each sample contained ≥55% ethanol by volume (for 480 µL of combined sample/SP3 beads, 600 µL of ethanol was added). The samples were incubated for 5 min at ambient conditions with shaking (1000 rpm). The beads were then washed three times with 600 µL 80 % ethanol in water. After washes, ethanol was removed using the magnetic rack, and beads were then resuspended in 200 µL 0.5 % SDS in PBS containing 2 M urea. 10 µL of 200 mM DTT (10 µM final concentration) was then added to each sample, and the samples were incubated at 65°C for 15 min. Following reduction, 10 µL of 400 mM iodoacetamide (20 µM final concentration) was added to each sample, and the samples were incubated for 30 min at 37°C with shaking at 300 rpm. Subsequently, 600 µL of 200-proof ethanol was added to each sample, and the samples were incubated for 5 min at ambient conditions with shaking (500 rpm). Beads were then washed three times with 80 % ethanol in water. Samples were then diluted with 150 µL 2 M urea in PBS, followed by the addition of reconstituted MS-grade trypsin (2 µg, Promega, V5111) and chymotrypsin (1 µg, ThermoFisher, PI90056). The samples were subjected to water bath sonication for 1 min and subsequently left to digest overnight (16 - 18hr) at 37oC and shaking at 200 rpm. The digested peptide solution and SP3 beads were then transferred into 15 mL falcon tubes. Peptides were then re-bound to SP3 beads by adding 3.8 mL of 100% acetonitrile for a final percentage of ≥95% acetonitrile by volume. The peptides were then shaken at 1000 rpm for 10 min at ambient temperature. Beads were collected using a magnet, and the solution was discarded. Samples were washed with 1 mL of 100% acetonitrile three times. Digested peptides were then eluted with 100 µL of 2% DMSO in water, shaking at 1000 rpm for 30 min at 37°C. The supernatant was collected on ice in a 1.5 mL centrifuge tube after separating SP3 beads with a magnetic rack. SP3 beads were resuspended with an additional 100 µL of 2% DMSO in water, shaking at 1000 rpm for 45 min at 37°C. The supernatant was collected after separating SP3 beads with a magnetic rack and combined with the previous elution volume (200 µL total).

*Streptavidin Enrichment*

PierceTM Streptavidin Agarose resin (Thermo ScientificTM, PI20353) (100 µL of resin) was first washed for a total of 3 times in 10 mL of PBS followed by centrifugation at 1,800 x g, 3 min each wash. Following the final wash with PBS, the buffer was carefully aspirated, ensuring it did not disturb spun-down agarose resin. The samples were then resuspended in 1 mL PBS in a 1.5 mL microcentrifuge tube. The 200 µL peptide elution from previously prepared SP3 clean-up and digestion method was added directly to the 1 mL of streptavidin resin in PBS. The samples were left rotating for 2h at ambient conditions. Following enrichment, the samples were spun down by centrifugation at 1,400 x *g* for 5 min, and the supernatant was aspirated and discarded. The streptavidin agarose was then subjected to washes, 2x with 1 mL of PBS and 2x with 1 mL of water. After each wash, the samples were spun down by centrifugation at 1,400 x g for 5 min.

*MS Sample Clean-up*

Following the manufacturer's protocol, the eluted peptides were subjected to buffer exchange using protein desalting columns, PierceTM C18 100 µL Tips (Thermo ScientificTM, 87784). First, 10 mL of the following four solutions were prepared: A) 100% acetonitrile (MeCN), B) 50:50 MeCN: ultra-pure water mixture, C) Ultrapure water with 0.1% trifluoroacetic acid, and D) 60% MeCN with 0.1% trifluoroacetic acid prepared in ultrapure water. MS sample clean-up was started by equilibrating one Pierce$^{TM}$ C18 100 µL Tip with 100 µL of solution A. This was repeated a total of two times while not allowing the tips to dry between washes. After equilibration, the tips were washed with 100 µL of solution B 2x. The final washing step was using 100 µL of solution C and repeating this wash step three times. Once the tips were washed with solution C, 100 µL of samples were loaded into the tips, making sure to pipette up/down at least 10x for higher peptide binding efficiency. Once peptides were bound, the tips were washed for a total of 2x with solution C. Finally, elution of peptides proceeded with 100 µL of solution D. Following MS sample clean-up, the 100 µL samples were completely dried by speedvac and the resulting dried peptides were then reconstituted in 20 µL of a solvent composed of 5% acetonitrile and 1% formic acid in ultrapure water.

Proteomics acquisition

The prepared mass spectrometry samples were analyzed using liquid chromatography-tandem mass spectrometry (LC-MS/MS) with a Thermo Scientific$^{TM}$ Orbitrap Eclipse$^{TM}$ Tribrid$^{TM}$ mass spectrometer (Thermo Scientific$^{TM}$) coupled to an Easy-nLC™ 1200 pump and to a High Field Asymmetric Waveform Ion Mobility Spectrometry (FAIMS) Interface, stated when used. The peptides were fractionated using a C18 reversed phase resin packed in-house and prepared using a 16 cm long, 100 µM inner diameter (ID) fused silica capillary (particle size, 1.9 µm; pore size, 100 Å; Dr. Maisch GmbH). An 80-minute water-acetonitrile gradient was delivered using the EASY-nLCTM 1200 system at varying flow rates (Buffer A: water with 3% DMSO and 0.1% formic acid and Buffer B: 80% acetonitrile with 3% DMSO and 0.1% formic acid). The detailed 80-minute gradient includes 0 – 5 min from 1 % to 15 % at 220 nL/min, 3 – 63 min from 15 % to 45 % at 220 nL/min, 63 – 73 min at 45% to 55% at 220nL/min, 73-74 min from 55 % to 95 % at 250 nL/min, and 74 to 80 min at 95% buffer B in buffer A. Data was collected with charge exclusion (1≥8). The data was acquired using a Data-Dependent Acquisition (DDA) method consisting of a full orbitrap MS1 scan (Resolution = 120,000) followed by sequential MS2 scans (Resolution = 15,000) to utilize the remainder of the 1-second cycle time. The precursor isolation window was set to 1 m/z and high energy c-trap dissociation (HCD) normalized collision energy was set to 30%. The run time was 80 minutes, and the injection volume was 5 µL. Data acquired with the FAIMS device utilized 3 compensation voltages (CV; -35, -45, -55V) as used in our previous study[6].

*Data analysis*

The proteomic workflow for the experiments was analyzed using FragPipe with initial tool parameters set as default. FragPipe output data was compiled using in-house Python scripts. Custom python scripts compiled modified_peptide_label_quant.tsv (quantification) outputs from FragPipe (v21.0). As a preprocessing step, the logged ratios from singleton peptides (ratios that come from peptides that are unpaired) were removed before further analysis. Unpaired heavy- or light-identified peptides remained by setting ratios to $\log_2(20)$ or $\log_2(1/20)$. Median heavy-over-light ratios were computed for the same cysteine residue from cysteine peptides of different charges and missed cleavages in the same replicate. The average of medians was calculated to obtain a final "average_of_medians" metric. This "average_of_medians" value was utilized to compare the isotopical quantification of the same identified peptides, modified cysteine residues and proteins across replicates within the same experiment.

*Protein, peptide, and cysteine identification*:
The MS RAW files were searched with MSFragger (v4.0) and FragPipe (v21.0). Precursor and fragment mass tolerance was set as 20 ppm. Missed cleavages were allowed up to 1. Peptide length was set to 7 – 50, and peptide mass range was set to 500 - 5000. For identification, cysteine residues were searched with differential modification C+. For ligandability quantification, MS1 labeling quant was enabled with Light set as C+463.2366 and Heavy set as C+469.2742. MS1 intensity ratio of heavy and light labeled cysteine peptides were reported with Ionquant (v1.8.10). Calibrated and deisotoped spectrum files produced by FragPipe were reused for further analysis. The MS search data have been deposited to the ProteomeXchange Consortium via the PRIDE partner repository with the dataset identifiers PXD053315. File details can be found in **Table S2**. Custom python scripts were implemented to compile labeled peptide datasets. Unique proteins, unique cysteines, and unique peptides were quantified for each dataset. Unique proteins were established based on UniProt protein IDs. Unique peptides were found based on sequences containing a modified cysteine residue. Unique cysteines were classified by an identifier consisting of a UniProt protein ID and the amino acid number of the modified cysteine (ProteinID_C#); residue numbers were found by aligning the peptide sequence to the corresponding UniProt protein sequences found in protein.fas FragPipe output files. A new identifier for each modified cysteine residue number was created for cases where multiple cysteines were labeled on the same peptide. Multiplexed peptide identifiers were reported as ProteinID_C#1 and ProteinID_C#2, instead of ProteinID_C#1_C#2.

*Cell Titer Glo ® Cell viability analysis*
Using a multichannel pipette, jurkat cells in complete RPMI media (100 µL of 1.0 x $10^6$ cells/mL) were added to a 96-well white/clear bottom tissue culture-treated plate. To each well, samples were treated with the indicated compounds at the indicated concentrations or vehicle (DMSO). Following 1h incubation, mega FasL (AdipoGen; AG-40B-0130-C010) was Cell viability assay (Promega, G9242) was added to each well, and the relative luminescence (RLU) was measured. Percent cell viability was calculated as reactive to the DMSO-treated Jurkat cells. Samples were analyzed in three biological replicates.

*Ellman's Assay*
10mM stock Ellman's Reagent solution was prepared by dissolving 4 mg of 5,5′-dithiobis-(2-nitrobenzoic acid) (DTNB) in 1 mL of Reaction Buffer (0.1M sodium phosphate, pH 8.0, containing 1mM EDTA). Reduced GSH standards (0, 31.25, 62.5, 125, 250 µM) were prepared using the same Reaction Buffer. In a 96-well plate, 2.5 µL of the 10mM Ellman's reagent was added to each 200 uL of the GSH standards, and the plates were incubated for 15 mins at room temperature in the dark. Next, the absorbance of the resultant yellow-colored 5-thio-2-nitrobenzoic acid (TNB) product was measured at 412 nm using a Tecan Infinite F500 microplate reader. The data was used to generate a standard curve. For the samples, the indicated compounds were added to

200uL of 125 µM GSH (to the specified final concentrations) and incubated at room temperature for 1 h. Next, 2.5 µL of the 10mM Ellman's reagent was added to this mix and incubated for 15 mins at room temperature. Next, the absorbance of the resultant yellow-colored TNB product was measured at 412 nm using a Tecan Infinite F500 microplate reader. The GSH consumption for each of the indicated compounds was determined from the standard curve.

*Statistics*
Statistical significance was calculated with an unpaired two-tailed Student's t-test using GraphPad Prism 10(version 10.3.0. GraphPad Software Inc., La Jolla). Data are shown as mean values and standard deviations for replicates ranging from n = 2 to 16, as stated for each experiment. P values of < 0.05 were considered significant.

*Z'-factor and z-score calculations*
Z-prime (Z'-factor) statistic was used to measure assay quality, showing the separation between the distribution of the positive control (DMSO) and the negative controls (noTEV treatment and **KB7** treatment). Z'-factor was calculated per plate as follows:

$$Z'\, factor \;=\; 1 - \frac{3\,x\,(\sigma p \;-\; \sigma n)}{|\mu p \;-\; \mu n|}$$

σ = standard deviation
µ = mean
p = positive controls (DMSO)
n = negative controls

The standard score or z-score is a measure of standard deviations from the mean for each readout value. z-scores were calculated to the positive controls of each plate (DMSO) for each readout. z-score was calculated as follows:

$$Z - score \;=\; \frac{sample \;-\; DMSO\; control\; mean}{standard\; deviation\; of\; DMSO\; control}$$

### (E) Chemistry Methods

**General Methods**. All reactions were performed in dried, clean glassware. Silica gel P60 (SiliCycle) was used for normal phase column chromatography. Thin layer chromatography (TLC) plates were visualized by fluorescence quenching under UV light or by staining with iodine, $KMnO_4$, or bromocresol green. Reagents were purchased from Sigma-Aldrich (St. Louis, MO), Alfa Aesar (Ward Hill, MA), EMD Millipore (Billerica, MA), Fisher Scientific (Hampton, NH), Oakwood Chemical (West Columbia, SC), Combi-blocks (San Diego, CA) and Cayman Chemical (Ann Arbor, MI) and used without further purification. Additionally, all isotopically enriched reagents were purchased from Sigma-Aldrich (St. Louis, MO) or Cambridge Isotope Laboratories (Cambridge, MA) and used without further purification. $^{1}H$ NMR and $^{13}C$ NMR spectra for characterization of novel compounds and monitoring reactions were collected in $CDCl_3$, $CD_3OD$, $D_2O$, or DMSO-$d_6$ (Cambridge Isotope Laboratories, Cambridge, MA) on a Bruker AV 400 MHz spectrometer in the Department of Chemistry & Biochemistry at The University of California, Los Angeles. All chemical shifts are reported in the standard notation of parts per million using the peak of residual proton signals of the deuterated solvent as an internal reference. Coupling constant (*J*) units are in Hertz (Hz). Splitting patterns are indicated as follows: br, broad; s, singlet; d, doublet; t, triplet; q, quartet; m, multiplet; dd, doublet of doublets; dt, doublet of triplets. Low-resolution mass spectrometry was performed on an Agilent Technologies InfinitiyLab LC/MSD single quadrupole LC/MS (ESI source). High-resolution mass spectrometry was performed on a Waters LCT Premier with ACQUITY LC and autosampler (ESI source).

**Synthesis of *tert*-butyl 4-(1*H*-imidazole-1-carbonothioyl)piperazine-1-carboxylate (1)**

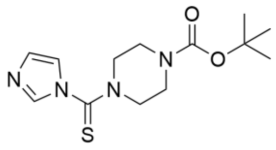

To a stirred solution of 1,1'-thiocarbonyldiimidazole (1.30 g, 7.50 mmol, 1.0 eq.) in $CH_2Cl_2$ (200 mL) was added *N*-Boc-piperazine (1.40 g, 7.50 mmol, 1.0 equiv.) and the reaction mixture was stirred overnight at room temperature (RT). The reaction mixture was quenched with water, dried ($Na_2SO_4$), filtered, and concentrated *in vacuo* to afford *tert*-butyl 4-(1*H*-imidazole-1-carbonothioyl)piperazine-1-carboxylate (**1**) (1.83 g, 6.17 mmol, 82 %) as a white solid : $^1$H NMR (400 MHz, $CDCl_3$) $\delta_H$ 7.86 (1 H, s), 7.18 (1 H, s), 7.08 (1 H, s), 3.95 – 3.78 (4 H, m), 3.63 – 3.41 (4 H, m), 1.46 (9 H, s); $^{13}$C NMR (101 MHz, $CDCl_3$) $\delta_C$ 179.3, 154.3, 137.4, 130.2, 119.2, 81.0, 51.4, 28.3; LRMS *m/z* (ESI$^+$) 297.1 [M+H]$^+$.

**Synthesis of *tert*-butyl 4-(2-hydrazineylethanethioyl)piperazine-1-carboxylate (2)**

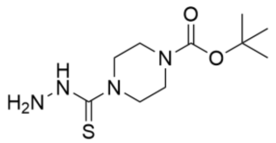

To a stirred solution of *tert*-butyl 4-(1*H*-imidazole-1-carbonothioyl)piperazine-1-carboxylate (**1**) (1.60 g, 5.40 mmol, 1.0 eq.) in EtOH (30 mL) was slowly added hydrazine monohydrate (307 µL, 5.94 mmol, 1.1 eq.). The reaction mixture was stirred at RT for 5 min then heated under reflux for 3 h. The resultant off-white precipitate was filtered, washed with *tert*-butyl methyl ether, and dried by suction to afford *tert*-butyl 4-(2-hydrazineylethanethioyl)piperazine-1-carboxylate (**2**) (670 mg, 2.57 mmol, 47%) as an off-white solid : $^1$H NMR (400 MHz, DMSO-$d_6$) δ 3.71 (4 H, dd, *J* 6.2, 4.3), 3.32 (4 H, d, *J* 10.5), 1.40 (9 H, s); $^{13}$C NMR (101 MHz, DMSO-$d_6$) δ 183.0, 154.3, 79.6, 47.4, 28.5; LRMS *m/z* (ESI$^-$) 259.1 [M-H]$^-$.

**Synthesis of *tert*-butyl 4-(5-phenyl-6H-1,3,4-thiadiazin-2-yl)piperazine-1-carboxylate (SO263)**

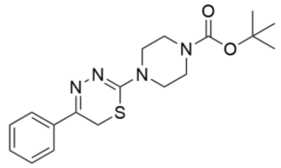

A solution of the 4-(2-hydrazineylethanethioyl)piperazine-1-carboxylate (**2**) (600 mg, 2.30 mmol, 1.0 eq.) and 2-bromoacetophenone (459 mg, 2.30 mmol, 1.0 eq.) in EtOH (10 mL) was stirred under reflux for 30 min. The hot reaction mixture was filtered, and dilute aq. $NH_3$ was added to the filtrate to achieve pH 8. The resulting precipitate was filtered and recrystallized in EtOH to afford *tert*-butyl 4-(5-phenyl-6*H*-1,3,4-thiadiazin-2-yl)piperazine-1-carboxylate (**SO263**) (180 mg, 499 mmol, 22%) as an off-white solid: $^1$H NMR (400 MHz, $CDCl_3$) $\delta_H$ 7.89 (2 H, s), 7.41 (4 H, d, *J* 7.1), 3.77 – 3.68 (4 H, m), 3.58 – 3.46 (6 H, m), 1.47 (9 H, s); $^{13}$C NMR (101 MHz, $CDCl_3$) $\delta_C$ 154.7, 151.7, 147.6, 135.5, 129.8, 128.7, 126.7, 80.3, 47.4, 28.4, 28.4, 22.7; LRMS *m/z* (ESI$^+$) 361.1 [M+H]$^+$.

**Synthesis of *tert*-butyl 4-(2-ethoxyacetyl)piperazine-1-carboxylate (3)**

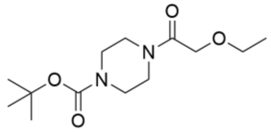

To a solution of *tert*-butyl piperazine-1-carboxylate (2.00 g, 10.7 mmol, 1.0 eq.) and DIPEA (4.50 mL, 26.0 mmol, 2.4 eq.) in $CH_2Cl_2$ (40 mL) was added 2-ethoxyacetyl chloride (1.41 mL, 12.9 mmol, 1.2 eq.) dropwise. The reaction mixture was stirred at RT for 2 h before being washed with 10% aq. $(NH_4)_2CO_3$ (2 × 40 mL) then sat. $NH_4Cl$ (2 × 40 mL). The organic component was dried ($Na_2SO_4$), filtered, concentrated *in vacuo* to afford *tert*-butyl 4-(2-ethoxyacetyl)piperazine-1-carboxylate (2.90 g, 10.7 mmol, 99%) as a yellow oil: $^1$H NMR (400 MHz, $CDCl_3$) $\delta_H$ 4.11 (2 H, s), 3.53 (4 H, t, *J* 7.0), 3.48 (2 H, d, *J* 12.8), 3.40 (4 H, s), 1.43 (9 H, s), 1.19 (3 H, t, *J* 7.0); $^{13}$C NMR (101 MHz, $CDCl_3$) $\delta_C$ 168.1, 154.6, 80.3, 70.4, 66.8, 45.1, 41.7, 28.4, 15.1; LRMS *m/z* (ESI$^+$) 186.1 [M+H]$^+$.

**Synthesis of 4-(2-ethoxyacetyl)piperazine-1-carbothiohydrazide (X)**

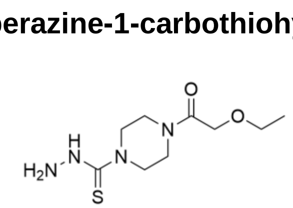

To *tert*-butyl 4-(2-ethoxyacetyl)piperazine-1-carboxylate (2.80 g, 10.3 mmol, 1.0 eq.) in $CH_2Cl_2$ (40 mL) was added 4 M HCl in dioxane (7.71 mL, 30.8 mmol, 3.0 eq.) dropwise. The reaction mixture was stirred at RT for 2 h before being concentrated *in vacuo* to afford the *tert*-butyl piperazine-1-carboxylate hydrochloride intermediate as an off-white solid. The residue was redissolved in $CH_2Cl_2$ (40 mL) and DIPEA (3.00 mL, 17.2 mmol, 1.7 eq.) was added followed by di(1*H*-imidazol-1-yl)methanethione (1.83 g, 10.3 mmol, 1.0 eq.). The reaction mixture stirred at RT for 18 h before being concentrated *in vacuo* to afford the 1*H*-imidazol intermediate as a yellow oil. The residue was redissolved in EtOH (40 mL) and hydrazine monohydrate (549 µL, 11.3 mmol, 1.1 eq.) was added and the reaction mixture was stirred under reflux for 3 h. The reaction mixture was cooled to rt and the volume concentrated to ca. 5 mL *in vacuo*. *tert*-Butyl methyl ether (40 mL) was added and the resultant precipitate was collected by centrifugation. The solid was washed with *tert*-butyl methyl ether (2 × 10 mL), collected by centrifugation, and dried *in vacuo* to afford 4-(2-ethoxyacetyl)piperazine-1-carbothiohydrazide (**X**) (686 mg, 2.78 mmol, 27%) as an off-white solid: $^{1}$H NMR (400 MHz, $CDCl_3$) $\delta_H$ 4.15 (2 H, d, *J* 2.08), 3.99 – 3.93 (2 H, m), 3.81 – 3.76 (2 H, m), 3.76 – 3.65 (4 H, m), 3.56 (2 H, q, *J* 7.03), 1.23 (3 H, t, *J* 7.03); $^{13}$C NMR; LRMS *m/z* (ESI$^+$) 247.1 $[M+H]^+$.

**Synthesis of 2-ethoxy-1-(4-(5-phenyl-6H-1,3,4-thiadiazin-2-yl)piperazin-1-yl)ethan-1-one (SO265) and 2-ethoxy-1-(4-(4-mercapto-5-phenyl-1*H*-pyrazol-3-yl)piperazin-1-yl)ethan-1-one (SO265i)**

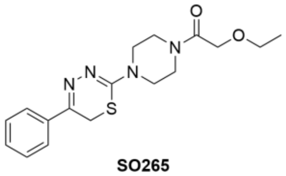


SO265

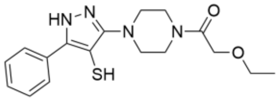


SO265i

A solution of 4-(2-ethoxyacetyl)piperazine-1-carbothiohydrazide (**X**) (300 mg, 1.22 mmol, 1.0 eq.) and 2-bromo-1-phenylethan-1-one (242 mg, 1.22 mmol, 1.0 eq.) in EtOH (10 mL) was stirred under reflux for 30 min. The reaction mixture was concentrated *in vacuo*, and H-NMR indicated an incomplete reaction. The residue was redissolved in EtOH (10 mL), further 2-bromo-1-phenylethan-1-one (121 mg, 0.61 mmol, 0.5 eq.) was added, and the reaction mixture was stirred under reflux for 30 min. The reaction mixture was concentrated *in vacuo* then redissolved in a ca. 1:1 MeOH and water (6 mL) solution before being purified by reverse phase column chromatography (elution with a gradient of 20 – 80% MeCN in water). Relevant fractions were dried by lyophilization to afford the isomer products. 2-Ethoxy-1-(4-(5-phenyl-6*H*-1,3,4-thiadiazin-2-yl)piperazin-1-yl)ethan-1-one (**SO265**) (49 mg, 0.14 mmol, 12%) was obtained as a pale yellow solid: R*f* 0.44 (EtOAc); $^{1}$H -NMR (400 MHz, DMSO-$d_6$) δ 7.93 (2 H, dd, *J* 8.13, 7.67), 7.50 – 7.43 (3 H, m), 4.15 (2 H, s), 3.77 (2 H, s), 3.72 – 3.63 (4 H, m), 3.57 – 3.52 (4 H, m), 3.49 (2 H, q, *J* 7.01), 1.14 (3 H, t, *J* 7.01); C-NMR (101 MHz, DMSO-$d_6$) δ 167.6, 153.6, 148.6,135.4, 130.1, 129.1, 127.1, 69.5, 66.3, 47.5, 44.4, 22.0, 15.5; LRMS *m/z* (ESI+) 347.1. 2-Ethoxy-1-(4-(4-mercapto-3-phenyl-1*H*-pyrazol-5-yl)piperazin-1-yl)ethan-1-one (**SO265i**) (131 mg, 378 µmol, 31%) was obtained as an orange oil: R*f* 0.00 (EtOAc); H-NMR (400 MHz, DMSO-$d_6$) $\delta_H$ 14.28 (1 H, br s), 9.34 – 8.75 (2 H, m), 7.63 – 7.48 (3 H, m), 4.40 (1 H, s), 4.17 (2 H, s), 4.03 – 3.82 (4 H, m), 3.78 – 3.59 (4 H, m), 3.51 – 3.41 (2 H, m), 1.17 – 1.07 (3 H, m); $^{13}$C -NMR (101 MHz, DMSO-$d_6$) $\delta_C$ 168.4, 162.8, 153.2, 132.8, 132.2, 129.6, 127.7, 69.4, 68.0, 66.3, 49.5, 40.4, 15.5; LRMS (ESI+) 347.1.

**Synthesis of 1,1'-((disulfanediylbis(5-phenyl-1*H*-pyrazole-4,3-diyl))bis(piperazine-4,1-diyl))bis(2-ethoxyethan-1-one) (SO265d)**

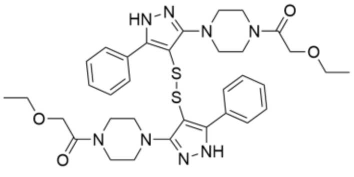

**SO265d**

A solution of 2-ethoxy-1-(4-(4-mercapto-3-phenyl-1*H*-pyrazol-5-yl)piperazin-1-yl)ethan-1-one (**SO265i**) in DMSO-$d_6$ (0.5 mL) was left at RT for 2 days. After which time, H-NMR analysis indicated that DMSO-mediated thiol oxidation had occurred, forming 1,1'-((disulfanediylbis(5-phenyl-1*H*-pyrazole-4,3-diyl))bis(piperazine-4,1-diyl))bis(2-ethoxyethan-1-one) **(SO265d)**: $^{1}$H-NMR (400 MHz, DMSO-$d_6$) $\delta_H$ (400 MHz, DMSO-*$d_6$*) 14.29 (2 H, br s), 7.82 – 7.75 (2 H, m), 7.73 – 7.67 (2 H, m), 7.56 – 7.42 (6 H, m), 4.20 – 4.10 (4 H, m), 3.81 – 3.49 (14 H, m), 3.49 – 3.45 (4 H, m), 3.45 – 3.39 (2 H, m), 1.14 – 1.08 (6 H, m); $^{13}$C -NMR (101 MHz, DMSO-*$d_6$*) $\delta_C$ 167.7, 157.5, 134.3, 129.7, 128.9, 126.0, 118.8, 68.7, 65.4, 49.0, 42.1, 14.6.

### (F) NMR Spectra

$^{1}H$ NMR of ***tert*-butyl 4-(1*H*-imidazole-1-carbonothioyl)piperazine-1-carboxylate (1)** in $CDCl_3$

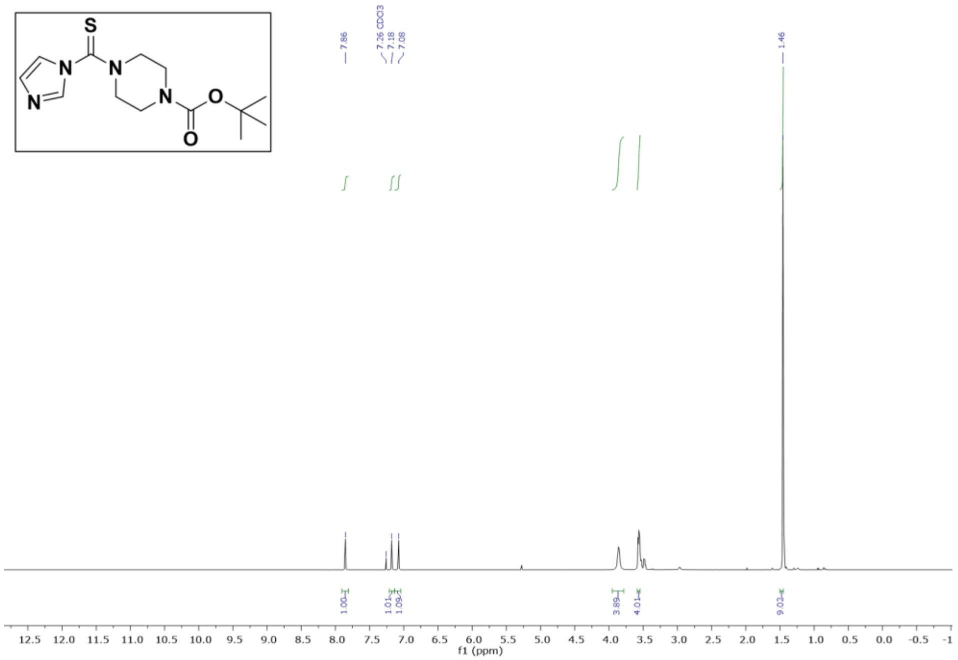


$^{13}C$ NMR of **tert-butyl 4-(1H-imidazole-1-carbonothioyl)piperazine-1-carboxylate (1)** in $CDCl_3$

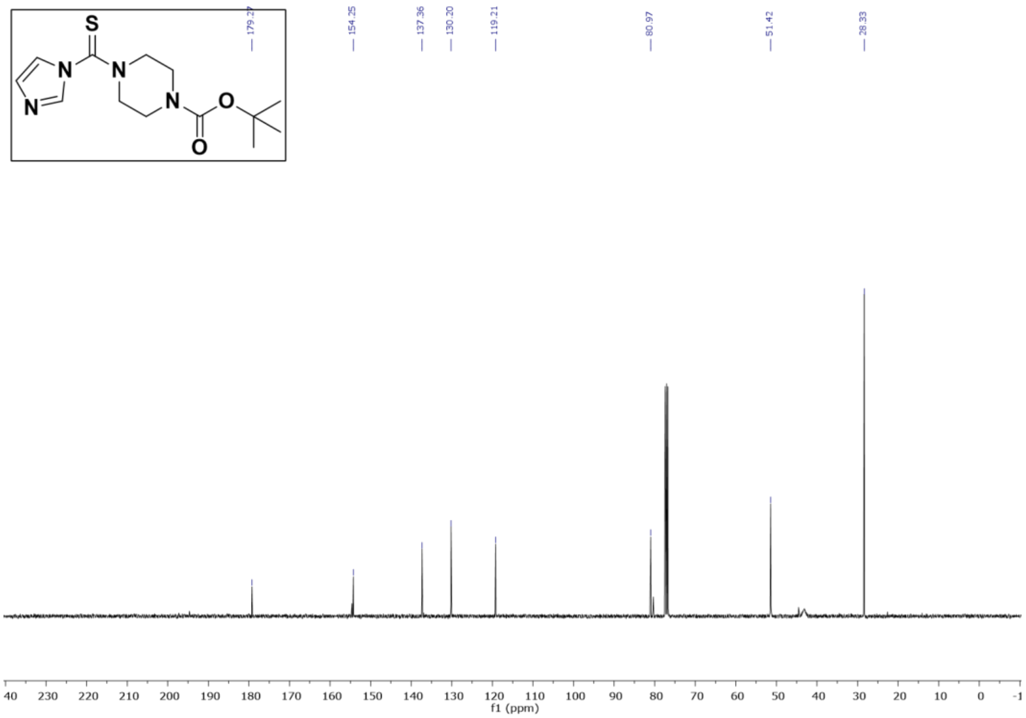


$^{1}H$ NMR of ***tert*-butyl 4-(2-hydrazineylethanethioyl)piperazine-1-carboxylate (2)** in DMSO-$d_6$

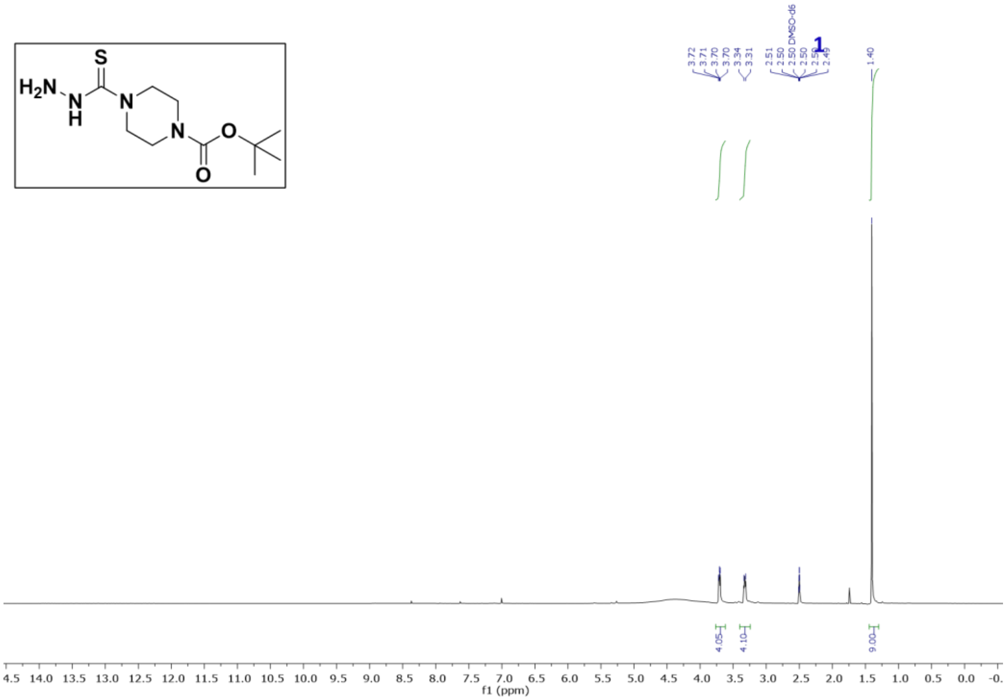


$^{13}C$ NMR of ***tert*-butyl 4-(2-hydrazineylethanethioyl)piperazine-1-carboxylate (2)** in DMSO-$d_6$

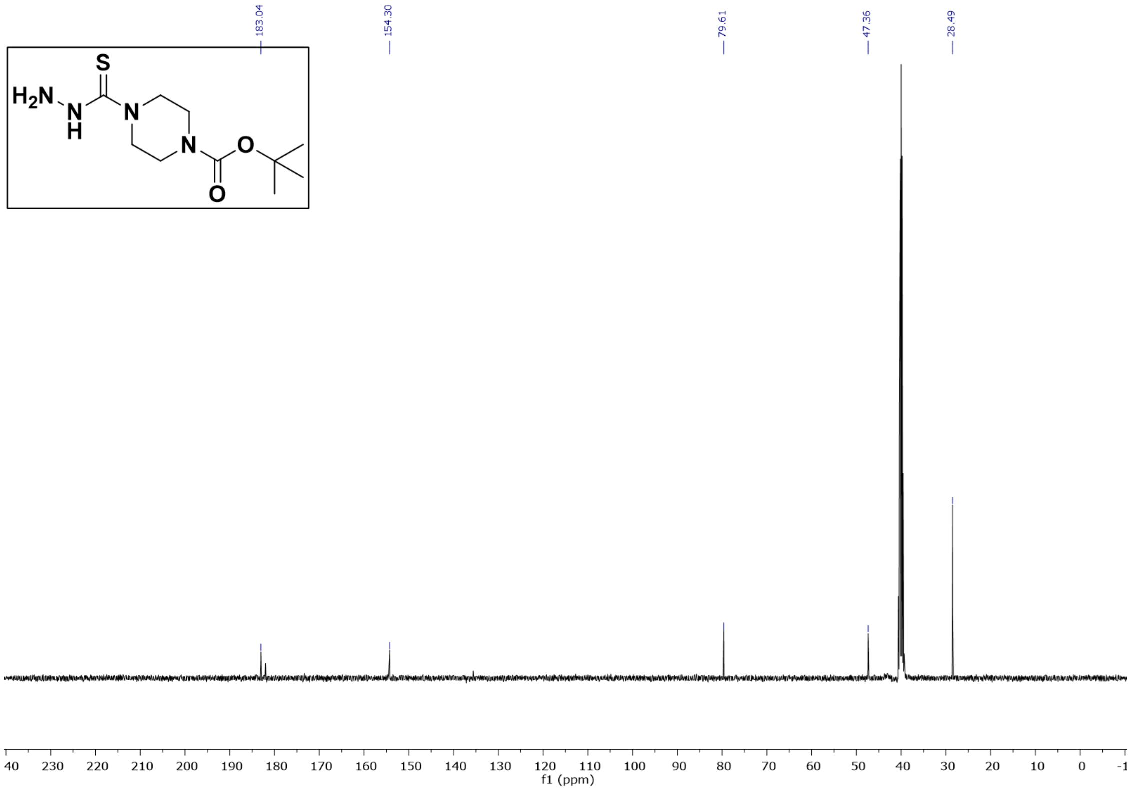


$^{1}H$ NMR of ***tert*-butyl 4-(5-phenyl-6H-1,3,4-thiadiazin-2-yl)piperazine-1-carboxylate (SO263)** in $CDCl_3$

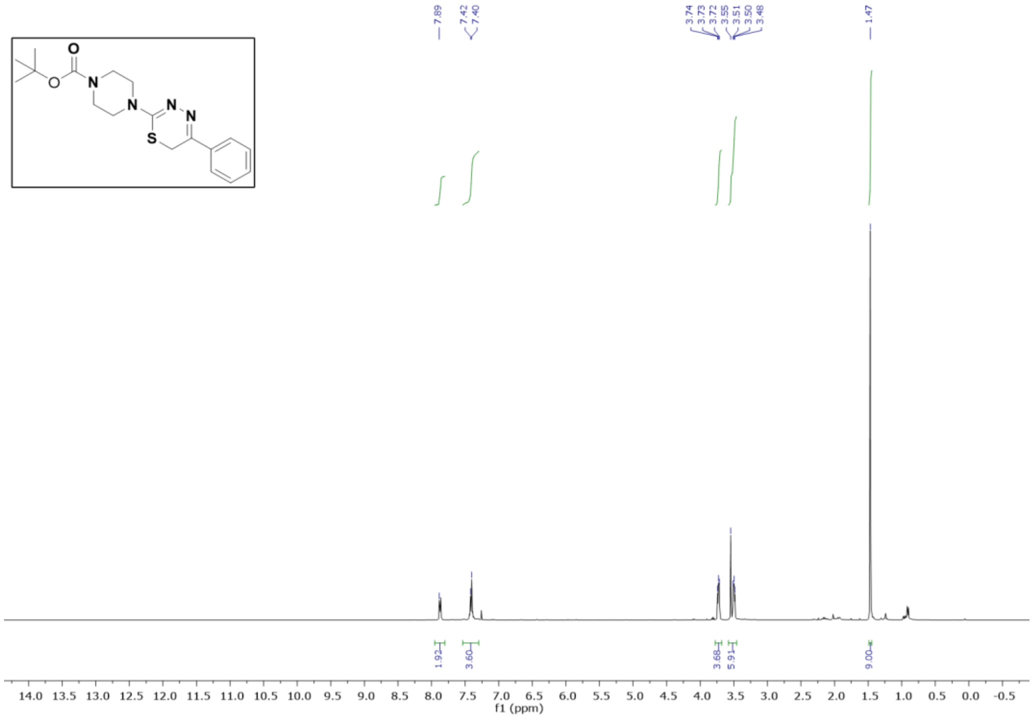


$^{13}$C NMR of ***tert*-butyl 4-(5-phenyl-6H-1,3,4-thiadiazin-2-yl)piperazine-1-carboxylate (SO263)** in $CDCl_3$

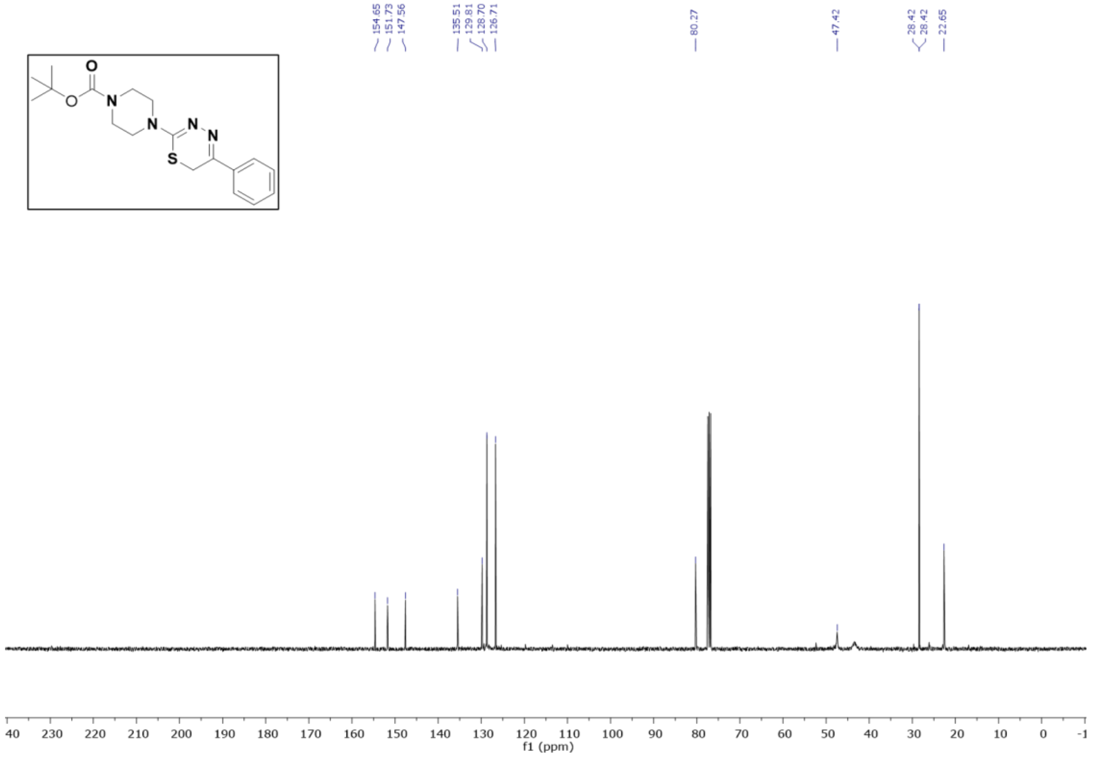


$^{1}$H NMR of ***tert*-butyl 4-(2-ethoxyacetyl)piperazine-1-carboxylate** in $CDCl_3$

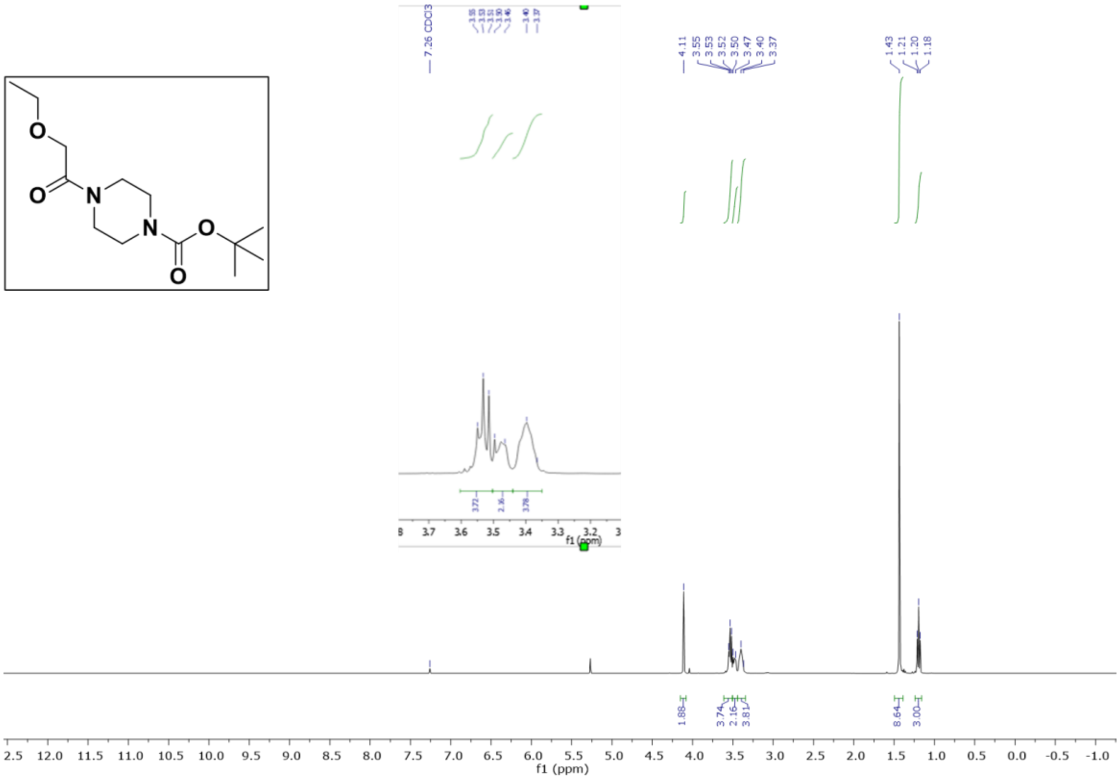


$^{13}C$ NMR of ***tert*-butyl 4-(2-ethoxyacetyl)piperazine-1-carboxylate** in $CDCl_3$

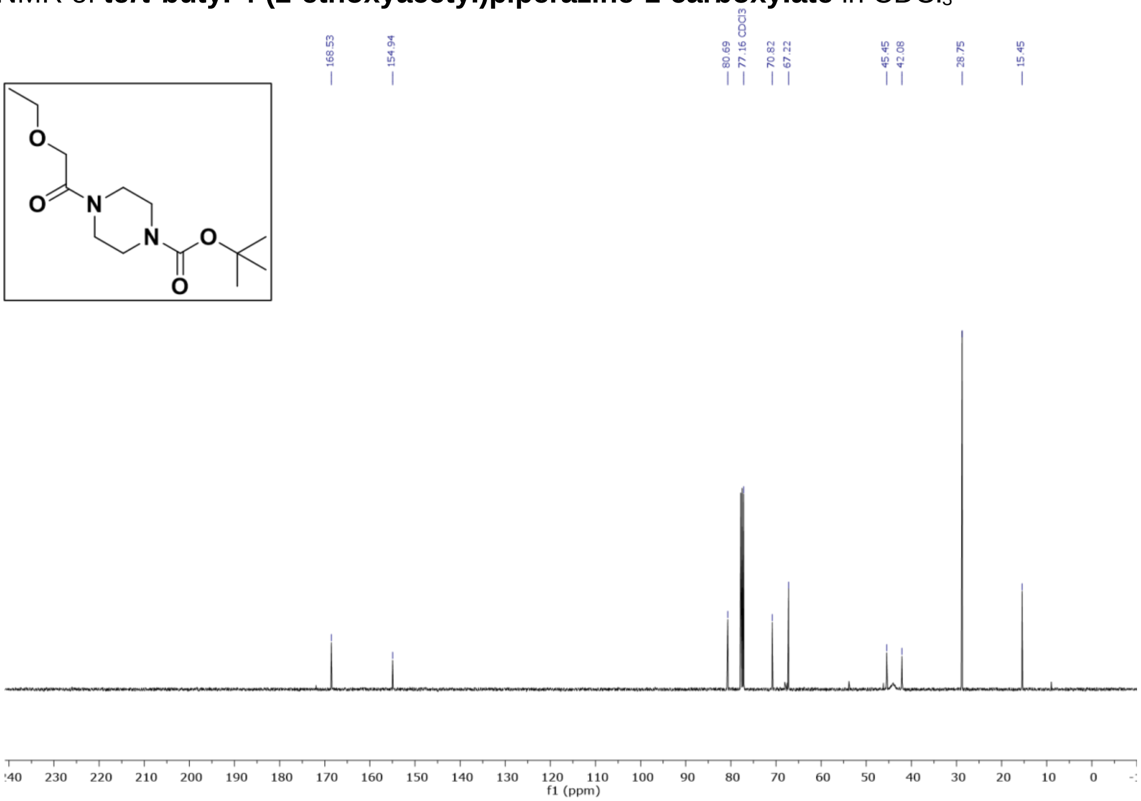


$^{1}H$ NMR of **4-(2-ethoxyacetyl)piperazine-1-carbothiohydrazide (X)** in $CDCl_3$

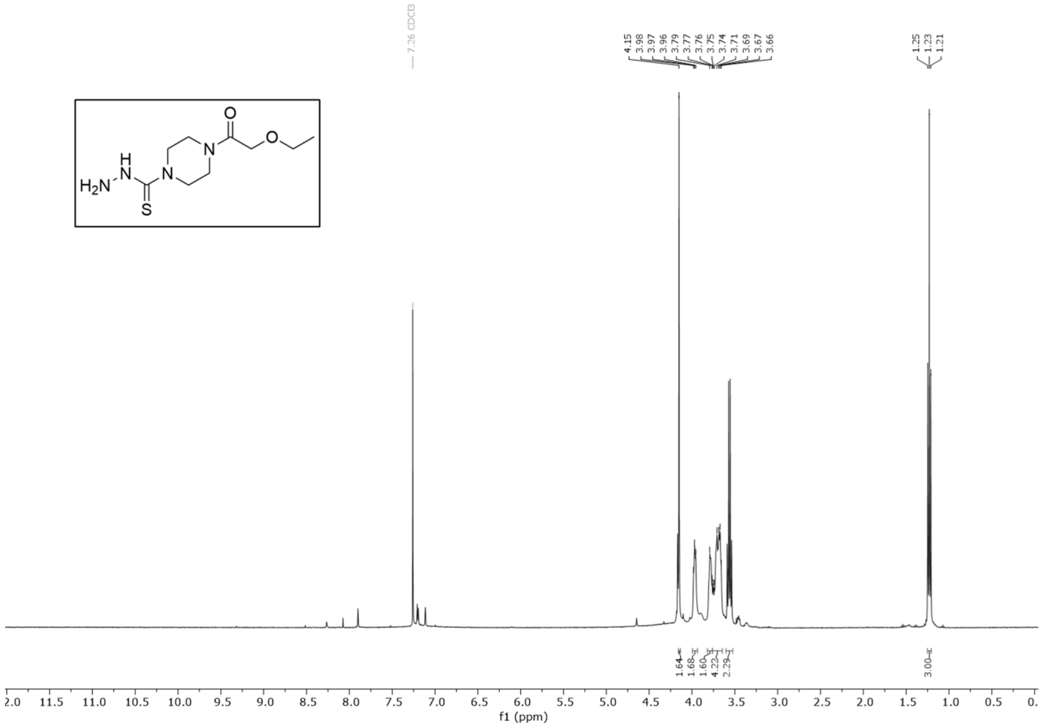


$^{13}$C NMR of **4-(2-ethoxyacetyl)piperazine-1-carbothiohydrazide (X)** in $CDCl_3$

$^{1}$H NMR of **2-ethoxy-1-(4-(5-phenyl-6H-1,3,4-thiadiazin-2-yl)piperazin-1-yl)ethan-1-one (SO265)** in DMSO-$d_6$

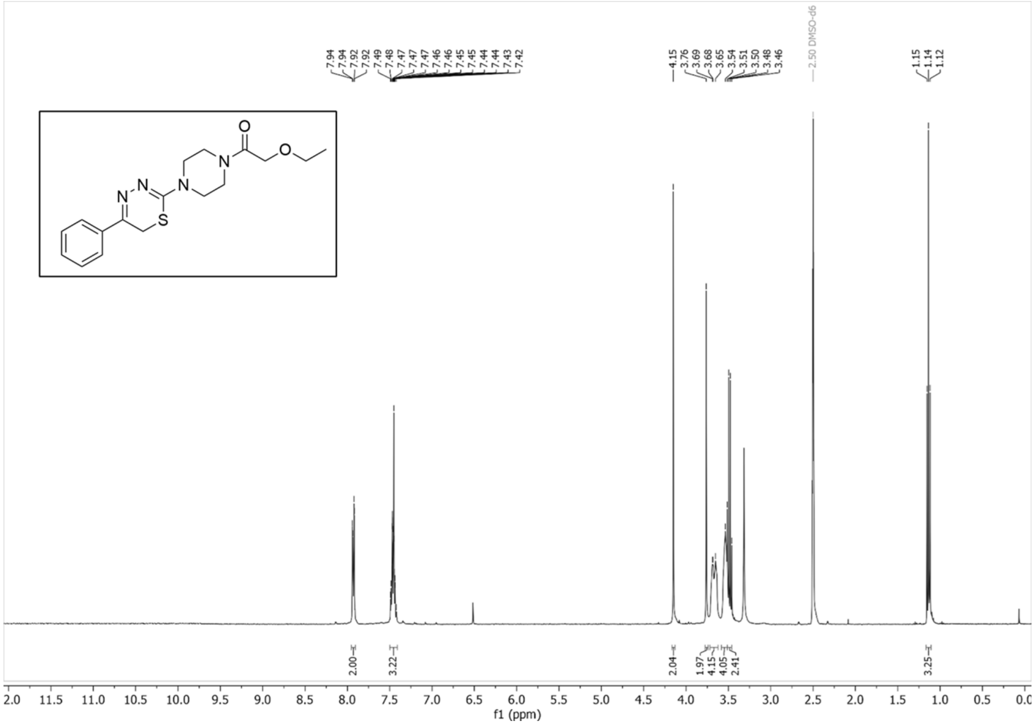


$^{13}C$ NMR of **2-ethoxy-1-(4-(5-phenyl-6H-1,3,4-thiadiazin-2-yl)piperazin-1-yl)ethan-1-one (SO265)** in DMSO

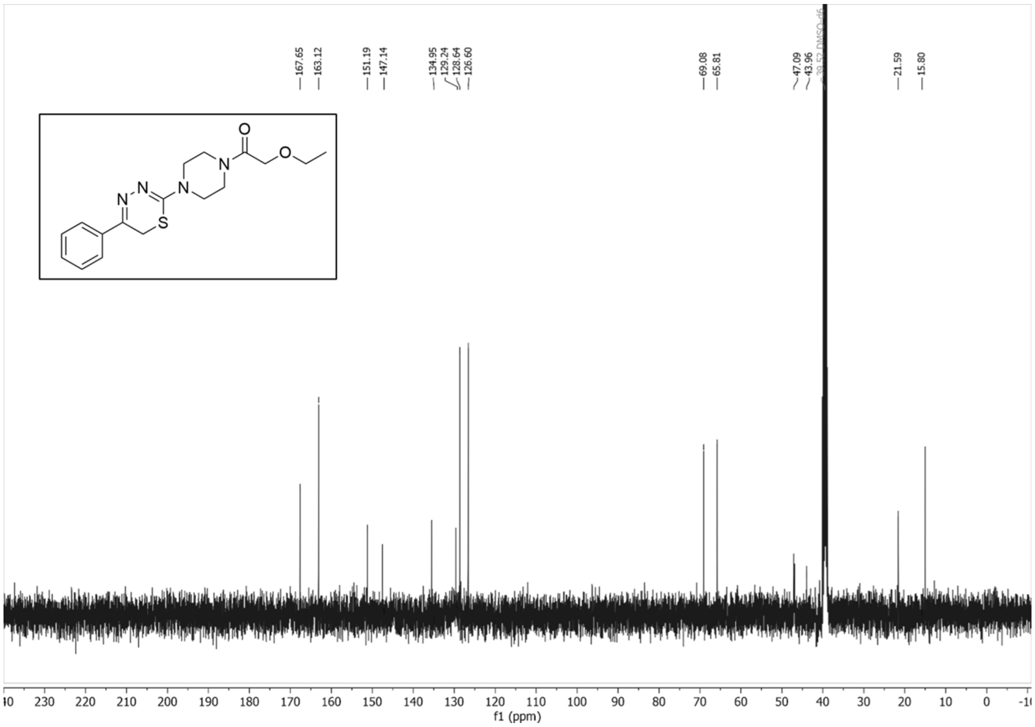


$^{1}H$ NMR of **2-ethoxy-1-(4-(4-mercapto-5-phenyl-1*H*-pyrazol-3-yl)piperazin-1-yl)ethan-1-one (SO265i)** in DMSO-$d_6$

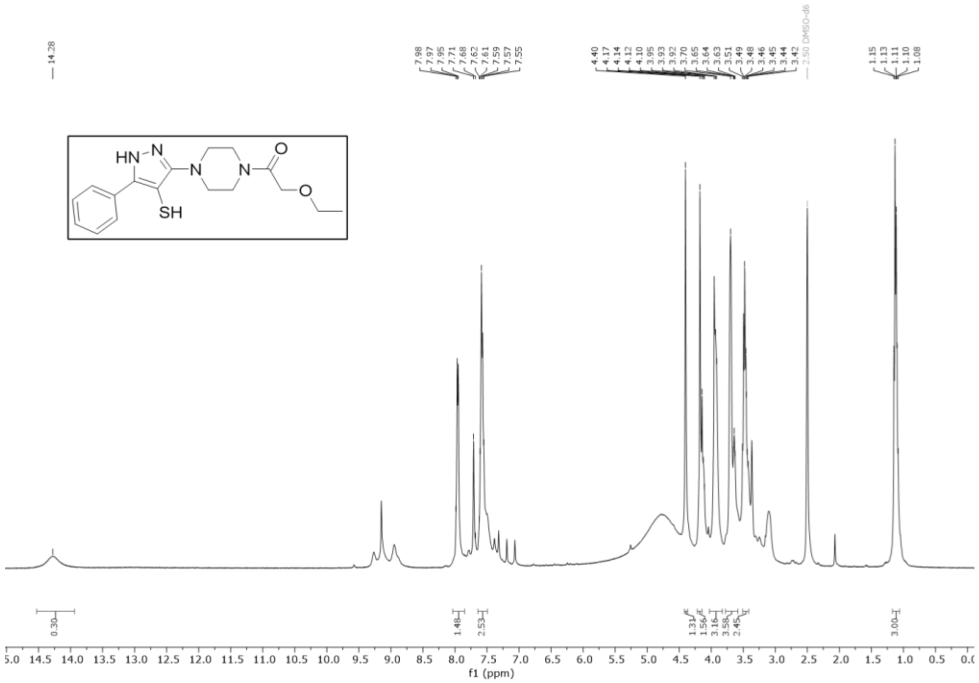


[13]C NMR of **2-ethoxy-1-(4-(4-mercapto-5-phenyl-1*H*-pyrazol-3-yl)piperazin-1-yl)ethan-1-one (SO265i)** in DMSO-$d_6$

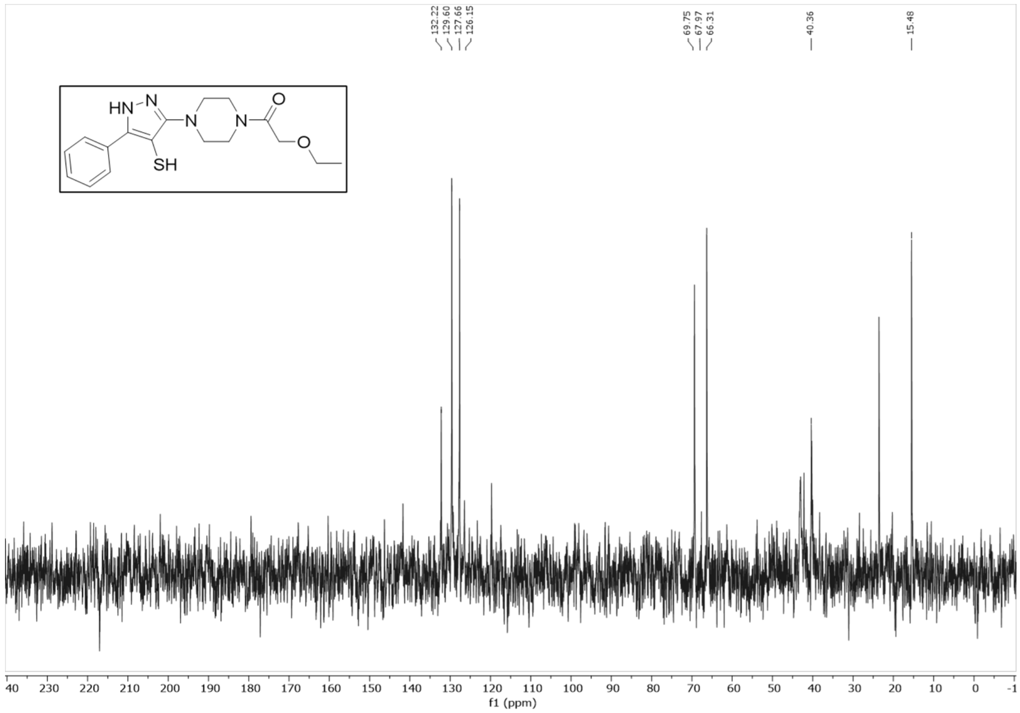


HMBC of **2-ethoxy-1-(4-(4-mercapto-5-phenyl-1*H*-pyrazol-3-yl)piperazin-1-yl)ethan-1-one (SO265i)** in DMSO-$d_6$

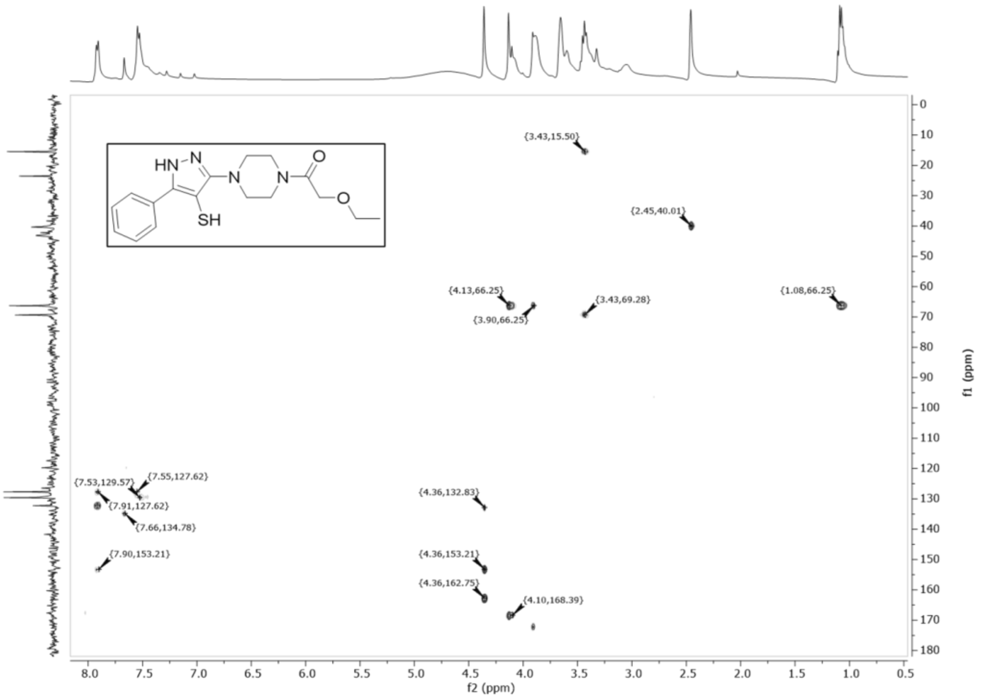


$^{1}$H NMR of **1,1'-((disulfanediylbis(5-phenyl-1*H*-pyrazole-4,3-diyl))bis(piperazine-4,1-diyl))bis(2-ethoxyethan-1-one) (SO265d)** in DMSO-$d_6$

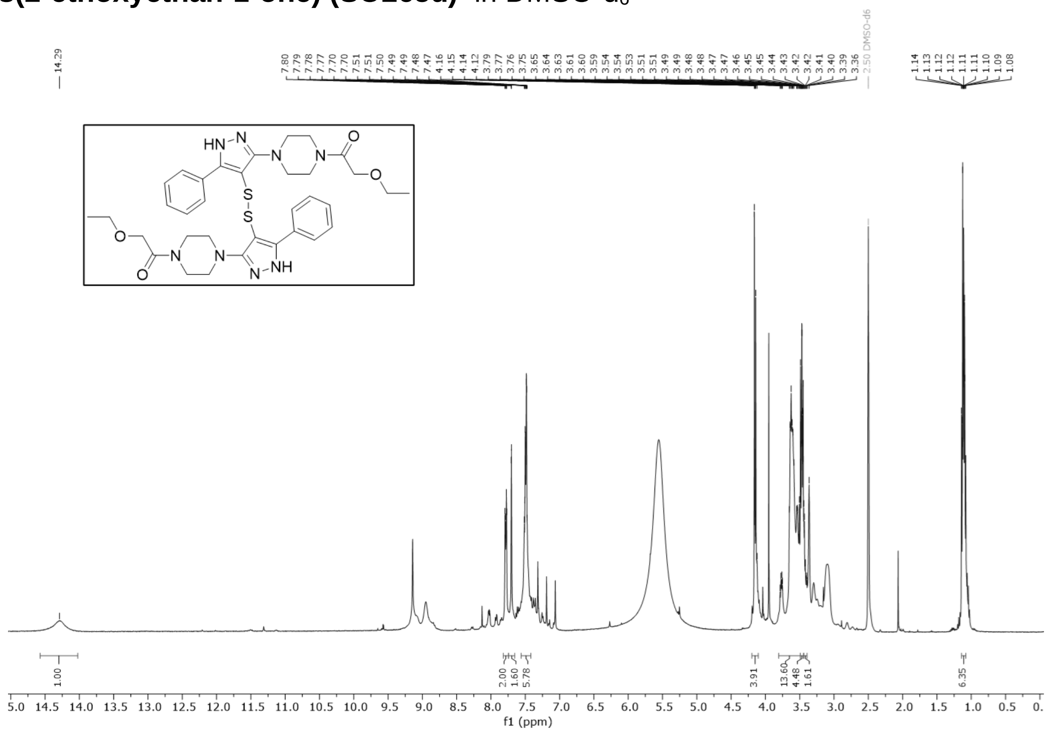


$^{13}$C NMR of **1,1'-((disulfanediylbis(5-phenyl-1*H*-pyrazole-4,3-diyl))bis(piperazine-4,1-diyl))bis(2-ethoxyethan-1-one) (SO265d)** in DMSO-$d_6$

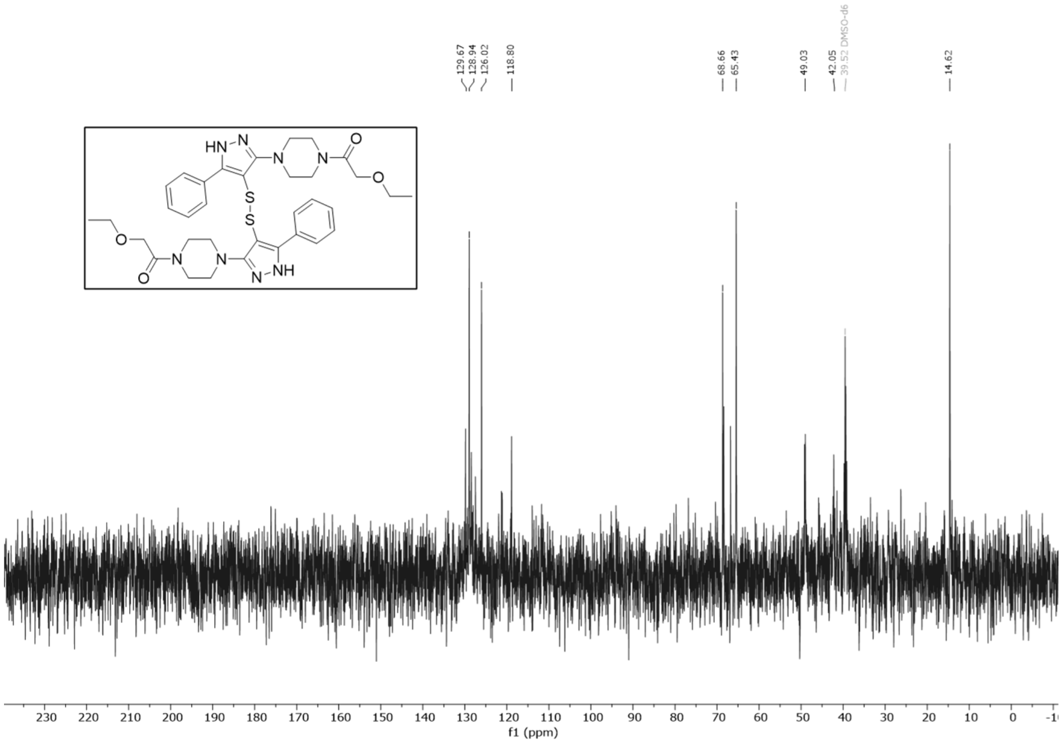


HMBC of **1,1'-((disulfanediylbis(5-phenyl-1*H*-pyrazole-4,3-diyl))bis(piperazine-4,1-diyl))bis(2-ethoxyethan-1-one) (SO265d)** in DMSO-$d_6$

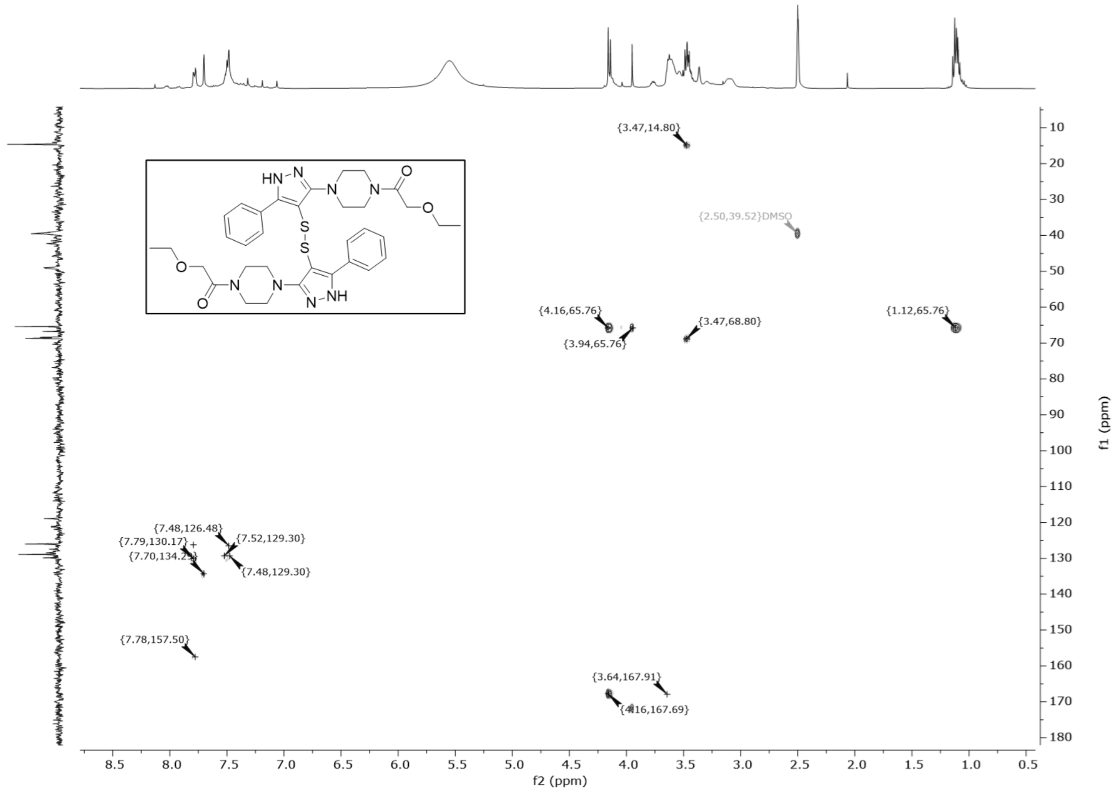

### (G) References


References

(1) Backus, K. M.; Correia, B. E.; Lum, K. M.; Forli, S.; Horning, B. D.; González-Páez, G. E.; Chatterjee, S.; Lanning, B. R.; Teijaro, J. R.; Olson, A. J.; et al. Proteome-wide covalent ligand discovery in native biological systems. *Nature* **2016**, *534* (7608), 570-574. DOI: 10.1038/nature18002.

(2) Sander, T.; Freyss, J.; von Korff, M.; Rufener, C. DataWarrior: an open-source program for chemistry aware data visualization and analysis. *J Chem Inf Model* **2015**, *55* (2), 460-473. DOI: 10.1021/ci500588j From NLM.

(3) Castellón, J. O.; Ofori, S.; Burton, N. R.; Julio, A. R.; Turmon, A. C.; Armenta, E.; Sandoval, C.; Boatner, L. M.; Takayoshi, E. E.; Faragalla, M.; et al. Chemoproteomics Identifies State-Dependent and Proteoform-Selective Caspase-2 Inhibitors. *Journal of the American Chemical Society* **2024**, *146* (22), 14972-14988. DOI: 10.1021/jacs.3c12240.

(4) Kedersha, N.; Panas, M. D.; Achorn, C. A.; Lyons, S.; Tisdale, S.; Hickman, T.; Thomas, M.; Lieberman, J.; McInerney, G. M.; Ivanov, P.; et al. G3BP–Caprin1–USP10 complexes mediate stress granule condensation and associate with 40S subunits. *Journal of Cell Biology* **2016**, *212* (7), 845-860. DOI: 10.1083/jcb.201508028.

(5) Shikwana, F.; Heydari, B.; Ofori, S.; Truong, C.; Turmon, A.; Darrouj, J.; Holoidovsky, L.; Gustafson, J.; Backus, K. CySP3-96 enables scalable, streamlined, and low-cost sample preparation for cysteine chemoproteomic applications. American Chemical Society (ACS): 2024.

(6) Yan, T.; Desai, H. S.; Boatner, L. M.; Yen, S. L.; Cao, J.; Palafox, M. F.; Jami-Alahmadi, Y.; Backus, K. M. SP3-FAIMS Chemoproteomics for High-Coverage Profiling of the Human Cysteinome**. *ChemBioChem* **2021**, *22* (10), 1841-1851. DOI: 10.1002/cbic.202000870.

(7) Desai, H. S.; Yan, T.; Yu, F.; Sun, A. W.; Villanueva, M.; Nesvizhskii, A. I.; Backus, K. M. SP3-Enabled Rapid and High Coverage Chemoproteomic Identification of Cell-State–Dependent Redox-Sensitive Cysteines. *Molecular & Cellular Proteomics* **2022**, *21* (4), 100218. DOI: 10.1016/j.mcpro.2022.100218.

(8) Hughes, C. S.; Moggridge, S.; Müller, T.; Sorensen, P. H.; Morin, G. B.; Krijgsveld, J. Single-pot, solid-phase-enhanced sample preparation for proteomics experiments. *Nature Protocols* **2019**, *14* (1), 68-85. DOI: 10.1038/s41596-018-0082-x.

(9) Hughes, C. S.; Foehr, S.; Garfield, D. A.; Furlong, E. E.; Steinmetz, L. M.; Krijgsveld, J. Ultrasensitive proteome analysis using paramagnetic bead technology. *Molecular Systems Biology* **2014**, *10* (10), 757. DOI: 10.15252/msb.20145625.
